# Pervasive aggregation and depletion of host and viral proteins in response to cysteine-reactive electrophilic compounds

**DOI:** 10.1101/2023.10.30.564067

**Authors:** Ashley R. Julio, Flowreen Shikwana, Cindy Truong, Nikolas R. Burton, Emil Dominguez, Alexandra C. Turmon, Jian Cao, Keriann Backus

## Abstract

Protein homeostasis is tightly regulated, with damaged or misfolded proteins quickly eliminated by the proteasome and autophagosome pathways. By co-opting these processes, targeted protein degradation technologies enable pharmacological manipulation of protein abundance. Recently, cysteine-reactive molecules have been added to the degrader toolbox, which offer the benefit of unlocking the therapeutic potential of ‘undruggable’ protein targets. The proteome-wide impact of these molecules remains to be fully understood and given the general reactivity of many classes of cysteine-reactive electrophiles, on- and off-target effects are likely. Using chemical proteomics, we identified a cysteine-reactive small molecule degrader of the SARS-CoV-2 non-structural protein 14 (nsp14), which effects degradation through direct modification of cysteines in both nsp14 and in host chaperones together with activation of global cell stress response pathways. We find that cysteine-reactive electrophiles increase global protein ubiquitylation, trigger proteasome activation, and result in widespread aggregation and depletion of host proteins, including components of the nuclear pore complex. Formation of stress granules was also found to be a remarkably ubiquitous cellular response to nearly all cysteine-reactive compounds and degraders. Collectively, our study sheds light on complexities of covalent target protein degradation and highlights untapped opportunities in manipulating and characterizing proteostasis processes via deciphering the cysteine-centric regulation of stress response pathways.

## INTRODUCTION

Protein homeostasis (proteostasis) is governed by an intricate interplay of spatiotemporally regulated cellular processes. Dysregulated protein abundance and quality control is central to nearly all human diseases, spanning cardiovascular disease^1,2^, cancer^3–5^, neurodegeneration^6–8^, and viral infections^9,10^. The latter is particularly noteworthy given both the recent COVID-19 pandemic and the preponderance of mechanisms by which host proteostasis processes are co-opted by pro-viral factors^11–15^. Consequently, the development of pharmacological agents that modulate protein homeostasis is a burgeoning area for both functional biology and drug development.

The established clinical impact of small molecule proteostasis modulators is showcased by a number of FDA-approved drugs, including proteasome inhibitors carfilzomib^16,17^ and bortezomib^18^, the estrogen receptor inhibitor fulvestrant^19,20^, and cereblon E3 ligase modulatory drug (CELMoD) agents^16,21–24^. In addition to these FDA approved proteostasis modulators, targeted protein degradation (TPD) technologies, including heterobifunctional molecules (two ligands linked together that recruit cellular degradation machinery to a protein of interest or POI) and molecular glues (one ligand that scaffolds a neo-interaction between the POI and ubiquitin ligase) continue to emerge at a breakneck pace. Their catalytic activity and capacity to target tough-to-drug classes of proteins distinguish degraders from other therapeutic modalities.

Hallmark examples of heterobifunctional technologies include PROteolysis Targeting Chimeras (PROTACs) ^25–28^, and the related opto-PROTACs^29^, PHOTACs^30^, and the degradation tag (dTAG) system^31^, all of which engage an E3 ubiquitin ligase to induce ubiquitylation and subsequent degradation of the POI. The utility of molecular glue degraders is exemplified by the discovery of CELMoD immunomodulatory drugs (IMiDs), including thalidomide, lenalidomide and pomalidomide, which have been used to treat multiple myeloma and autoimmune disease^21,22,32,33^. As shown by the substrate profiling of cereblon and VHL^34–41^, the two E3 ligases most widely commandeered by degrader molecules, favorable ternary complex formation is critical to efficient protein degradation^42–45^. As a result, some protein targets are more refractory to PROTAC/glue-mediated degradation, due in part to the absence of suitable E3 ligases. Thus, a number of new approaches including lysosome-targeting chimeras (LYTACs)^46^, autophagy-targeting chimeras (AUTACs)^47^, cytokine receptor-targeting chimeras (KineTACs)^48^, and hydrophobic tagging^49^ have further broadened the target scope amenable to pharmacological depletion.

While most small molecule degraders engage target proteins via a noncovalent mechanism, covalent PROTACS^50–58^ and covalent molecular glues^59–63^ are another promising class of degrader compounds. These compounds function by covalent modification of either an E3 ubiquitin ligase or a bait protein, primarily at cysteine residues, followed by ternary complex formation and subsequent ubiquitylation and proteasomal degradation of the POI. Recently, the covalent degrader paradigm has been reported to extend to E2 ubiquitin conjugating enzyme-recruiting molecular glues^64^ and E3 ligase adapter protein-targeting PROTACs^65^.

One key advantage of covalent degraders is the ready availability of a rich set of potential targets—as demonstrated by recent reports, including our own, the human proteome is rife with potentially druggable cysteine residues^66–76^ —each of these cysteines represents a possible labeling site for a covalent degrader molecule. The clinical success of several blockbuster electrophilic drugs, including afatinib, ibrutinib, and sotorasib^77–83^, further showcase the potential translation relevance of covalent degraders.

However, cysteine reactive compounds, even those found in drugs^84^, often react with a multitude of targets, which complicates mode of action studies. Exemplifying this challenge, several recent reports have demonstrated that electrophilic compound treatment, most notably with RA190^85^ and eeyarestatin^86,87^, afford widespread increased cellular ubiquitylation. While the mechanism driving this activity remains unclear, these findings hint at the likelihood of electrophile-induced global rewiring of cellular proteostasis mechanisms.

Here, we identify a class of covalent compounds that induce rapid depletion of the SARS– CoV-2 viral nonstructural protein 14 (nsp14). Multi-modal chemoproteomic analysis implicates covalent modification of both nsp14 and host protein disulfide isomerase (PDI) proteins in the observed rapid proteasome-mediated degradation of nsp14. Beyond nsp14, we observe global changes in cellular proteostasis in response to established and novel electrophilic degraders, including markedly elevated ubiquitylation, proteasome activation, and widespread depletion of host proteins, including most notably components of the nuclear pore complex (NPC). Seemingly synergistic with these activities, formation of stress granules (SGs), namely phase-separate organelles that form to promote cell survival during stress^88–90^, was observed to be a ubiquitous feature of cysteine-reactive drugs and drug-like molecules, rationalizing the observed pervasive aggregation and depletion of nucleoporins. Alongside stress granule formation, parallel electrophile-induced protein recruitment into aggresomes was also prevalent, including notably for nsp14. Taken together, our study connects cell stress processes to the mode-of-action of novel and established covalent degraders and provides a roadmap for delineating cysteine-specific modulation of proteostasis processes.

## RESULTS

### Chemoproteomic profiling of SARS-CoV-2 nonstructural proteins

Three years into the COVID19 pandemic, SARS-CoV-2 transmission continues at high frequency world-wide, warranting a need for new strategies to target viral proteins. Motivated by the success of the covalent cysteine-reactive drug Paxlovid^91^, we speculated that the SARS-CoV- 2 viral genome likely encodes additional ligandable cysteines, which could serve as sites for future drug development campaigns. Due to their functional significance and cysteine-rich sequences (**Table S1, Figure S1)**, we opted to subject cells overexpressing four individual non-structural proteins (nsps) (**Figure 1A**, nsp9^92–94^, nsp10^95–97^, nsp14^92,97–99^, and nsp16^96,100^) to cysteine chemoproteomic analysis using isotopic tandem orthogonal proteolysis- activity based protein profiling (isoTOP-ABPP)^67,68^ (**Figure 1B**) with a focused panel of electrophilic fragments (**Figure 1C, Scheme S1**).

**Figure 1.**
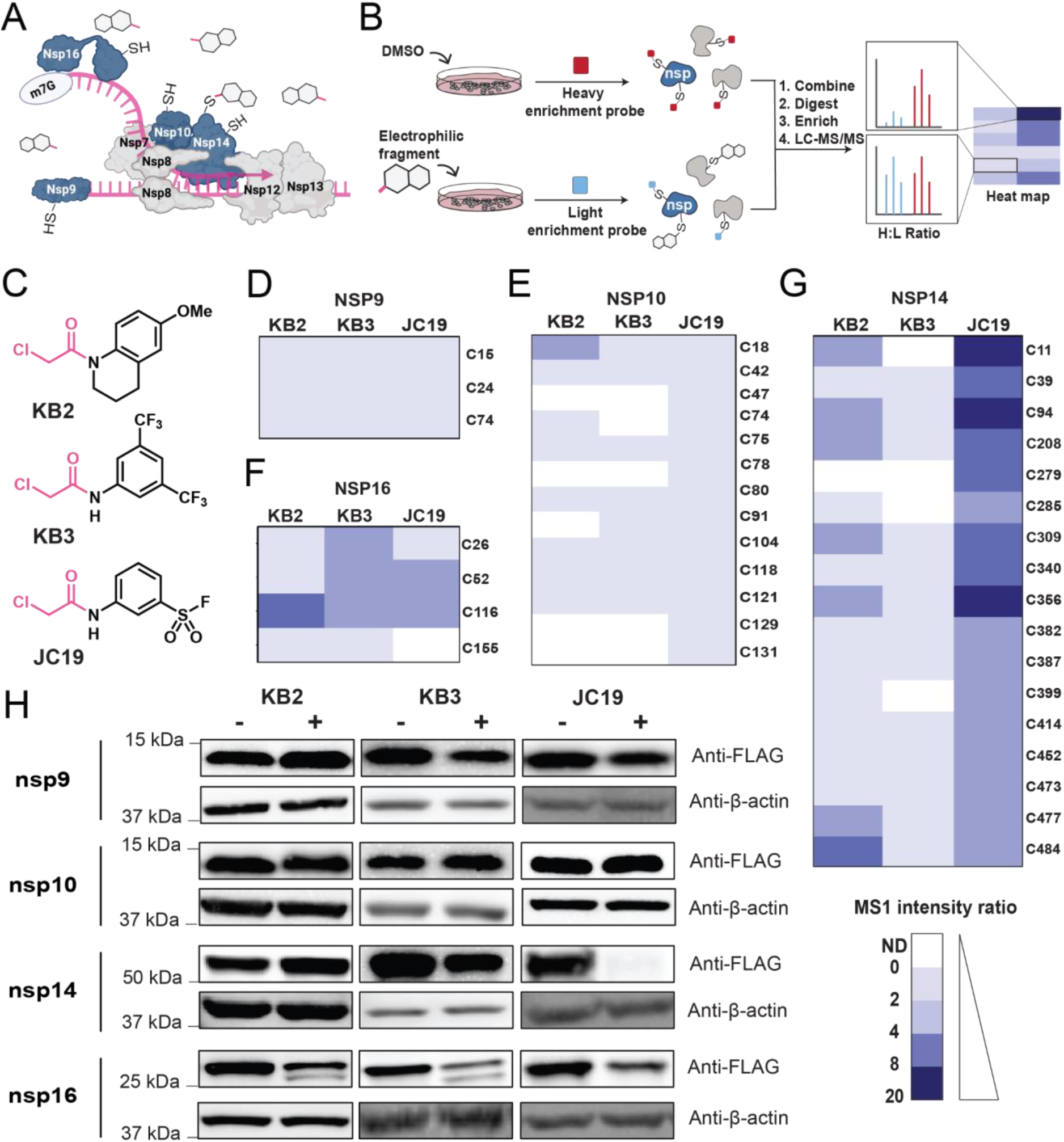
Chemoproteomic profiling of SARS-CoV-2 nonstructural proteins with electrophilic scout fragments identifies JC19 as a putative degrader of nsp14. (A) Schematic of SARS-CoV-2 nonstructural proteins (nsps). Blue nsps represent those that were screened against electrophilic fragments in this study. (B) Isotopic tandem orthogonal proteolysis- activity based protein profiling (isoTOP-ABPP) workflow used to identify ligandable cysteines within select nsps. (C) Chemical structures of the three electrophilic fragments used in isoTOP-ABPP experiments: KB2, KB3, and JC19. Electrophilic chloroacetamide moiety is colored pink. (D-G) Heatmaps representing the unlogged MS1 isoTOP-ABPP ratio for each identified cysteine in nsp9 (D), nsp10 (E), nsp16 (F), and nsp14 (G) based on the median value from multiple biological replicates of treatment with 100 µM KB2 (*n* ≥ 3), 100 µM KB3 (*n* ≥ 3), and 100 µM JC19 (*n* ≥ 3) for 1 hour. ND = not detected. (H) Immunoblot analysis of nsp abundance changes in response to treatment with each electrophilic fragment compound (KB2, KB3, or JC19) at 100 µM for 1 hour, relative to a DMSO- treated control. MS experiments shown in ‘D-G’ were conducted in ≥ 3 biological replicates in HEK293T cells. All MS data can be found in **Table S1.**

Promiscuously reactive scout molecules KB2 and KB3^67^ were selected to provide rapid insight into the ligandability of nsp cysteines. Bifunctional compound JC19^69^ was chosen to test the hypothesis that molecules incorporating sulfonyl fluorides in addition to cysteine-reactive chloroacetamide warheads might function as covalent molecular glues^101^, capable of inducing new protein complexes. Across all isoTOP-ABPP experiments, we detected 3, 13, 17, and 4 cysteines in nsp9, nsp10, nsp14, and nsp16, respectively, which corresponds to 100%, 100%, 74%, and 80% of the total cysteines in each of these proteins. While the majority of cysteines in each nsp were detected, our analysis revealed that nsp9 and nsp10 are generally insensitive to labeling by all three molecules tested, with only C18 of nsp10 showing a modest isoTOP-ABPP

MS1 H/L ratio (2.30), consistent with ∼50% occupancy at this residue (**Figure 1D,E**). By contrast, nsp16 showed elevated ratios across the majority of cysteines detected for KB3 and JC19 (**Figure 1F**). Nsp14, instead, showed elevated ratios that were specific to treatment with the bifunctional molecule JC19, again with heighted ratios observed for nearly all detected cysteines (**Figure 1G**). As prior studies, including our own, had revealed that generally only a small fraction of proteinaceous cysteines in a given protein sequence were likely to be liganded by an electrophilic molecule^67,70,73^, we speculated that the observed high ratios might instead stem from compound-induced depletion of nsp14 and nsp16, likely due to a protein degradation mechanism. To test this hypothesis, we assessed compound-dependent changes to each nsp abundance by immunoblot, which revealed a pronounced compound-induced decrease in detectable nsp16 and nsp14 (**Figure 1H**). Nsp16 abundance was moderately sensitive to all three scout molecules, whereas nsp14 depletion was specific to JC19 (**Figure 1H**).

### Both chloroacetamide and sulfonyl fluoride moieties are required for rapid depletion of SARS-CoV-2 nsp14

Intrigued by the marked compound-dependent depletion of nsp14 and nsp16 viral proteins, we opted to further characterize this effect. As the loss of nsp14 was both more scaffold-specific and more pronounced when compared to nsp16, we pursued focused structure-activity relationship (SAR) analysis with nsp14, with the goal of establishing a panel of active and inactive compounds required for mechanistic studies. To guide our SAR studies, we tested the time-, concentration- and cell-type generalizability of JC19-induced depletion in order to select optimal treatment conditions for library screening. We found that nsp14 depletion occurs rapidly, within 15 minutes of treatment, with near complete loss observed within 1 hour (**Figure 2A**). The depletion of nsp14 by JC19 was corroborated by immunocytochemical analysis of nsp14-expressing cells (**Figure S2A,B**). To confirm the generality of this effect, we extended this analysis to HeLa cells, which replicated the rapid nsp14 depletion (**Figure S2C**). JC19 was observed to be modestly potent, with an apparent degradation coefficient (DC50) of 8.7 µM with a 30 minute treatment time (**Figure 2B,C**).

**Figure 2.**
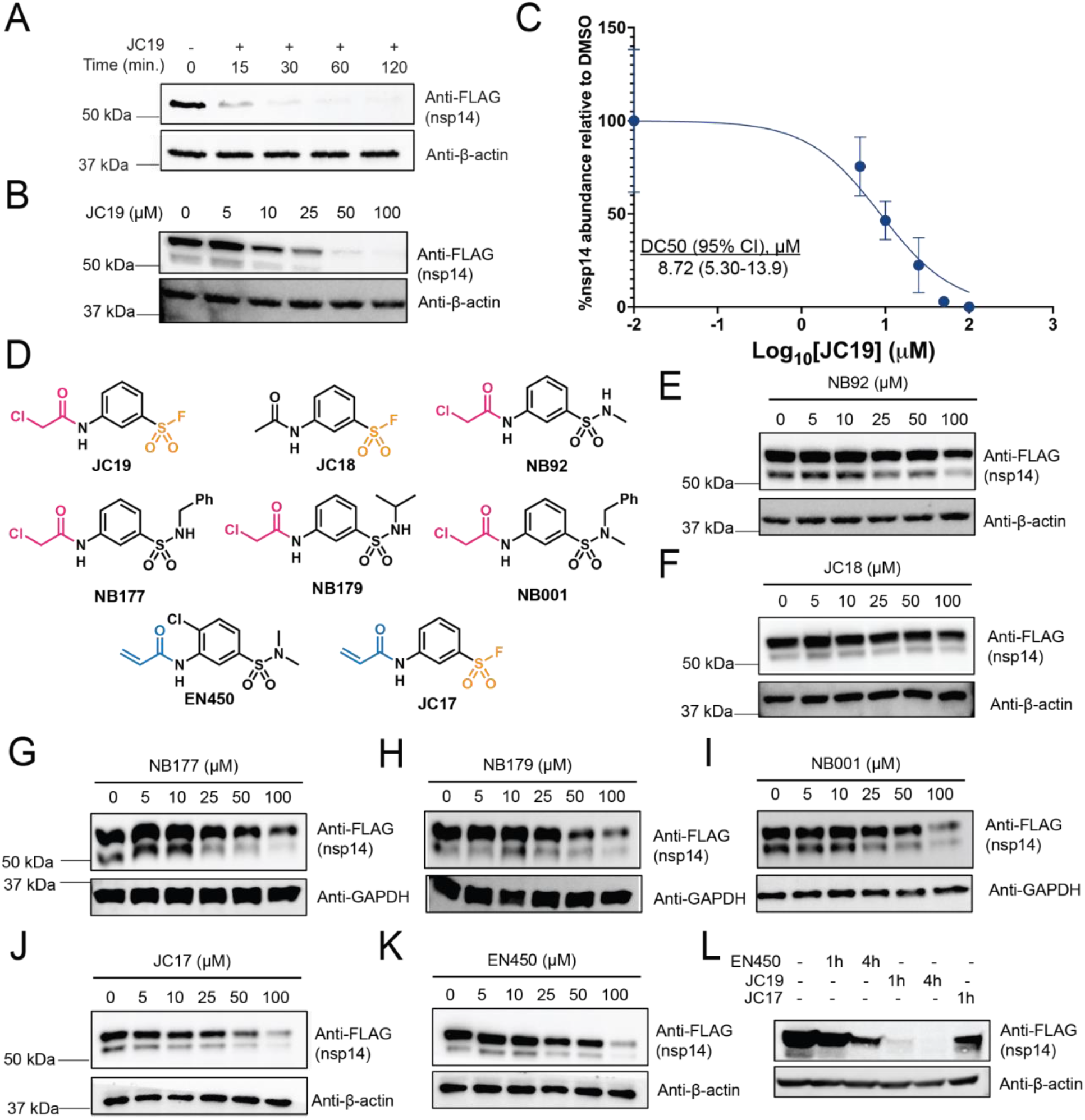
Both the chloroacetamide and sulfonyl fluoride electrophiles are required for the rapid depletion of nsp14. (A) HEK293T cells transiently expressing nsp14-FLAG were treated with 100 µM JC19 for the indicated times and nsp14 abundance assayed by immunoblot at each time point. (B) HEK293T cells transiently expressing nsp14-FLAG were treated with the indicated concentrations of JC19 for 30 minutes and the abundance of nsp14 was assayed by immunoblot at each concentration. (C) Replicate immunoblots (*n* = 3) were used to construct a degradation coefficient (DC50) curve with a calculated DC50 of 8.717 µM (95% confidence interval: 5.302-13.85 µM). (D) Chemical structures of compounds composing the focused library, including JC19 and JC19 analogues. Cysteine-reactive chloroacetamide warhead is colored pink, cysteine-reactive acrylamide warhead is colored blue, and sulfonyl fluoride warhead is colored orange. (E-K) HEK293T cells transiently expressing nsp14-FLAG were treated with the indicated concentrations of NB92 (E), JC18 (F), NB177 (G), NB179 (H), NB001 (I), JC17 (J), or EN450 (K) for 30 minutes, and nsp14 abundance assayed by immunoblot. (L) HEK293T cells transiently expressing nsp14- FLAG were treated with 100 µM JC19, EN450, or JC17 for the indicated times and nsp14 abundance assayed by immunoblot.

Guided by these findings, we next assembled a focused library of JC19-related analogues (**Figure 2D, Scheme S1**), including previously reported compounds EN450^64^, JC17, and JC18^69^, and newly synthesized molecules NB92, NB177, NB179, and NB001. Our main goals were to establish the portions of the molecule required for activity, to generate structurally matched inactive control probes, and to establish whether more potent lead compounds could be readily obtained. Included in this library were compounds JC18, containing only the sulfonyl fluoride electrophile, and compound NB92, containing only the chloroacetamide electrophile, which were targeted at testing whether both electrophiles were required for activity. NB177 (phenyl sulfonamide), NB179 (isopropyl sulfonamide), and NB001 (methyl phenyl sulfonamide) were selected to test whether the sulfonyl fluoride moiety could be harnessed as a point for late stage diversification, as has been reported previously^102^. We also included compound EN450, which was previously reported as a molecular glue and was observed to have notable structural resemblance to JC19^64^, and compound JC17 (an acrylamide analog of JC19), both of which were selected to test whether acrylamide-based compounds would afford comparable nsp14 loss. Given the more attenuated reactivity of the acrylamide group^67^, if active, these compounds could serve as starting points for more drug-like elaborated lead molecules.

The activity of each compound was tested by immunoblot, which revealed a markedly enhanced potency for JC19 above all other compounds. In contrast to JC19 (**Figure 2B**), NB92 afforded only slight nsp14 depletion at the highest concentration tested (**Figure 2E**), while JC18 afforded no nsp14 depletion at any concentration (**Figure 2F**), indicating that the dual electrophilic nature of JC19, particularly its cysteine reactivity, is crucial to its activity. Similarly, compounds NB177, NB179, and NB001, all of which lacked the sulfonyl fluoride group, showed markedly reduced activity relative to JC19 (**Figure 2G-I**). JC17 afforded a substantial, yet not complete, decrease in nsp14 abundance, supporting that the more reactive chloroacetamide of JC19 is important for enhanced activity (**Figure 2J**). Consistently, EN450, which features the less reactive acrylamide and lacks the sulfonyl fluoride, was significantly less active than JC19 (**Figure 2K**), but similar nsp14 depletion was observed with longer treatment times (**Figure 2L**). Collectively, although all JC19 analogues were significantly less potent, nsp14 depletion could still be observed at higher concentrations and longer treatment times, hinting at the likelihood of multiple parallel mechanisms of action.

### Nsp14 depletion is partially dependent upon the ubiquitin-proteasome system

Given the numerous recent reports of cysteine-reactive covalent degraders^50–60,64^, we next investigated whether JC19-induced nsp14 depletion was a result of protein degradation by cellular homeostasis machinery. To investigate the contribution of the ubiquitin-proteasome system (UPS), we pretreated cells with proteasome inhibitors MG132 or bortezomib, which each conferred partial protection against JC19-induced nsp14 degradation (**Figure 3A**). Analogous treatments using autophagy inhibitors revealed a similar partial rescue afforded by bafilomycin (**Figure S3**), which blocks lysosomal acidification by inhibiting V-ATPase^103^. However, no substantial protection was observed for other autophagy modulators, including ammonium chloride and hydroxychloroquine sulfate^104^ (**Figure S3**).

**Figure 3.**
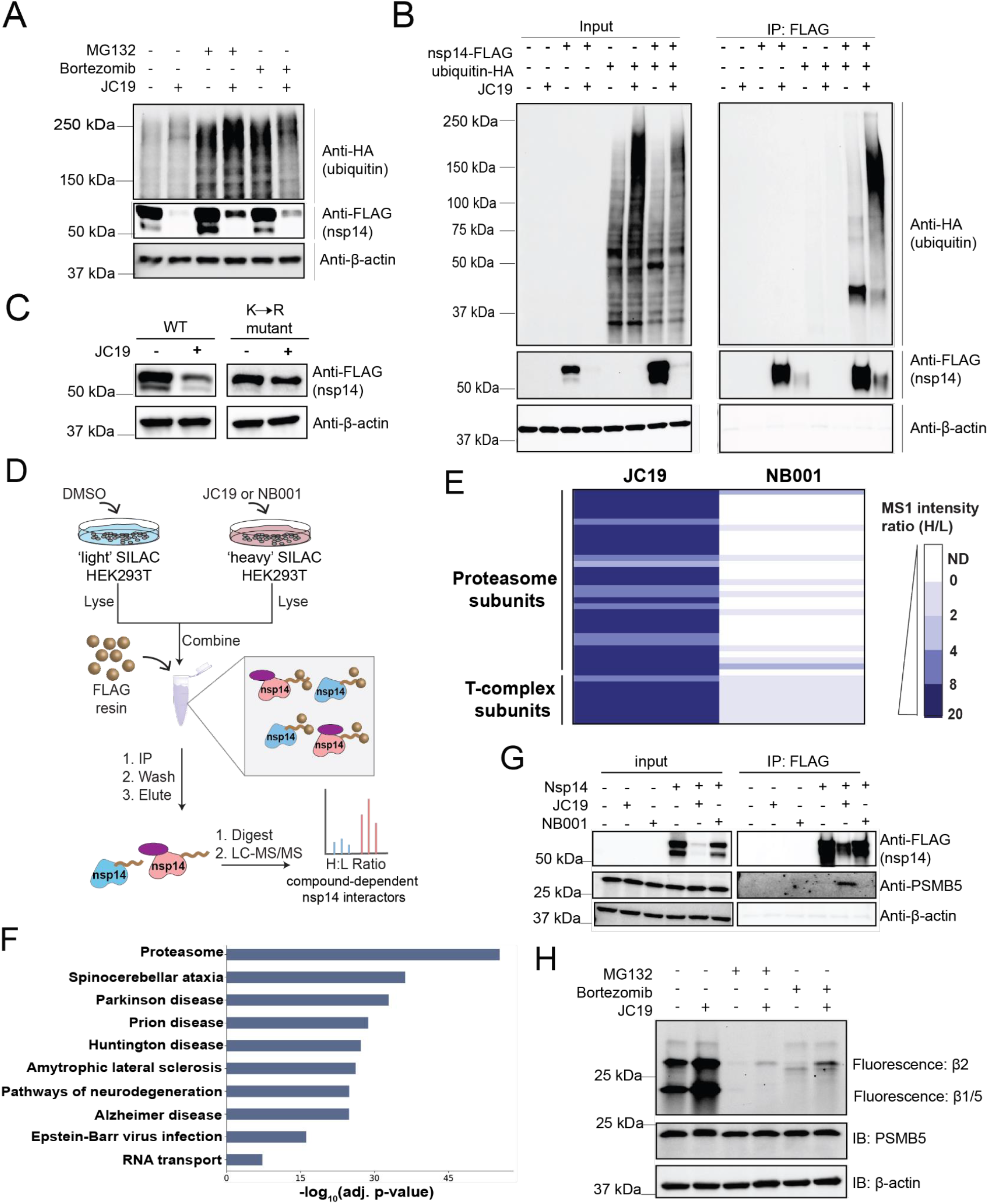
JC19-induced nsp14 degradation is mediated by ubiquitylation and the proteasome. (A) HEK293T cells transiently expressing nsp14-FLAG were pretreated with proteasome inhibitors MG132 (50 µM) and bortezomib (20 µM) for 14 hours, then treated with DMSO or 50 µM JC19 for 30 minutes, and immunoblot used to assay nsp14 abundance. (B) HEK293T cells transiently expressing nsp14-FLAG, ubiquitin-HA, neither, or both were treated with DMSO or 100 µM JC19 for 1 hour, immunoprecipitated on FLAG resin, and ran on immunoblot to assay ubiquitylation of nsp14. (C) HEK293T cells transiently expressing either wild-type (WT) or arginine mutant nsp14 (K13R, K34R, K288R, K318R, K339R, K423R, K433R, K440R, K469R) were treated with DMSO or 25 µM JC19 for 30 minutes and immunoblot used to assay nsp14 abundance. (D) Workflow for SILAC-based affinity purification mass spectrometry (AP-MS) of nsp14 in the presence and absence of JC19 or NB001. (E) Heatmap indicating mean unlogged MS1 intensity H/L ratio of proteasome and T-complex subunits immunoprecipitated with nsp14 in JC19 and NB001 AP-MS experiments (*n* = 3 for each compound). (F) KEGG pathway analysis of proteins that are immunoprecipitated with nsp14 in a JC19-dependent manner (mean SILAC log_2_ ratio > 1.5), using entire AP-MS dataset as background. KEGG 2021 Human database used for analysis. (G) Mock and nsp14- FLAG-expressing HEK293T cells were treated with 100 µM JC19, NB001, or DMSO for 1 hour, immunoprecipitated on FLAG resin, and immunoblot analysis used to depict immunoprecipitation of PSMB5. (H) HEK293T cells were pretreated with DMSO, 10 µM MG132, or 5 µM bortezomib for 3 hours, then treated with 100 µM JC19 for 1 hour. Cells were then treated with fluorescent proteasome activity probe Me4BodipyFL-Ahx3Leu3VS (500 nM, 1 hour) and in-gel fluorescence used to quantify proteasome activity. MS experiments shown in ‘E-F’ were conducted in 3 biological replicates in HEK293T cells. All MS data can be found in **Table S2.**

### JC19-induced nsp14 depletion is independent of canonical cell death pathways and electrophile sensing machinery

Puzzled by the only modest protection afforded by our panel of proteasome and autophagosomal inhibitors, we next expanded our mechanistic studies to interrogate whether JC19 was activating mechanisms involved in sensing electrophiles. Given the comparatively modest, low micromolar, DC50, we reasoned that electrophile-induced depletion of intracellular glutathione, well known to induce ferroptosis, could be involved in the mechanisms of depletion. Treatment with the widely utilized glutathione depleting reagent buthionine sulfoximine (BSO)^105^ did not afford appreciable nsp14 depletion (**Figure S4A**), and bulk cellular glutathione levels were insensitive to compound treatment (**Figure S4B**). Treatment with RSL3, an established GPX4 inhibitor and ferroptosis inducer^106–108^, failed to impact nsp14 abundance (**Figure S4C**). Beyond ferroptotic cell death, we also considered apoptotic cell death. PARP cleavage, which is a marker of programmed cell death, was not observed after a 1 hour treatment (**Figure S5A**) and membrane permeability was maintained during the treatment course for both active JC19 and relatively inactive EN450 (**Figure S5B**).

Looking beyond cell death processes, we investigated the involvement of the KEAP1- NRF2 system, which is central to cellular response to electrophiles^109–111^, and the unfolded protein response (UPR), the mechanism by which cells sense and eliminate misfolded proteins from the ER^112^. No NRF2 activation was observed in response to JC19 (**Figure S6A**), which indicated that during the comparatively short 1h treatment times, NRF2 transcriptional activity is likely not essential for JC19-induced nsp14 depletion. Similarly, we also observed that JC19 itself did not upregulate known UPR markers (**Figure S6B**), and nsp14 abundance was insensitive to treatment with established UPR activators, including thapsigargin, tunicamycin, and rapamycin (**Figure S6B**), further supporting a cysteine-dependent mechanism of depletion.

### JC19-induced nsp14 depletion is not due to lack of nsp10 expression or altered translation

As viral proteins, particularly when expressed in isolation, can be unstable, we next assessed the half-life of nsp14. Cycloheximide (CHX) treatment revealed a comparatively long (∼8h) protein half-life, with full nsp14 turnover not observed until 24 hours after translation inhibition (**Figure S7A**). Nsp14 complexes with nsp10 during viral infection, which is known to stabilize nsp14 and aid its enzymatic activity^113–116^. Therefore, we also assessed whether coexpression of nsp10 with nsp14 would preclude nsp14 depletion by JC19. Coexpression afforded enhanced nsp14 abundance, suggestive of increased protein stability stemming from complexation with nsp10, but this did not preclude nsp14 depletion. JC19 treatment of cells expressing both nsp10 and nsp14 revealed comparative insensitivity for nsp10 and marked nsp14 depletion (**Figure S7B**). These data support that JC19-induced nsp14 depletion is comparatively insensitive to nsp10 expression.

### JC19 induces ubiquitylation of nsp14 and nsp14 association with the proteasome

As the mechanism of nsp14 depletion was observed to be, at least partially, proteasome-dependent (**Figure 3A**), we revisited the contributions of proteasome-mediated degradation to nsp14 depletion. We found that JC19 treatment induced robust polyubiquitylation of nsp14 (**Figure 3B**), with no ubiquitylation observed upon treatment with comparatively inactive compound NB001 (**Figure S8**). Notably, pronounced compound-dependent global polyubiquitylation was also observed, indicating that the activity of JC19 might extend to other proteins beyond nsp14. To further assess nsp14 polyubiquitylation involvement in the mechanism of nsp14 depletion, we mutated nine potential lysine ubiquitylation sites to arginine residues; we included residues identified as ubiquitylated (GlyGly modified) in a JC19-dependent manner by LC-MS/MS (**Figure S9**, **Table S2**) as well as other solvent accessible lysines within nsp14 (PDB: 7QGI^99^). The mutated nsp14 construct was nearly completely insensitive to JC19 treatment (**Figure 3C**).

To capture proteostasis factors responsible for nsp14 polyubiquitylation, we next established an affinity purification-mass spectrometry (AP-MS) platform utilizing Stable Isotope Labeling by Amino Acids in Cell Culture (SILAC)^117^ (**Figure 3D**). Comparison of the interactomes of nsp14 versus GFP revealed 205 total nsp14-specific interactors (mean SILAC log_2_ ratio <1.5) including IMPDH2 and GLA, previously reported high-confidence nsp14 interactors^13^, providing evidence of the robustness of our approach (**Table S2**). In comparing nsp14 interactomes in the presence and absence of compound treatment, JC19 and NB001 afforded 133 and 21 total compound-induced interactions, respectively (mean SILAC H/L log_2_ ratio >1.5) (**Table S2**). Of the significant interactors, proteasome and T-complex subunits (components of a multiprotein chaperonin that aids in eukaryotic protein folding^118^) were significantly enriched in the JC19 dataset as compared to the less active NB001 treatment (**Figure 3E**), substantiating the role of the proteasome in JC19-mediated nsp14 depletion. KEGG pathway analysis of JC19-induced interactome revealed substantial enrichment for components of the proteasome together with proteins related to processes of protein aggregation (**Figure 3F**). Providing further evidence of a JC19-dependent nsp14-proteasome interaction, proteasome subunit beta type-5 (PSMB5) co-immunoprecipitated by nsp14 upon JC19, but not NB001, treatment (**Figure 3G**).

### Knockdown of individual E3 ubiquitin ligases is not sufficient to rescue nsp14 depletion

While prior reports have demonstrated that direct recruitment of substrate proteins to the proteasome subunit can bypass requirements for E3 ubiquitin ligase recruitment^119,120^, nearly all molecular glue and heterobifunctional degrader molecules reported to-date require engagement of ubiquitin ligase machinery. As AP-MS identified five E3 ubiquitin ligases that interacted with nsp14 in a JC19-dependent manner, HUWE1, UBR4, STUB1, TRIM21, and HECTD1, we next opted to interrogate the possible involvement of each E3 in nsp14 degradation. We performed siRNA knockdowns of each identified E3, followed by transient nsp14 overexpression and compound treatment. No change in nsp14 abundance nor sensitivity to compound treatment was observed upon knockdown of HUWE1, a known target of electrophilic compounds^66^ which has been recently implicated in clearance of protein aggregates^121^, nor STUB1, UBR4, or TRIM21 (**Figure S10**). HECTD1 knockdown afforded a modest depletion of nsp14, which we ascribed to the previously reported reduction in cellular proliferation upon HECTD1 knockout^122,123^ (**Figure S10E**).

### Cysteine-reactive electrophiles activate the proteasome

As we were surprised by the general insensitivity of nsp14 to depletion of individual E3 ubiquitin ligases, we chose to next explore two unusual aspects of this system, namely to revisit the incomplete MG132 rescue (**Figure 3A**) and the compound-induced increased global polyubiquitylation (**Figure 3B, S8**). This latter observation parallels that reported previously for several promiscuous cysteine-reactive compounds, including eeyarestatin^87^ and RA190^85^. Exemplifying the functional similarities between these compounds and JC19, both RA190 and eeyarestatin induce depletion of nsp14 (**Figure S11**). Given these marked global impacts on ubiquitylation, we hypothesized that electrophilic compounds might impact proteasome function directly. To test this hypothesis, we harnessed an established activity-based proteasome probe, Me4BodipyFL-Ahx3Leu3VS^124,125^. Consistent with complete proteasome inhibition, both MG132 and bortezomib treatment abolished detectable probe signal for the B1/5 and B2 bands, as assayed by in-gel fluorescence (**Figure 3H, S12**). For both JC19 and RA190 treatments, in the absence of proteasome inhibitor treatment, we observe a striking increase in Me4BodipyFL-Ahx3Leu3VS labeling, indicative of increased proteasome activity (**Figure 3H, S12**). The enhanced proteasome activity was independent of proteasome abundance (**Figure 3H, S12**). Upon pretreatment with either MG132 or bortezomib, followed by treatment with JC19 or RA190, increased proteasome activity was observed relative to treatment with either proteasome inhibitor alone (**Figure 3H, S12**). This observation implicates electrophilic compound treatment-induced residual proteasome activity as likely contributing to the only partial rescue from JC19-induced compound degradation afforded by proteasome inhibitors (**Figure 3A**). ABPP analysis of an extended panel of cysteine-reactive compounds revealed proteasome activation as a ubiquitous feature of all compounds analyzed, including both nsp14 active and inactive degraders (**Figure S12**), and previously reported NFKB degraders^64^ (**Figure S12**), suggesting that although proteasome activation likely aids in the JC19-mediated depletion of nsp14, it alone is not sufficient to induce the depletion.

### Covalent modification of nsp14 cysteines contributes to nsp14 depletion

Given the requirement for a cysteine-reactive electrophile to achieve robust nsp14 depletion, together with the highly cysteine-rich nature of nsp14, we postulated that direct covalent alkylation of nsp14 was required for protein depletion. We generated an alkynelated version of JC19 (JC19yne; **Scheme S1**) and subjected nsp14 overexpressing cells to a competition-enrichment experiment following the workflow shown in **Figure 4A**. We observe robust enrichment and competition of nsp14 **(Figure 4B, Table S3)** for cell-based labeling, with less substantial competition in the corresponding lysate-based experiment (**Figure S13A, Table S3**). These findings support direct covalent modification of nsp14, which occurs preferentially in cell-based labeling, and was further recapitulated by streptavidin-based affinity pulldown and immunoblot analysis (**Figure 4C**). Nsp14 was also preferentially enriched by JC19yne over JC19 when treatment was performed in cells (**Figure S13B, Table S3**), which mitigates the possible confounding effects of rapid nsp14 depletion during dual JC19 and JC19yne treatments. When compared to in situ labeling, lysate labeling with JC19yne afforded a more modest enrichment of nsp14 (**Figure S13C, Table S3**).

**Figure 4.**
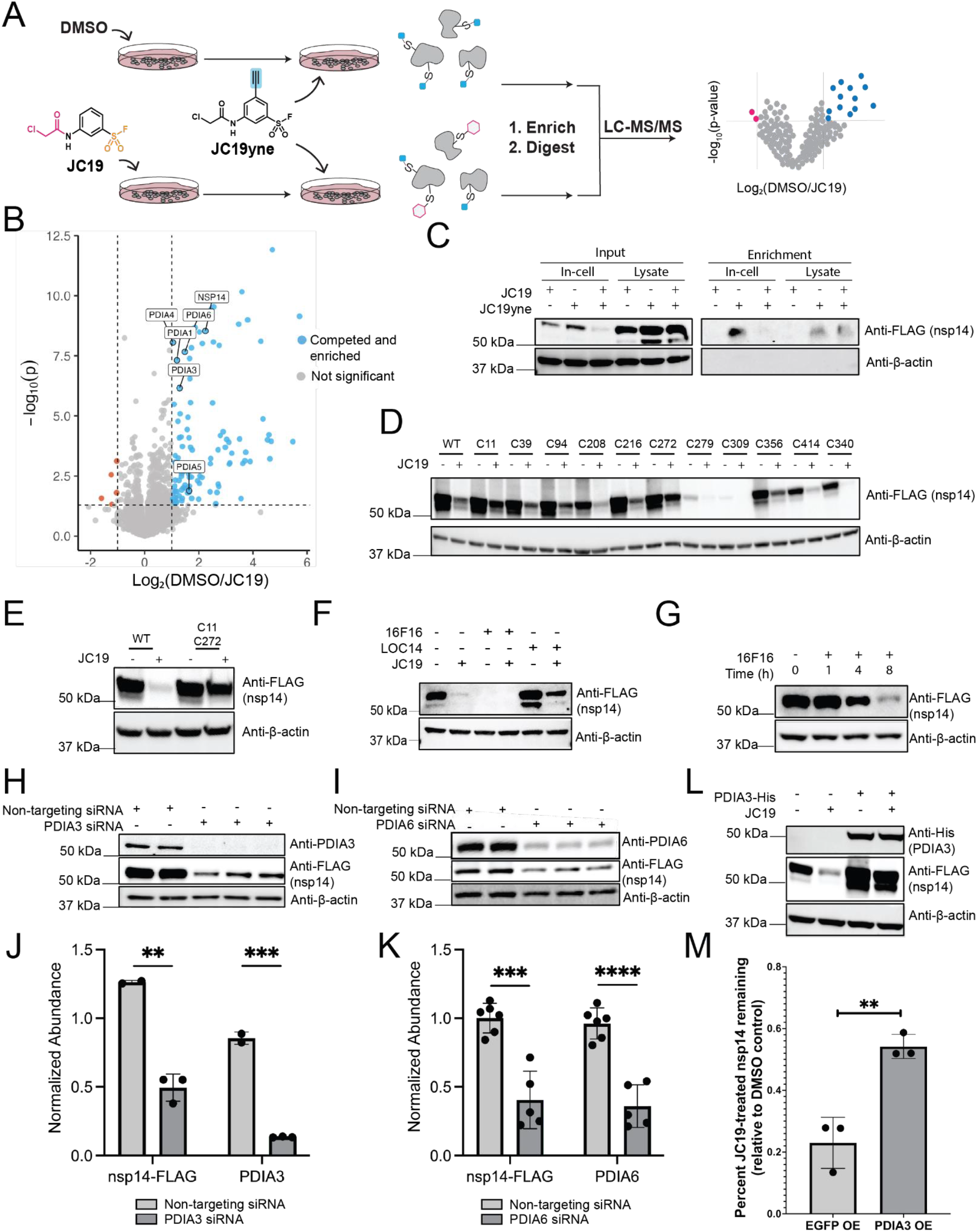
Direct labeling of nsp14 and the protein disulfide isomerase (PDI) family potentiates nsp14 degradation. (A) Workflow of JC19 competition experiment. HEK293T cells transiently expressing nsp14-FLAG were treated with either DMSO (*n* = 6) or 100µM JC19 (*n* = 6) for 20 minutes, and all replicates then treated with 100 µM JC19yne for 20 minutes. After cell lysis, proteins labeled by JC19yne were conjugated to biotin via copper-catalyzed azide-alkyne cycloaddition (CuAAC) and enriched on streptavidin resin. On- resin tryptic digest released peptides of labeled proteins for LC-MS/MS analysis to produce a volcano plot of significantly competed and enriched proteins. Label-free quantification was used to quantify protein abundance in each replicate. (B) Volcano plot depicting the results from the experiment outlined in (A), showing enrichment of nsp14 along with protein disulfide isomerases (PDIs). A Student’s t-test was performed between the DMSO (*n* = 6) and JC19 (*n* = 6) treated groups to calculate the p-values for the volcano plot. Significantly enriched/competed proteins were those with a log_2_ fold-change > 1 and p-value < 0.05. (C) Immunoblot analysis of nsp14 enrichment by JC19yne in the presence and absence of JC19 competition, for both cell and lysate treatments. Cells and lysates were treated as described in (A), but immunoblot analysis was used instead of LC-MS/MS. (D) HEK293T cells transiently expressing wild-type (WT) nsp14-FLAG or the indicated nsp14 cysteine mutants (all cysteine to alanine mutations, except Cys208, which is a cysteine to valine mutant) were treated with either DMSO or 50 µM JC19 for 1 hour and nsp14 abundance detected by immunoblot. (E) HEK293T cells transiently expressing WT nsp14-FLAG or the Cys11Ala/Cys272Ala double nsp14 mutant were treated with either DMSO or 50 µM JC19 for 1 hour and nsp14 abundance detected by immunoblot. (F) HEK293T cells transiently expressing nsp14-FLAG were treated with either DMSO, 10 µM 16F16, or 10 µM LOC14 for 14 hours, then treated with either DMSO or 50 µM JC19 for 1 hour, and immunoblot used to assay nsp14 abundance. (G) HEK293T cells transiently expressing nsp14-FLAG were treated with 10 µM 16F16 for the indicated times and nsp14 abundance assayed by immunoblot. (H) Immunoblot of the impact of siRNA-mediated knockdown of PDIA3 on nsp14- FLAG abundance (*n* = 3). (I) Immunoblot of the impact of siRNA-mediated knockdown of PDIA6 on nsp14- FLAG abundance (*n* = 5). (J) Quantification of nsp14 and PDIA3 abundance following siRNA knockdown of PDIA3 (*n* = 3). Statistical significance calculated with Student’s t-tests ** p<0.005, *** p<0.0005. (K) Quantification of nsp14 and PDIA6 abundance following siRNA knockdown of PDIA6 (*n* = 5). Statistical significance calculated with Student’s t-tests *** p<0.0005, **** p<0.0001. (L) HEK293T cells transiently coexpressing nsp14-FLAG and EGFP (as a control) or nsp14-FLAG and PDIA3-His were treated with either DMSO or 25 µM JC19 for 30 minutes and abundance of nsp14 assayed by immunoblot. (M) Quantification of the percent of nsp14-FLAG remaining after 25 µM JC19 treatment (30 minutes) of cells coexpressing nsp14-FLAG and EGFP (*n* = 3) or cells coexpressing nsp14-FLAG and PDIA3-His (*n* = 3), relative to a DMSO-treated control of the respective condition (*n* = 3). Statistical significance calculated with Student’s t-test ** p<0.005. MS experiments shown in (B) were conducted in 6 biological replicates in HEK293T cells. All MS data can be found in **Table S3.**

As these data support direct modification of nsp14 by JC19, we next sought to pinpoint the likely site(s) of covalent alkylation. We generated and evaluated a focused set of nsp14 cysteine mutants (Cys11Ala, Cys39Ala, Cys94Ala, Cys208Ala, Cys309Ala, Cys356Ala, Cys414Ala), which were selected based on elevated isoTOP-ABPP ratios (**Figure 1G**). No substantial protection from JC19 treatment was observed for any of the cysteine mutants, and the Cys424Ala and Cys208Ala mutants showed substantially reduced expression (**Figure S14**)—the latter cysteine has been implicated in the SARS-CoV orthologue nsp14 stability^97^. We therefore expanded our mutant set to include several additional cysteines including those that showed lower isoTOP-ABPP ratios (Cys279Ala and Cys340Ala), the Cys208Val mutant to increase residue hydrophobicity and thus stabilize the mutant protein, and several key zinc-finger proximal cysteines (Cys216Ala, Cys272Ala, Cys414Ala), with the hypothesis that covalent modification at these metal coordination sites could prove highly destabilizing. Notably, both Cys216 and Cys272 are found in non-proteotypic (>65 amino acid) tryptic peptides. Gratifyingly, with a reduced dose of JC19 (25 µM), we observed that the Cys11Ala and Cys272Ala mutants each afforded partial protection from compound-induced nsp14 depletion (**Figure 4D**). Providing further evidence of these cysteines as the likely labeling sites, the double mutant (Cys11Ala and Cys272Ala) was largely insensitive to JC19 treatment (**Figure 4E**).

### Covalent labeling of PDIA3 and PDIA6 contribute to nsp14 depletion

Given the comparatively modest micromolar DC50 observed for JC19 (**Figure 2C**), we reasoned that a number of additional host protein cysteines would be subject to covalent modification alongside the aforementioned nsp14 labeling sites. Therefore, we broadened our analysis of our JC19yne enrichment datasets to consider host proteins that showed both robust enrichment and competition by pre-treatment with JC19, in both cell-based labeling and lysate-labeling (**Figure 4B**, **S13**). Four members of the protein disulfide isomerase (PDI) family (PDIA1, PDIA3, PDIA4, and PDIA6) were observed to be markedly enriched by JC19yne in cells and lysates (**Figure S13B,C, Table S3**), with this enrichment competed by pre-treatment with JC19 in cells and lysates (**Figure 4B, S13A, Table S3**). To identify the PDI cysteines labeled by JC19 and related analogues, we expanded our isoTOP-ABPP analysis to include proteome-wide ligandability landscapes for JC19, KB2, NB92, NB177, NB179, NB001, JC17, EN450, KB7, and, JC36. In aggregate, our isoTOP-ABPP analysis of compound-treated HEK293T cells identified 14665 total unique cysteines from 4839 proteins, including 20 PDI cysteines (**Table S4**), multiple of which were labeled by JC19 (**Figure S15**). Notably, no widespread PDI labeling was observed for EN450, JC17, and NB179, which had all exhibited substantially attenuated nsp14 depletion activity (**Figure 2H,J-L**), particularly at shorter analysis timepoints. As the PDI family are ER-resident oxidoreductases that play essential roles in protein folding^126,127^, with covalent PDI inhibitors implicated in inducing activation of the Protein Response (UPR)-associated transcription factor ATF6^128,129^, we postulated that JC19 nsp14 depletion activity was potentiated by covalent PDI engagement.

To test this hypothesis, we pursued parallel chemical (**Figure 4F,G**) and genetic approaches (**Figure 4H-M**). We find that treatment with the previously reported irreversible covalent PDI inhibitor 16F16^130^ afforded nsp14 depletion (**Figure 4F**), albeit requiring substantially longer treatment times when compared with JC19 (**Figure 4G**). Intriguingly, pre-treatment with the reversible covalent PDI inhibitor LOC14^131^, which, in vitro, has been reported to stabilize PDIA1 in an inactive conformation afforded a statistically significant partial protection from JC19 treatment (**Figure 4F, S16**). The seemingly contradictory activities of these two PDI inhibitors prompted us to conduct siRNA knockdowns of each identified PDI, with the expectation that only a subset of these highly homologous proteins would impact nsp14 abundance. Consistent with this model, we find that siRNA depletion of PDIA3 or PDIA6 significantly decreased nsp14 abundance, as detected by immunoblot and by quantitative LC-MS/MS analysis (**Figure 4H-K**). In contrast to PDIA6 and PDIA3 gene expression knockdowns, nsp14 was insensitive to PDIA1 and PDIA4 gene expression knockdown (**Figure S17A-D**). Bulk proteomic analysis substantiated these findings revealing significant nsp14 depletion afforded by PDIA6 knockdown (**Figure S17E,F, Table S3**). In total 103 proteins were sensitive to PDIA6 knockdown in nsp14 overexpressing cells (**Figure S17E**) with 59 proteins also depleted by PDIA6 knockdown in cells lacking nsp14 expression (**Figure S17F**), supporting a comparatively specific mode of action. Further implicating PDI activity, we find that heterologous overexpression of PDIA3 significantly decreased nsp14 sensitivity to JC19 treatment (**Figure 4L,M**). Collectively, our data support a mechanism by which rapid nsp14 depletion is achieved by destabilization of nsp14 via both direct labeling of nsp14 cysteines and through PDI inhibition.

### Multimodal proteomics implicates global rewiring of the proteome by electrophilic compounds

Inspired by the handful of proteins found to be sensitive to PDI depletion (**Figure S17E,F**), we broadened our analysis to assess the impact of cysteine-reactive electrophiles on the host proteome. Building upon our findings for nsp14 (**Figure 1G**) and those of prior studies^70^, we harnessed our rich isoTOP-ABPP datasets (**Table S1**) to pinpoint compound-induced changes to protein abundance, inferred from identification of multiple high-ratio cysteines belonging to individual proteins. For the JC19, EN450, KB2, and JC36 datasets, ∼15-25% of all proteins for which more than one cysteine was identified were found to harbor multiple high-ratio cysteines (log_2_(H/L) > 1), termed here multiple high-ratio cysteine proteins (**Figure 5A**). Given the outsized impact of this subset of compounds on the proteome (**Figure S18**), we postulated that additional insights into the mechanism of nsp14 depletion might be gleaned from assessing whether specific fractions of the proteome were disproportionately impacted by compound treatment. Therefore, we next stratified our isoTOP-ABPP datasets by UniProtKB cellular component annotations. While multiple high-ratio cysteine proteins were present in all cellular components (**Figure S18**), we observed a marked increase in the frequency of high-ratio cysteines for protein subunits of the nuclear pore complex (NPC), when compared to the rest of the proteome (**Figure 5B**)—the NPC is a large (∼100 mDa) complex made up of >30 unique proteins that regulate biomolecule transport between the nucleus and cytosol^132^. Across all compounds, 18 nucleoporins were multiple high-ratio cysteine proteins.

**Figure 5.**
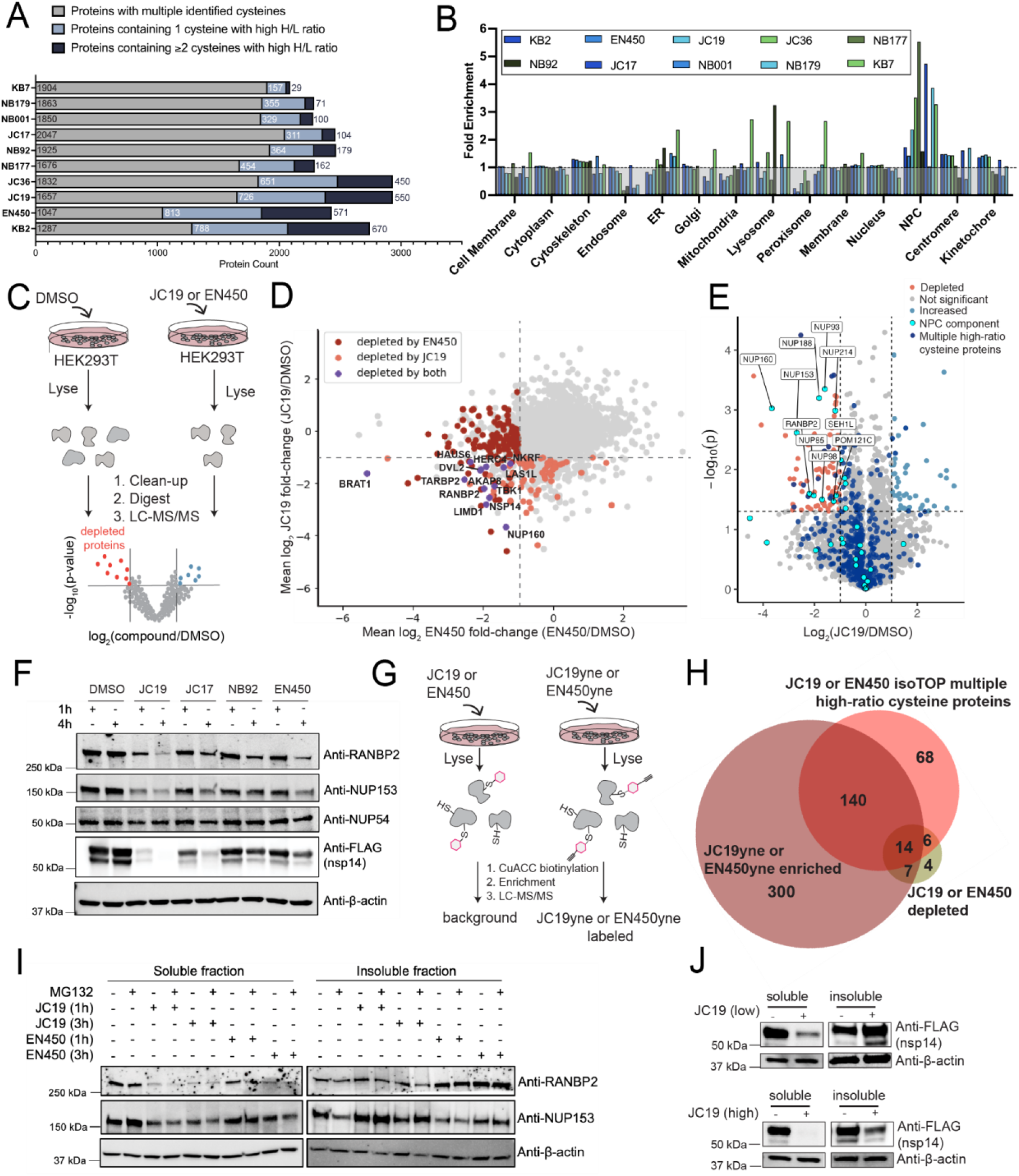
Multimodal proteomics implicates global rewiring of the proteome by electrophilic compounds. (A) Bar plot depicting the number of proteins in indicated isoTOP-ABPP experiments with multiple (> 1) identified cysteines (gray bars), 1 cysteine with a log_2_ H/L ratio > 1 (light blue bars), and multiple (≥ 2) cysteines with a log_2_ H/L ratio > 1 (navy blue bars) (*n* ≥ 2 for all experiments). (B) Cellular compartment (Uniprot annotated) enrichment analysis of proteins with multiple (≥ 2) cysteines with a log_2_ H/L ratio > 1 in the indicated isoTOP-ABPP experiments, as compared to all proteins with multiple identified cysteines. Fold change enrichment calculated by dividing the frequency in multiple high-ratio cysteine proteins by frequency in all identified multi-cysteine proteins. (C) Workflow for label-free quantification (LFQ) global proteomics in response to compound treatment. (D) Scatter plot depicting all proteins identified in both JC19 (100 µM, 1 hour, *n* = 3 for both JC19 and DMSO) and EN450 (100 µM, 4 hours, *n* = 3 for both EN450 and DMSO) bulk proteome experiments with light red colored dots corresponding to proteins significantly depleted by JC19 and dark red colored dots corresponding to proteins significantly depleted by EN450. Proteins with a log_2_(fold-change) < -1 and p-value < 0.05 are considered significantly depleted. Proteins depleted by both compounds are colored purple and labeled. (E) Volcano plot of JC19 bulk proteome analysis. HEK293T cells transiently expressing nsp14-FLAG were treated with DMSO (*n* = 3) or 100 µM JC19 (*n* = 3) for 1 hour, then prepared for LC-MS/MS analysis via SP3 clean-up. Label free quantification was used to measure protein abundance in each replicate, and the fold-change of the abundance for each protein plotted against its p-value. Proteins with a log_2_(fold-change) < -1 and p-value < 0.05 are considered significantly depleted. Proteins that were identified as having multiple high-ratio cysteines in the JC19 isoTOP-ABPP are colored navy blue. Nucleoporin proteins are colored cyan. Significantly depleted nucleoporins are labeled. (F) HEK293T cells transiently expressing nsp14-FLAG were treated with DMSO or 100 µM JC19, JC17, NB92, or EN450 for 1 or 4 hours and immunoblot used to assay changes in abundance for nsp14-FLAG, RANBP2, NUP153, and NUP54. (G) Workflow for JC19yne and EN450yne enrichment experiments. (H) Overlap of proteins identified as statistically depleted by JC19 or EN450 in LFQ bulk proteome datasets (*n* = 3 for each compound), proteins identified as statistically enriched by JC19yne or EN450yne (*n* = 6 for JC19yne and *n* = 4 for EN450yne), and proteins identified as having multiple (≥ 2) cysteines with elevated ratios for JC19 or EN450 by isoTOP-ABPP (*n* ≥ 3 for each compound). Proteins included in the Venn diagram include only those identified in all representative datasets. (I) Immunoblot analysis of RANBP2 and NUP153 abundance within the soluble and insoluble fractions in response to 100 µM JC19 or EN450 treatment at the indicated time points, both with and without a MG132 pretreatment (25 µM for 5 hours). (J) HEK293T cells transiently expressing nsp14-FLAG were treated with DMSO, a low dose of JC19 (25 µM for 30 minutes), or a high dose of JC19 (100 µM for 1 hour) and subject to immunoblot analysis. Cells were lysed in 0.3% CHAPS in PBS to generate the soluble lysate, and after clearance by centrifugation the insoluble debris was solubilized in 8 M urea in PBS to generate “insoluble’ lysate. All MS data can be found in **Table S4.**

Although the presence of multiple high-ratio cysteines in a single protein is suggestive of an abundance change, a subset of these cysteines could instead, or additionally, represent bona fide covalent alkylation sites. Therefore, we performed bulk proteomic analysis following the workflow shown in **Figure 5C** to assess compound-induced changes to protein abundance proteome-wide. We opted to compare chloroacetamide JC19 and acrylamide EN450, given their shared impact on the NPC together with JC19’s unique PDI labeling activity. In aggregate, JC19 and EN450 treatment afforded decreased abundance for 245 total proteins (log_2_(fold-change) < - 1 and p-value < 0.05 relative to DMSO). Suggestive of non-redundant modes of protein depletion, only 13 proteins were significantly sensitive to both compounds (**Figure 5D**). Gratifyingly, and consistent with our immunoblot analysis (**Figure 2L**), nsp14 was found to be sensitive to both JC19 (1 hour treatment) and EN450 (4 hour treatment) (**Figure S19A,B)**. Curiously, we observed that NFKB1 was not depleted by JC19 or EN450 (**Figure S19A,B**), in contrast with the prior report using this compound^64^. Immunoblot analysis further substantiated that NFKB1 abundance was insensitive to compound treatment (**Figure S19C**) both in the presence and absence of nsp14. As comparable compound concentrations were used in both studies, we ascribe this difference in depleted targets to our shorter treatment conditions (1–4 hours) compared with 24 hours used previously^64^. We did, however, observe that the NFKB regulator, NKRF^133–135^, was highly sensitive to both JC19 and EN450 (**Figure S19A,B**).

Consistent with our analysis of the multiple high-ratio cysteine protein isoTOP-ABPP subset (**Figure 5B**), we observed that the compound-depleted protein subsets from both JC19 and EN450 bulk proteomic datasets were also enriched for components of the nuclear pore (**Figure 5E, S20, S21**). RANBP2 and NUP160 were depleted by both JC19 and EN450. When compared to EN450, JC19 treatment was observed to more substantially impact the NPC, affording depletion of 10 nucleoporins out of 32 identified. Four of the 33 identified nucleoporins were sensitive to EN450, including AAAS and NUP133 (**Figure S20**), which were not significantly sensitive to JC19. The electrophile-sensitive nucleoporins were observed to be located throughout the NPC (**Figure S22**). Immunoblot analysis substantiated these findings and showed that the effect extended to additional cysteine-reactive electrophiles, with RanBP2 (Nup358), Nup153, and to a lesser extent, Nup54 all sensitive to JC19, JC17, NB92, and EN450 (**Figure 5F, S23**). Nsp14 overexpression was observed to moderately potentiate compound-induced depletion of RANBP2 and NUP205. In contrast, NUP153, NUP54, NUP133, and RAE1 depletion was unaffected by nsp14 expression (**Figure S24**).

In order to understand whether the NPC depletion was related to nsp14 depletion, or an independent consequence of electrophile treatment, we opted to further characterize this observation. As we had found that direct modification of nsp14 cysteines was involved in protein depletion (**Figure 4E**), we postulated that cysteine modifications could be a more generalized feature of depletion-sensitive proteins. To test this model, we first returned to our JC19 and EN450 isoTOP-ABPP datasets. We find that most nucleoporins sensitive to compound depletion had at least one high isoTOP-ABPP ratio cysteine, with 2 of 4 and 8 of 10 nucleoporins falling into the multiple high-ratio cysteine subsets for EN450 and JC19, respectively. More broadly, of the entire 245 depleted protein set, 120 were found to harbor at least one high ratio cysteine. To further delineate the contributions of direct versus indirect labeling, we compared the protein targets statistically enriched by clickable EN450yne or JC19yne (**Figure 5G**) with the proteins statistically depleted by the corresponding parent compounds (**Figure 5C**), and proteins containing multiple high-ratio cysteines in the JC19 or EN450 isoTOP-ABPP. While a substantial overlap between these groups was observed, a number of proteins that are significantly depleted by electrophilic fragments are not enriched by either alkyne analogue (**Figure 5H**), suggesting direct labeling of the target is not always required for protein depletion. Exemplifying this process, while nsp14 was robustly enriched by JC19yne (**Figure 4B, S13B**) and depleted by JC19 (**Figure S19A**), EN450yne failed to capture nsp14 (**Figure S25B**), despite also affording marked protein depletion (**Figure S19B**). Similarly, SEC13 was the only nucleoporin significantly enriched by JC19yne (**Figure S25A**), despite the dramatic impact of JC19 on nucleoporin abundance. None of the five nucleoporins enriched by EN450yne were significantly depleted in the bulk dataset (**Figure S25B**), which provides evidence that direct modification is not required for depletion of most nucleoporins. Taken together, these data hit at the likelihood of multiple mechanisms of depletion occurring in response to electrophilic compound treatment, with direct alkylation of proteins synergizing with indirect activity.

To further tease apart the shared and unique features of the nucleoporin and nsp14 depletion mechanisms, we revisited the involvement of the aforementioned E3 ubiquitin ligases and PDIs. We find that generally the majority of nucleoporins are insensitive to PDI depletion (**Figure S26, S17E-F, Table S3**). While HUWE1, UBR4, STUB1, and TRIM21 knockdowns did not impart a substantial protective effect against nucleoporin depletion (**Figure S27A-D**), HECTD1 knockdown revealed protection against JC19-mediated depletion of NUP153 (**Figure S27E**). However, overexpression of HECTD1 afforded no notable protection of NUP153 from JC19-induced depletion (**Figure S27F**), suggestive of the activity of HECTD1 depletion as linked to its broader effects on cell cycle progression^122^. Similarly, proteasome inhibition also failed to confer protection from JC19- and EN450-induced depletion of RANBP2 or NUP153 (**Figure 5I**).

Guided by the seemingly proteasome-independent mechanism of depletion and the prior reports of nucleoporin aggregation in the context of Mallory-Denk bodies^136^, we broadened our analysis to include assessment of compound-induced protein aggregation. Consistent with protein aggregation, we observed marked compound-induced accumulation of RANBP2 and NUP153 in the detergent insoluble cellular fraction by immunoblot (**Figure 5I**). Bulk proteomic analysis also revealed a marked increase in frequency of NPC subunits in the insoluble fraction upon compound treatment, relative to control-treated samples (**Figure S28A**). These data implicate compound-dependent nucleoporin aggregation as central to the nucleoporin depletion mechanism. Looking more broadly, analysis of both the JC19 and EN450 bulk proteomic datasets revealed substantial overlap in protein identifications between soluble and insoluble datasets (**Figure S28B, C**). Of the proteins identified in both soluble and insoluble datasets for each compound, a handful of proteins were shown to be significantly depleted in both soluble and insoluble (bone fide degraded), while a greater number of proteins were significantly depleted from the soluble and significantly accumulated in the insoluble (aggregated) (**Figure S28D, E**). The widespread aggregation of proteins in response to cysteine-reactive electrophiles inspired us to ask whether the aggregation mechanism extended to nsp14. Immunoblot analysis revealed accumulation of nsp14 in the insoluble fraction upon treatment with lower doses of JC19, suggestive of aggregation, whereas treatment with higher doses of JC19 induced depletion of nsp14 in both the soluble and insoluble fractions (**Figure 5J**), suggesting bone fide degradation. Extension of this analysis to nsp16, which also displayed compound-dependent abundance changes (**Figure 1F,H**), revealed that although there was no accumulation in the insoluble fraction at lower doses, higher doses of JC19 similarly induced depletion from both fractions (**Figure S29**). Collectively, our data suggest that treatment with cysteine-reactive electrophiles induces aggregation of both host and viral proteins, and that this aggregation can potentiate protein degradation.

### Cysteine-reactive small molecules induce formation of stress granules

We next considered the possible involvement of general cell stress response mechanisms. We were by the many features of our system suggestive of cellular stress and dysregulated proteostasis, including (1) the accumulation of proteins into the detergent insoluble fraction, which was suggestive of the formation of protein aggregates; (2) enrichment of aggregation-related processes in our AP-MS dataset (**Figure 3F**); (3) compound-induced increases in global polyubiquitylation (**Figure 2B**); (4) increased proteasome activity (**Figure 2H** and **Figure S12**); (5) involvement of protein folding machinery (**Figure 4H-K**); (6) and rapid kinetics (**Figure 2A**).

One widespread response to numerous cellular stressors (e.g. osmotic stress, oxidative stress, heat shock, nutrient deprivation, viral infection) is the formation of stress granules (SGs), namely, dynamic, phase-separated, membraneless organelles that sequester RNA and RNA-binding proteins within the cytoplasm to promote cell survival during stress^89,90,137,138^. Intriguingly, a number of NPC nucleoporins have been reported to accumulate into SGs in response to cell stress^8,139^.

Inspired by the presence of NPCs in SGs and the shared cysteine reactivity of electrophilic small molecules and stress-granule-inducing oxidative stressors (e.g. H_2_O_2_ and sodium arsenite), we next asked whether JC19 or EN450 could induce stress granules—we hypothesized that stress-induced protein aggregation was contributing to the depletion of nsp14, nsp16, and the compound-sensitive NPC subunits. We generated a U2OS cell line that stably expresses doxycycline-inducible V5-tagged G3BP1, a core component and well-known nucleator of stress granules^138,140^. Treatment of our reporter cells with sodium arsenite afforded robust formation of characteristic G3BP puncta, consistent with SG formation (**Figure 6A**). Similar puncta were also observed in cells treated with JC19, EN450, or NB001, indicating robust SG-formation in response to cysteine-reactive electrophilic fragments. Formation of SGs was observed at modest compound doses (25-50 µM) (**Figure S30**) that have been used, in the case of EN450, for molecular glue degrader analyses^64^.

**Fig 6.**
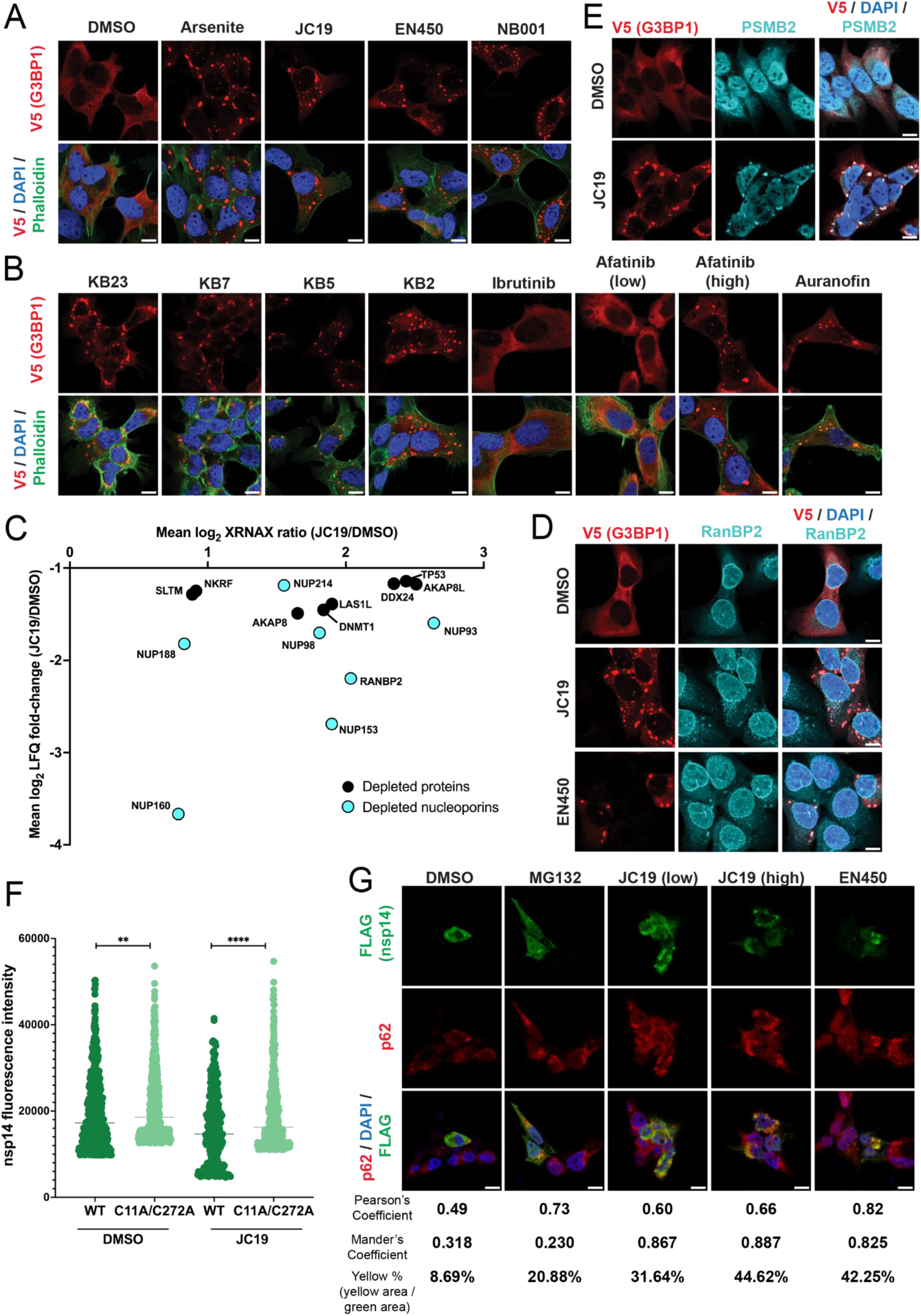
Cysteine-reactive electrophilic fragments induce global remodeling of the proteome through protein aggregation and stress granule formation. (A) Immunofluorescence microscopy of G3BP1- containing stress granules (SGs) in response to DMSO (vehicle), sodium arsenite (400 µM, 1 hour), JC19 (100 µM, 1 hour), EN450 (100 µM, for 1 hour), and NB001 (100 µM, 1 hour). U2OS cells were imaged on an LSM880 confocal microscope at 63X objective with 2X manual zoom. Scale bar = 10 µm. (B) Immunofluorescence microscopy of G3BP1-containing stress granules (SGs) in response to KB23 (100 µM, 1 hour), KB7 (100 µM, 1 hour), KB5 (100 µM, 1 hour), KB2 (100 µM, 1 hour), ibrutinib (10 µM, 14 hours), low afatinib (100 nM for 14 hours), high afatinib (10 µM for 1 hour), and auranofin (10 µM for 1 hour). Images were acquired on an LSM880 confocal microscope at 63X objective and 2X manual zoom. All scale bars = 10 µm. (C) Scatter plot of the 15 proteins shared between the JC19 XRNAX dataset and the proteins significantly depleted by JC19 in the LFQ bulk proteome dataset (log_2_(fold-change) < -1, p-value < 0.05), with nucleoporins highlighted in cyan. Proteins are plotted based on the mean XRNAX log_2_(H/L) SILAC ratio and mean bulk proteome log_2_(fold-change). (D) Immunofluorescence microscopy of G3BP1 and RANBP2 in response to DMSO, JC19 (100 µM, 1 hour), and EN450 (100 µM, 1 hour). Images were acquired on an LSM880 confocal microscope at 63X objective and 2X manual zoom. All scale bars = 10 µm. (E) Immunofluorescence microscopy of G3BP1 and PSMB2 in response to DMSO and JC19 (100 µM, 1 hour). Images were acquired on an LSM880 confocal microscope at 63X objective and 2X manual zoom. All scale bars = 10 µm. (F) HEK293T cells transiently expressing either WT or C11A/C272A nsp14-FLAG were treated with vehicle control (DMSO) or 100 µM JC19 for 1 hour. nsp14-FLAG fluorescence intensity was quantified by ImageJ (*n* > 300). Statistical significance calculated with unpaired Student’s t-tests *** p<0.0005, **** p<0.0001. (G) HEK293T cells were transiently transfected with nsp14-FLAG, treated with DMSO, MG132 (10 µM, 5 hours), a low dose of JC19 (25 µM, 30 minutes), a high dose of JC19 (100 µM, 1 hour), or EN450 (100 µM, 1 hour), fixed, permeabilized, and subject to immunofluorescence microscopy to visualize nsp14 and endogenous p62. Images were acquired on an LSM880 confocal microscope at 63X objective and 2X manual zoom. All scale bars = 10 µm.

Given the widespread use of such modestly potent electrophilic scout fragments in cell-based chemoproteomic target hunting pipelines^57,60,67,70,73,141–153^, we next broadened the scope of compounds tested for SG-forming activity to include additional scout fragments KB2, KB5, KB7, and covalent-reversible compound KB23 that features a cyanoacrylamide^67^ (**Scheme S1**). All scout fragments robustly induced formation of G3BP1 puncta (**Figure 6B**). Thus, we further broadened the scope of compounds assessed to include RA190, which, like other cysteine-reactive electrophiles, increased proteasome activity in our proteasome activation assays (**Figure S12A**), bardoxolone, a widely utilized NRF2 activator^154^, sulforaphane, a covalent modulator of the ubiquitin-proteasome and autophagic pathways with anticancer activity^155^, as well as chemotherapeutic agents such as FDA-approved ibrutinib and afatinib, which are covalent kinase inhibitors, auranofin, which targets TXNRD1/2^156,157^ and was recently reported to increase the activity of the DNA damage checkpoint kinase CHK1 via oxidation of an allosteric cysteine residue^147^. We subjected cells to compound dosing consistent with prior reports—for afatinib and ibrutinib we chose to treat with 10 µM compound, which, while a dosage used widely^158–163^ are in substantial excess of those required to achieve full target engagement^84^. With the exception of ibrutinib, all cysteine-reactive compounds induced G3BP puncta consistent with SG formation (**Figure 6B, S31A**). Notably, 10 µM afatinib, which is substantially in excess of the concentration required to achieve full EGFR inhibition, was the minimum concentration required to induce SG formation (**Figure S31B**).

In addition to G3BP1 accumulation into puncta, stress granules are defined by sequestration of RNA and RNA-binding proteins. Therefore, we next asked whether compound treatment would induce protein-RNA interactions, consistent with those previously reported for SGs^164^. XRNAX^165^ analysis, which pairs UV crosslinking to trap RBPs on RNA with LC-MS/MS analysis, revealed marked increases in RNA association (log_2_(H/L) SILAC ratio > 1) in response to JC19 treatment (**Table S5**) for many proteins previously reported as SG proteins, including FUS, and FUBP1, and EIF4B. Additionally, nearly all (13 of 16) nucleoporins identified showed increased association with RNA upon treatment with JC19. In total, 15 of the 96 proteins that were statistically depleted by JC19 in our bulk proteome dataset were identified in our XRNAX dataset. This subset includes 7 nucleoporin proteins, 5 of which (NUP93, NUP214, NUP153, NUP98, RANBP2) displayed increased RNA association upon JC19 treatment, suggestive of sequestration into stress granules. (**Figure 6C**). Immunofluorescence analysis corroborated the compound-induced accumulation of RanBP2 into the G3BP1 puncta upon both JC19 and EN450 treatment (**Figure 6D**). Looking beyond the nucleoporins, additional proteins that were depleted by JC19 also showed increased association with RNA by XRNAX, including AKAP8, AKAP8L, and TP53 (**Figure 6C**). The accumulation of TP53 is particularly noteworthy given its tumor suppressor activity and the prior report of naturally occurring isothiocyanate induced selective depletion of TP53 in a manner insensitive to proteasome inhibition^166^.

Returning to nsp14 and nsp16, we next asked whether either of these SARS-CoV-2 proteins were similarly recruited into the electrophilic compound induced SGs. Gratifyingly, we observe robust compound-induced and cysteine-dependent depletion of nsp14 by immunofluorescence (**Figure 6F**), which aligns with our immunoblot analysis (**Figure 4E**). As prior reports have shown sequestration of proteasome subunits to SGs to modulate SG dynamics^167,168^, and our data implicated a partially proteasome-dependent mechanism of nsp14 depletion, we asked if compounds induced sequestration of the proteasome into SGs to mediate nsp14 depletion. We observe robust association of the proteasomal subunit PSMB2 with G3BP1 puncta in a compound-dependent manner (**Figure 6E**). However, no substantial colocalization was observed between nsp14 and G3BP1 (**Figure S32**) in the presence or absence of compound treatment. While we did observe nsp16 sequestration into discrete puncta upon JC19 treatment, these puncta did not colocalize substantially with G3BP1 puncta, suggesting that they are structurally distinct from stress granules (**Figure S32**).

The general lack of nsp14 and nsp16 accumulation into SGs, which contrasts with the robust accumulation for the NPC components, hinted at the possibility of multiple parallel processes that lead to protein aggregation and degradation. To glean further insights into possible alternate mechanisms, we revisited the insoluble proteomics dataset that highlights protein accumulation in the insoluble fraction upon JC19 treatment (**Figure S28**). Gene Ontology analysis revealed that the aggresome was the only enriched cellular component among the aggregated protein subset (**Figure S33**). The aggresome is a perinuclear-localized body of misfolded and aggregated proteins that forms when the cellular degradation machinery is overwhelmed^169,170^. In order to determine whether nsp14 and/or nsp16 are recruited to the aggresome upon compound treatment, we visualized colocalization of the viral proteins with the ubiquitin-binder p62, a well-known component and modulator of the aggresome^171^. Gratifyingly, we observe increased colocalization of both nsp14 and nsp16 with the p62 aggregates upon JC19 and EN450 treatment (**Figure 6G and Figure S34**), substantiating that aggregation is dependent upon compound treatment. These data support the hypothesis that cell-based treatment with cysteine-reactive electrophilic compounds can induce aggresome formation and implicate compound-dependent protein destabilization in the mechanism of nsp14 depletion. By revealing that protein aggregation is a ubiquitous feature of cysteine-reactive small molecules, which is intimately intertwined with protein degradation, our work highlights both the complexities and opportunities for ongoing and future efforts to achieve precise pharmacological control over protein abundance.

## Discussion

Using complementary chemoproteomic strategies, we have found that cysteine-reactive molecules, including drugs, trigger widespread rapid and pervasive aggregation and degradation of host and viral proteins. Focusing first on SARS-CoV-2 viral proteins, we found that the exonuclease nsp14, a key regulator of virulence, is rapidly depleted from the soluble and insoluble proteome in response to the dual electrophile JC19. Mechanistic studies reveal that JC19 effects proteasome-mediated nsp14 degradation via covalent modification at two cysteines in nsp14 (C11/C272) alongside labeling of host protein disulfide isomerases and compound-induced protein accumulation in aggresomes.

Looking beyond nsp14, we find that the host proteome is also dramatically remodeled in response to JC19 and structurally related analogues, including the recently reported molecular glue EN450 when used at the reported dosage^64^. The observed depletion of the nucleoporins was particularly striking given the well-established long-lived nature of the NPC^172,173^ and still poorly defined mechanism of NPC turnover in mammalian cells—in yeast the NPC is degraded by a selective autophagy process^174^. More broadly, we observe formation of stress granules in response to nearly every cysteine-reactive compound evaluated, including the drugs afatinib and auranofin, with the latter recently implicated in increasing nuclear H_2_O_2_ levels^147^. Notably, afatinib-induced stress granules were only observed to occur at 10 µM, which, while a dosage that is reported in a number of prior studies^158–163^, is >50-fold above^175^ the dose require to effect full covalent modification of EGFR^84^. The parallel recruitment of proteasome subunits into stress granules, which aligns with prior reports of stress-induced phase separation of the proteasome^167^, allows us to put forth a model whereby covalent modifiers cause widespread protein aggregation and subsequent proteasome-mediated degradation, driven by elevated local concentrations of hyper-active proteasomes.

Taken together, our study implicates the confluence of covalent protein alkylation and activation of cellular stress response mechanisms in causing widespread proteome remodeling in response to electrophilic compounds. In contrast with classic heterobifunctional and glue-based degraders, which requires compound-induced ternary complex formation between a POI and an E3, our data points towards covalent-modification-induced protein aggregation followed by recruitment of multiple components of the cellular proteostasis machinery including chaperones, ubiquitin ligases, and the proteasome itself. Conceptually the mechanism of clearing compound-induced aggregates shares some similarities with the degradation of aggregated BCL6, after compound-induced protein self-assembly^176^. Parsing the relative contributions of non-specific cell stress response mechanisms from target-specific aggregation, as was the case for BCL6, will be essential to realize the potential for such aggregation-degraders.

Cell-based chemoproteomic screening using low-mid-micromolar doses (e.g. 10-50 µM) of covalent compounds, such as the molecules reported by our study, is an increasingly common strategy for target hunting^57,60,67,70,73,141–153^. Given the ubiquitous nature of the cell stress response mechanisms observed here, we expect that many of the molecules used in prior studies at the indicated concentrations could foreseeably also have stress-causing activity that may, in some cases, complicate target identification and subsequent delineation of on- versus off-target activity. As these “scout” molecules engage most targets with only modest potency (e.g. ∼10-20 µM apparent IC50), they fail to fulfill the important and more stringent criteria of bona fide chemical probes, such as those put forth by the Chemical Probes portal and related resources^177–180^. Despite this limitation, we do still believe that, with suitably stringent controls, covalent fragments can serve as useful pathfinder compounds^177^ for target discovery and remain meritorious of their new-found position in the target-discovery toolbox. We look forward to continuing to build upon established best practices to ensure on-target mechanisms, including screening “smarter” compound libraries that feature more tempered cysteine reactive electrophiles and enantiomeric compound pairs, selection of targets that show with clear SAR, and careful use of covalent compound-resistant cysteine mutants, that together help delineate on-target activity from more general electrophilic stress^67,70,150–152,181–185^.

Together with these valuable practices, we additionally suggest the following strategies to help delineate on-and off-target activities of covalent fragments: (1) The pairing of cell- and lysate-based chemoproteomic studies to delineate cell-specific activity—additional care should be taken in progressing hits that fail to reconfirm in vitro; (2) Assessment of the number of cysteines per protein identified with high ratios together with measures of bulk changes in protein abundance to delineate direct labeling from changes in abundance or protein stability; (3) The use of protein-directed ABPP^146^ analysis using tailored click probes to confirm target engagement and occupancy; (4) Restricting follow-up studies to cysteines labeled at near complete (∼100%) occupancy by prioritizing lead compounds that show reproducible high fold change competition values for the cysteine of interest across replicate experiments, (5) Confirmation of clear SAR using multiple active and inactive control compounds, including enantiomeric pairs of compounds when possible; (6) Assessment of generalized cell stress (e.g. stress granule formation, global polyubiquitylation, aggresome formation, and relative compound reactivity towards thiols such as glutathione; (7) Community-wide publication-associated data deposition into centralized repositories, such as CysDB^66^, to facilitate identification of pan-electrophile sensitive proteins for exclusion from future studies; (8) For covalent compounds targeting non-essential cysteines, the use of compound-resistant cysteine mutants (e.g., cysteine-to-alanine/serine mutants) as controls; and (9) For covalent degraders, ternary complex formation should be assessed in vitro, and the POI and implicated ligase assessed via stringent genetic and biochemical characterization. We would additionally encourage caution when no clear SAR is apparent, when relative degradation correlates with relative compound reactivity towards thiols, with more reactive compounds resulting in more degradation, and when high compound doses are required to effect degradation.

Looking to the future, we are excited to further delineate additional protein biomarkers and mechanisms of cell stress activation. We expect that our rich multimodal datasets will serve as useful resource for the delineation of covalent labeling sites from stress-induced changes to the proteome and will serve as a launch point for the discovery of cysteines that are requisite for protein aggregation and liquid-liquid phase separation processes. We are particularly optimistic that our work will help to bolster the chemoproteomic discovery of actionable target-compound pairs that can serve as starting points and additions to the existing toolbox of high quality covalent chemical probes and drugs.

## Acknowledgements

This study was supported by a Beckman Young Investigator Award (K.M.B.), DOD-Advanced Research Projects Agency (DARPA) D19AP00041 (K.M.B.), National Institutes of Health DP2 OD030950-01 (K.M.B.), Packard Fellowship (K.M.B.), and NIGMS UCLA Chemistry Biology Interface T32GM136614 (A.R.J). We thank Dr. Irene Zohn for providing the HECTD1 plasmid. We additionally thank all members of the Backus labs for helpful suggestions and the UCLA Broad Stem Cell Resource Center Microscopy Core for assistance with microscopy. Figure 1A was made using Biorender.com.

## Author Contributions

A.R.J., F.S, and K.M.B. conceived of the project. A.R.J., F.S., and K.M.B designed experiments. A.R.J., F.S., C.T. and A.C.T collected data. N.R.B, E.D, and J.C. performed synthesis. A.R.J., F.S. and C.T. performed data analysis. C.T. wrote software. A.R.J. and F.S, contributed to the figures. A.R.J., F.S, and K.M.B. wrote the manuscript with assistance from all authors.

## Data Availability

The MS data have been deposited to the ProteomeXchange Consortium (http://proteomecentral.proteomexchange.org) via the PRIDE^186^ partner repository with the dataset identifiers PXD046278 (isoTOP-ABPP data in Figures 1 and 5) and PXD046393 (all other proteomic data corresponding to Figures 3-6).

