## Supplementary Information for "Pervasive aggregation and depletion of host and viral proteins in response to cysteine-reactive electrophilic compounds"

[a]Department of Biological Chemistry Department David Geffen School of Medicine, UCLA, Los Angeles, CA 90095 (USA); [b]Department of Chemistry and Biochemistry, UCLA, Los Angeles, CA 90095 (USA); [c]DOE Institute for Genomics and Proteomics, UCLA, Los Angeles, CA 90095 (USA); [d]Jonsson Comprehensive Cancer Center, UCLA, Los Angeles, CA 90095 (USA); [e]Eli and Edythe Broad Center of Regenerative Medicine and Stem Cell Research, UCLA, Los Angeles, CA 90095 (USA).

[%]These authors contributed equally to this work

**Table of Contents**

**(A) Supplementary Figures 2-27**
**(B) Supplementary Tables 27-37**
**(C) Biology Methods 37-48**
**(D) Chemistry Methods 45-57**
**(E) NMR Spectra 57-75**
**(F) References 76-77**

### (A) Supplementary Figures

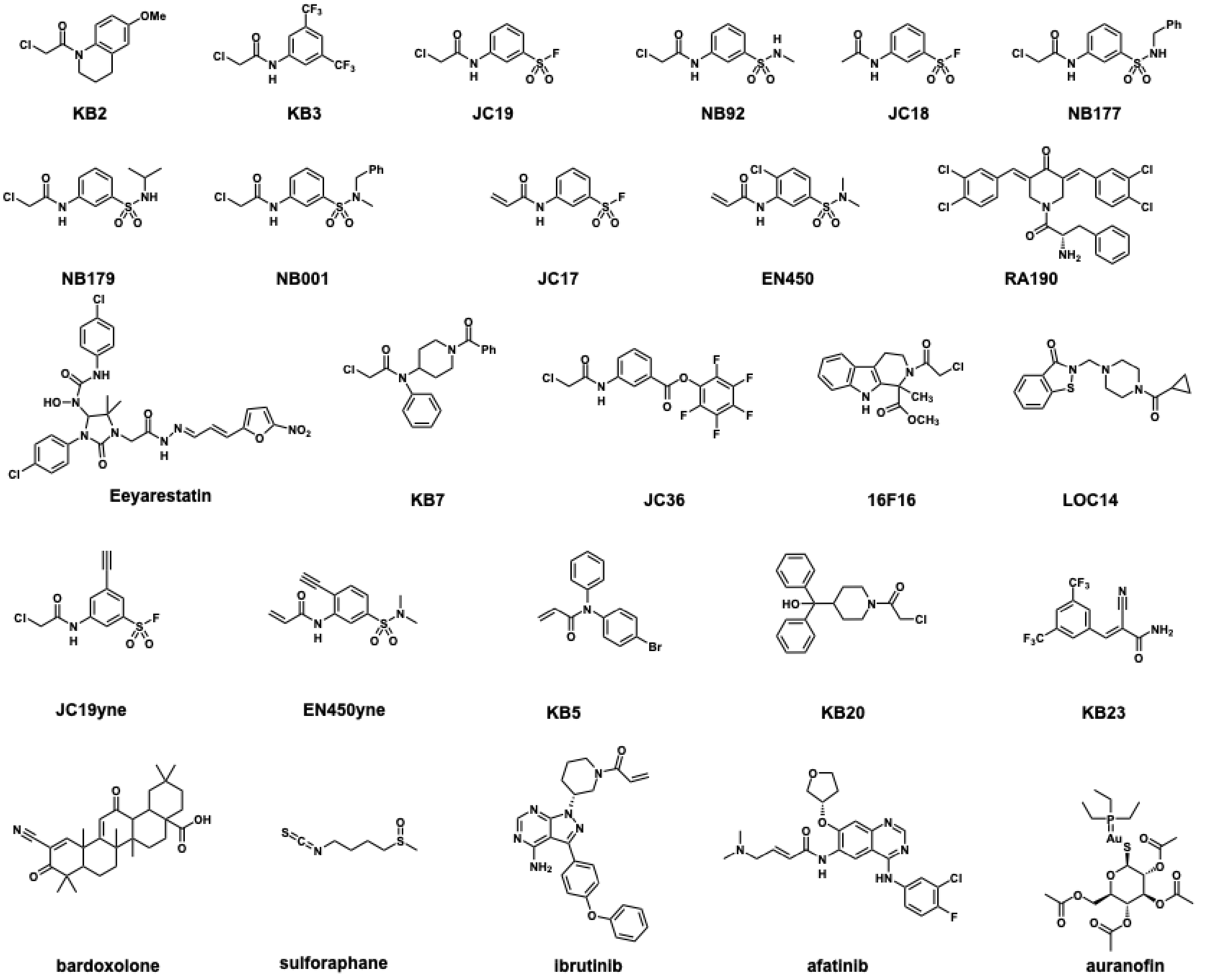


**Scheme S1.** Structures of all compounds used for the study, ordered in the order of appearance in the text.

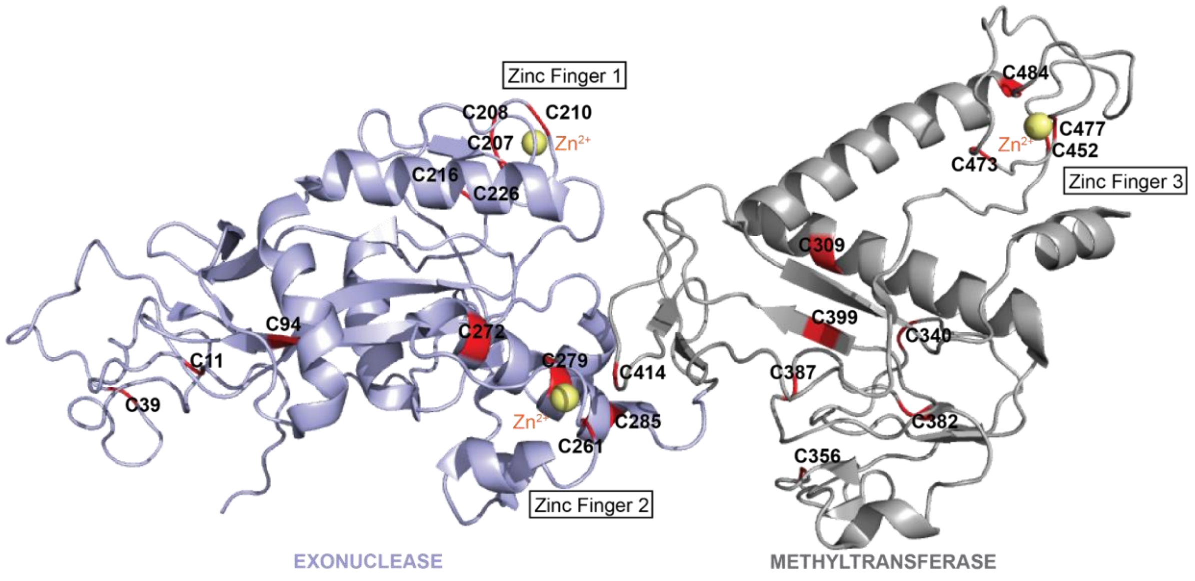


**Figure S1. Structure of SARS-CoV-2 nonstructural protein 14 (nsp14) used in isoTOP-ABPP experiments.** Nsp14 (PDB: 7QGI)[1] with cysteine residues highlighted in red and labeled with residue number. The exonuclease domain is colored light blue and the methyltransferase domain is colored gray. Zinc ions of the zinc fingers are colored yellow with an orange Zn label. Protein structure generated using The PyMOL Molecular Graphics System, Version 2.0 Schrödinger, LLC.

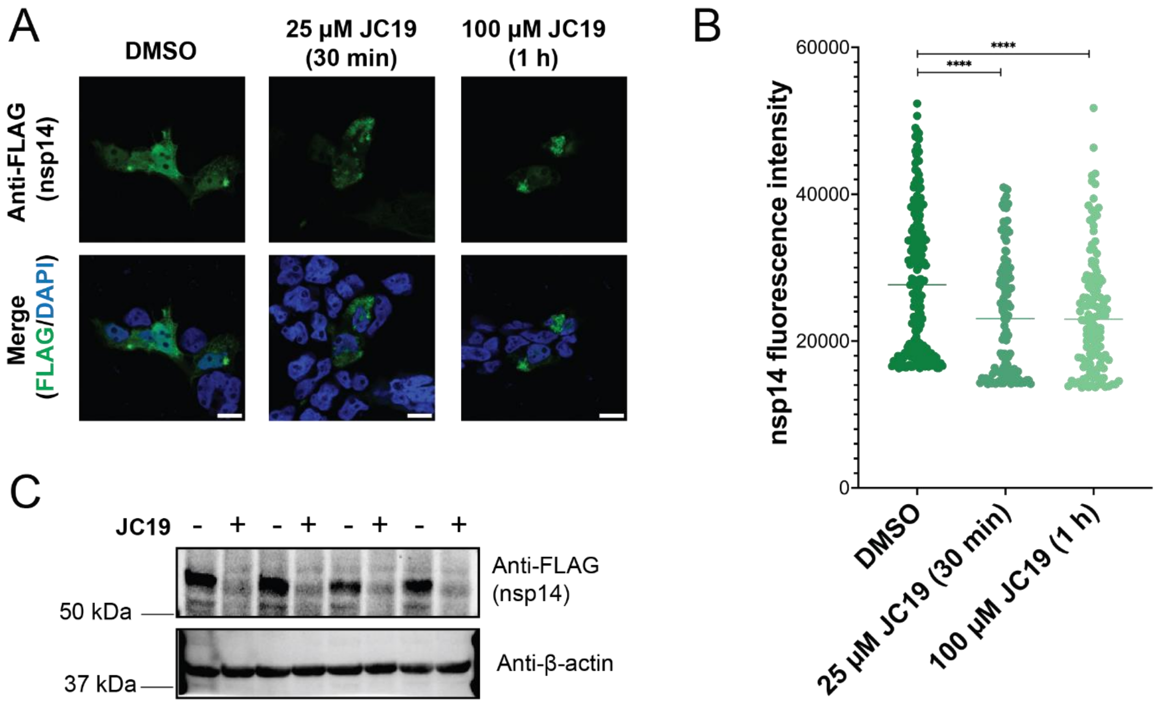


**Figure S2. Nsp14 is depleted in HEK293T and HeLa cells.** (A) HEK293T cells transiently expressing nsp14-FLAG were treated with vehicle control (DMSO) or the indicated concentrations of JC19 for the indicated times. Cells were then fixed, permeabilized, and stained for immunofluorescence imaging. Cells were imaged on an LSM880 confocal microscope at 63X objective with 2X manual zoom. Scale bar = 10 µm. (B) Quantification of nsp14-FLAG fluorescence intensity (ImageJ[2]) taken from 4 images prepared as described in (A) and acquired at 10X objective ($n$ > 100). Statistical significance calculated with unpaired Student's t-tests * $p<0.05$, **** $p<0.0001$. (C) HeLa cells transiently expressing nsp14-FLAG were treated with either vehicle control (DMSO) or 100 µM JC19 for 1 hour ($n$ = 4) and immunoblot analysis was used to visualize abundance of nsp14 in the soluble lysate fraction.

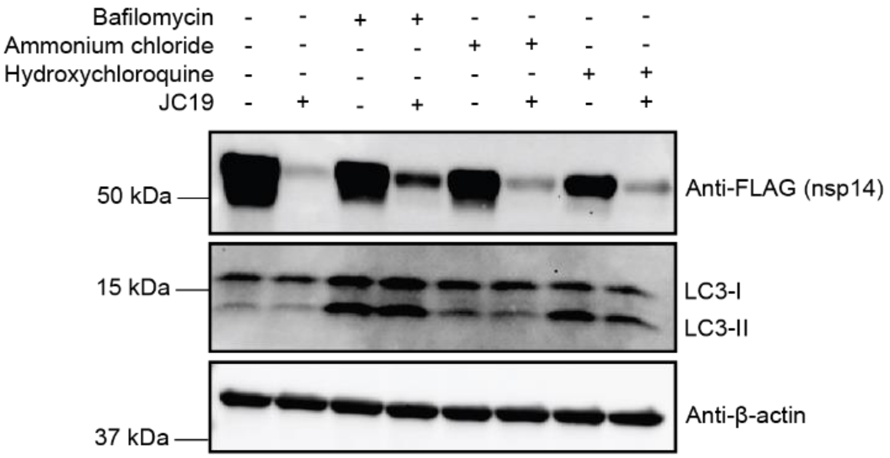


**Figure S3. Autophagy and lysosomal inhibitors impart partial protection from nsp14 depletion.** HEK293T cells transiently expressing nsp14-FLAG were pretreated with either DMSO, bafilomycin (1 µM, 6 hours), ammonium chloride (10 mM, 6 hours), or hydroxychloroquine (100 µM, 6 hours), followed by treatment with either DMSO or 50 µM JC19 for 30 minutes. Immunoblot analysis was used to visualize abundance of nsp14 in the soluble lysate. LC3-II used as a marker of autophagy inhibition.

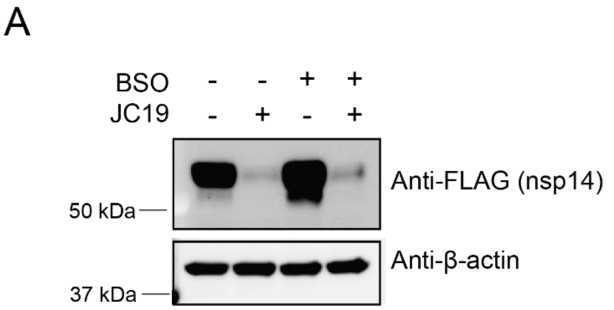


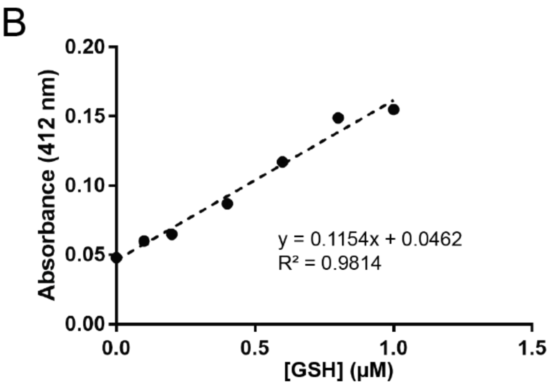


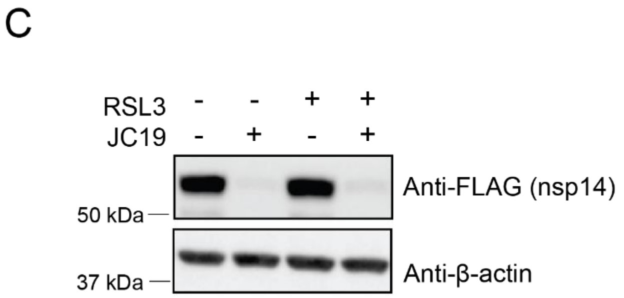


| | Absorbance | [GSH] (µM in assay) | [GSH] (µM per cell) |
|---|---|---|---|
| DMSO | 0.218 | 1.49 | 117 |
| JC19 (100 µM) | 0.217 | 1.48 | 116 |
| BSO (500 µM) | 0.062 | 0.14 | 10.8 |

**Figure S4. JC19-mediated nsp14 degradation is not a result of altered glutathione levels.** (A) HEK293T cells transiently expressing nsp14-FLAG were pretreated with either DMSO or 500 µM buthionine sulfoximine (BSO) for 24 hours to reduce glutathione levels, and then treated with either

DMSO or 100 µM JC19 for 1 hour. Immunoblot analysis was used to visualize abundance of nsp14 in the soluble lysate fraction for each condition. (B) Glutathione detection kit (G Biosciences, Cat#786-075) was used to detect glutathione (GSH) levels in cells treated with DMSO or 100 µM JC19 for 1 hour (500 µM BSO treatment for 24 hours used as positive control). Table depicts the standard curve of GSH measured in the assay, and the table below summarizes measurements of experimental samples (vehicle, JC19, and BSO). (C) HEK293T cells transiently expressing nsp14-FLAG were pretreated with 5 µM RSL3 for 5 hours to induce ferroptosis, and then treated with either DMSO or 50 µM JC19 for 1 hour. Immunoblot analysis was used to visualize abundance of nsp14 in the soluble lysate fraction for each condition.

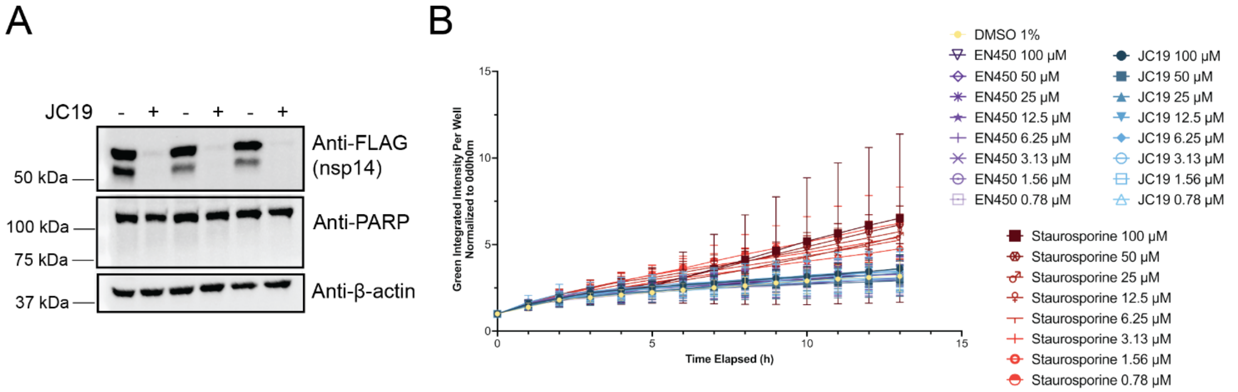


**Figure S5. JC19 does not induce cytotoxicity to afford nsp14 depletion.** (A) HEK293T cells transiently expressing nsp14-FLAG were treated with either 100 µM JC19 or equal volume of DMSO for 1 hour (*n* = 3). Immunoblot analysis was used to visualize abundance of nsp14 and PARP cleavage in the soluble lysate fraction for each condition. (B) HeLa cells were treated with the indicated concentrations of compounds (*n* = 3 per conditions) and imaged on a IncuCyte SX5 Live-Cell Imaging and Analysis System (Sartorius) using Cytotox Green reagent (Cat. No. 4633) (final dilution 1:4000). Measurements were taken every hour over the course of 13 hours. Staurosporine was used as a positive control.

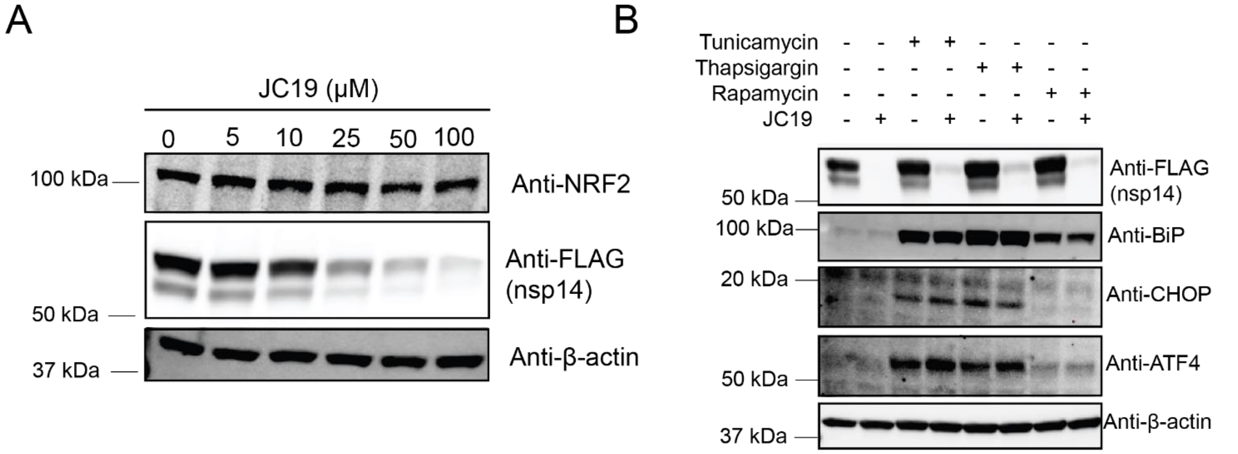


**Figure S6. Common cell stress-sensing pathways are not responsible for nsp14 depletion.** (A) HEK293T cells transiently expressing nsp14-FLAG were treated with the indicated concentrations of JC19 for 1 hour. Immunoblot analysis was used to visualize abundance of nsp14 and induction of NRF2 expression in the soluble lysate fraction for each condition. (B) HEK293T cells transiently expressing nsp14-FLAG were pretreated with either DMSO, tunicamycin (12 µg/mL, 8 hours), thapsigargin (2 µM, 8 hours), or rapamycin (1 µM, 8 hours), then either treated with DMSO or 50 µM JC19 for 1 hour. Immunoblot analysis was used to visualize abundance of nsp14 and induction of UPR markers in the soluble lysate fraction for each condition.

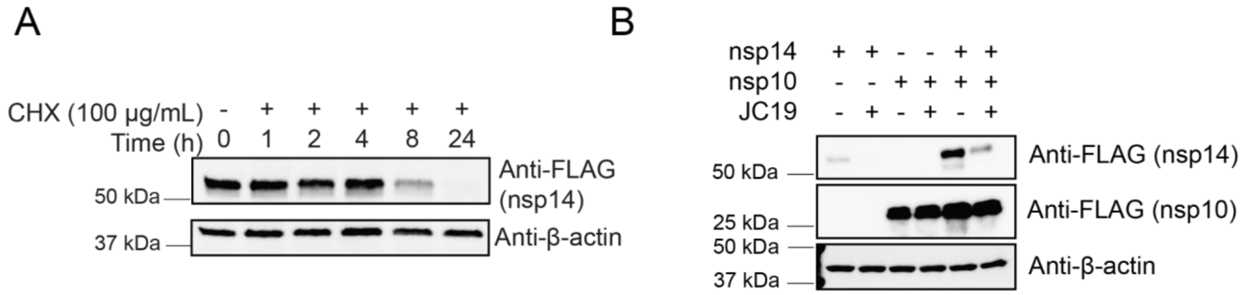


**Figure S7. Inherent nsp14 instability does not cause rapid turnover.** (A) HEK293T cells transiently expressing nsp14-FLAG were treated with 100 µg/mL cycloheximide (CHX) for the indicated times to inhibit translation. Immunoblot analysis was used to visualize abundance of nsp14 in the soluble lysate fraction to assay nsp14 turnover. (B) Nsp14 and nsp10 were transfected individually and coexpressed in HEK293T cells and treated with either DMSO or 100 µM JC19 for 1 hour. Immunoblot analysis was used to visualize abundance of nsp14 in the soluble lysate fraction for each condition.

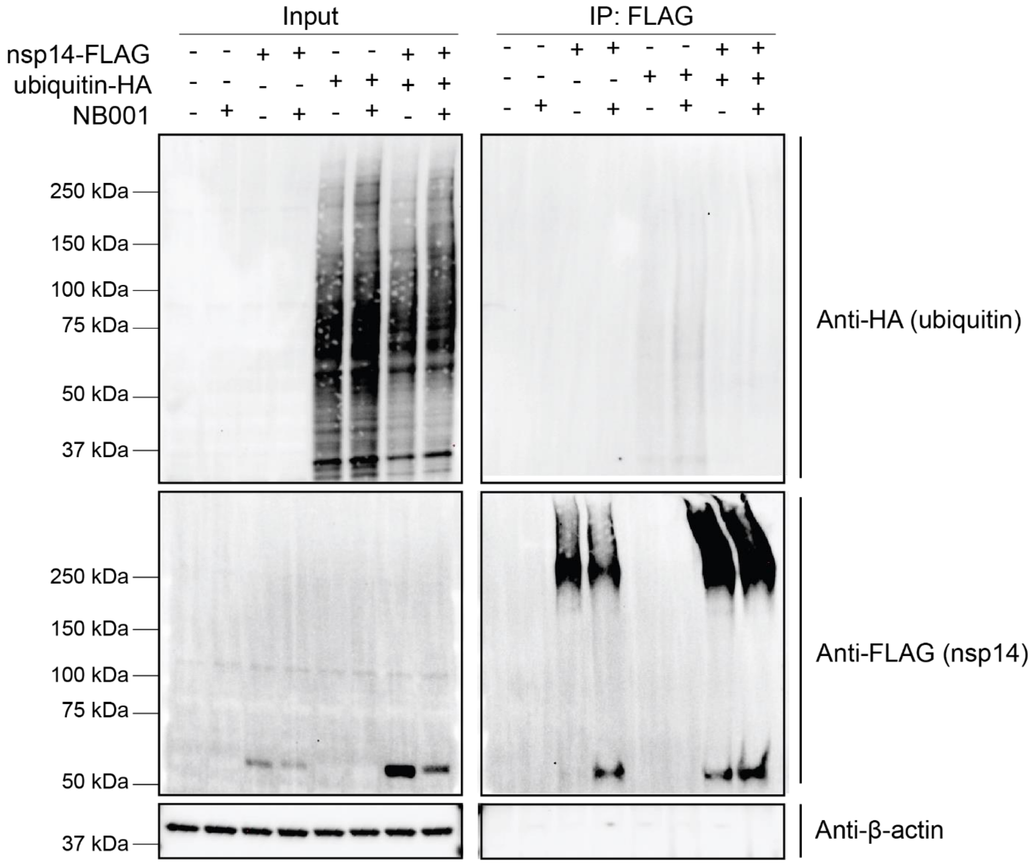


**Figure S8. Nsp14 is not ubiquitinated by NB001.** HEK293T cells transiently expressing nsp14-FLAG, ubiquitin-HA, neither, or both were treated with either DMSO or 100 µM NB001 for 1 hour, immunoprecipitated on FLAG resin, and ran on immunoblot to assay ubiquitination of nsp14.

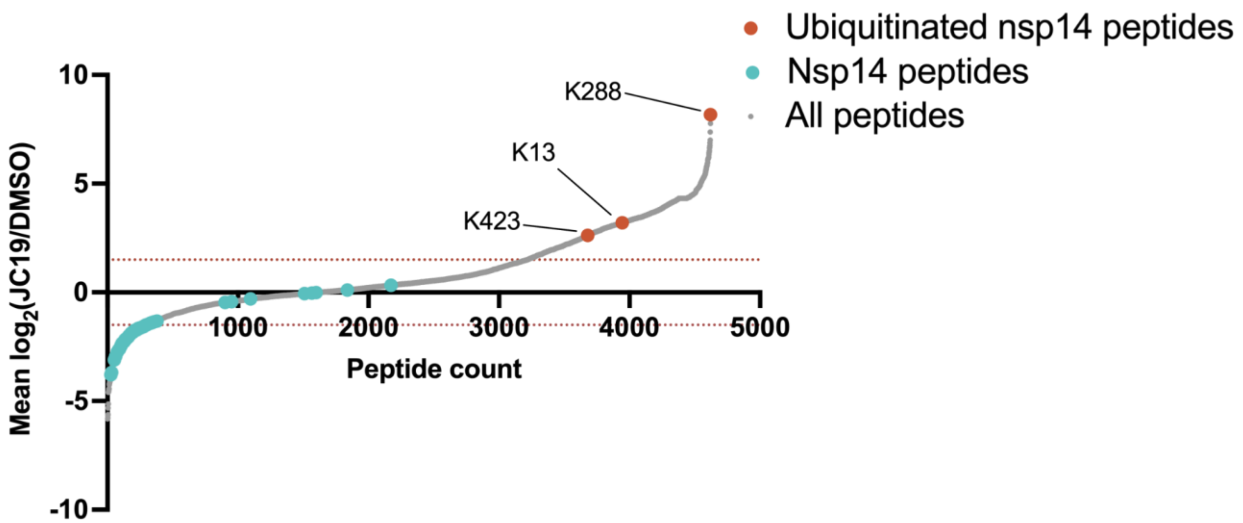


**Figure S9. Nsp14 lysines are ubiquitinated in response to JC19 treatment.** 'Heavy' SILAC HEK293T cells transiently expressing nsp14 were treated with 100μM JC19 for 1 hour ($n$ = 3), while 'light' SILAC HEK293T cells transiently expressing nsp14 were treated with an equal volume of DMSO for 1 hour ($n$ = 3). Lysates were combined, immunoprecipitated on FLAG resin, proteolyzed, and subject to LC-MS/MS analysis. Heavy/light MS1 ratios of detected peptides are depicted on a waterfall plot, highlighting all nsp14 peptides and ubiquitinated nsp14 peptides. All MS data can be found in **Table S2**.

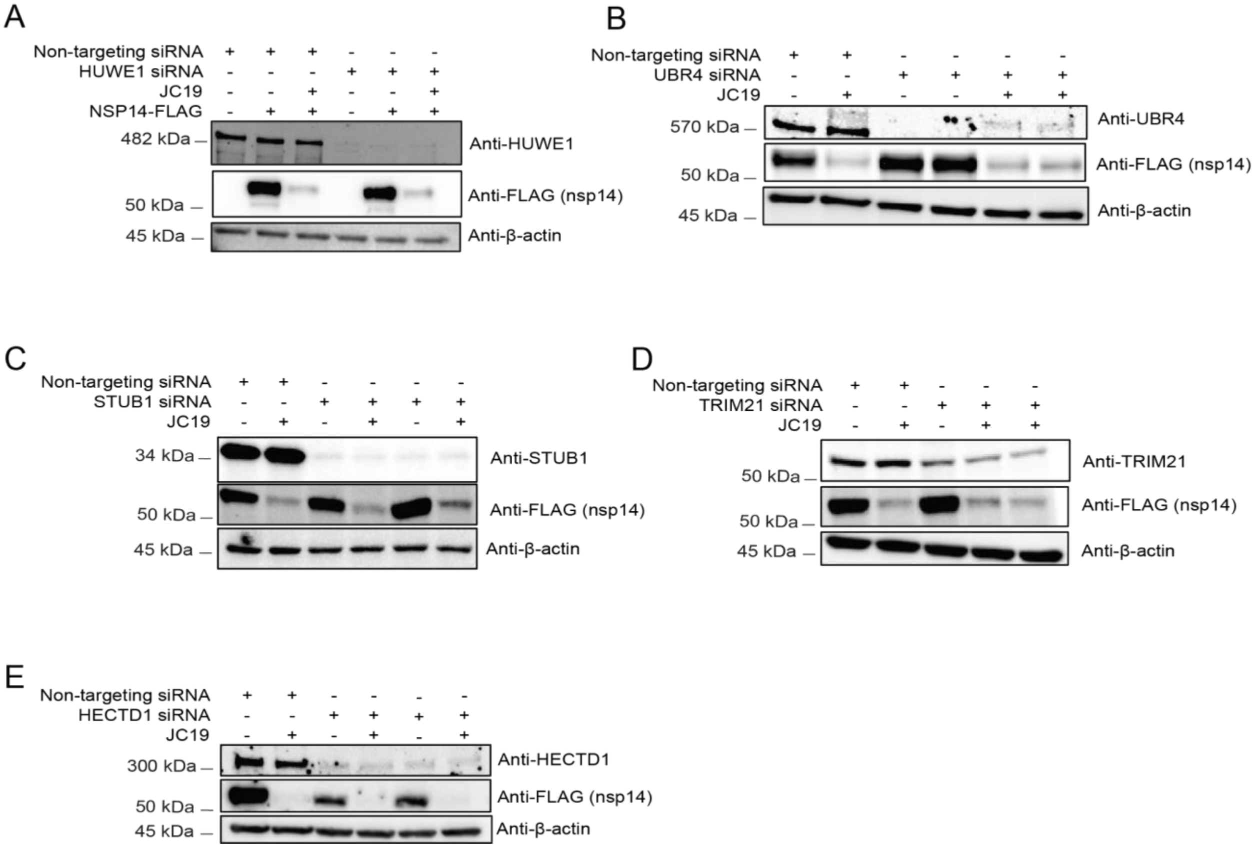


**Figure S10. Knockdown of AP-MS identified E3 ubiquitin ligases does not protect nsp14 from JC19-mediated degradation. (**A) Immunoblot of siRNA knockdown of HUWE1 in HEK293T cells and the effect on nsp14-FLAG abundance in the presence and absence of JC19 treatment (25 µM for 0.5 hr). (B) Immunoblot of siRNA knockdown of UBR4 in HEK293T cells and the effect on nsp14-FLAG abundance in the presence and absence of JC19 treatment (25 µM for 0.5 hr). (C) Immunoblot of siRNA knockdown of STUB1 in HEK293T cells and the effect on nsp14-FLAG abundance in the presence and absence of JC19 treatment (25 µM for 0.5 hr). (D) Immunoblot of siRNA knockdown of TRIM21 in HEK293T cells and the effect on nsp14-FLAG abundance in the presence and absence of JC19 treatment (25 µM for 0.5 hr). (E) Immunoblot of siRNA knockdown of HECTD1 in HEK293T cells and the effect on nsp14-FLAG abundance in the presence and absence of JC19 treatment (100 µM for 1 hr).

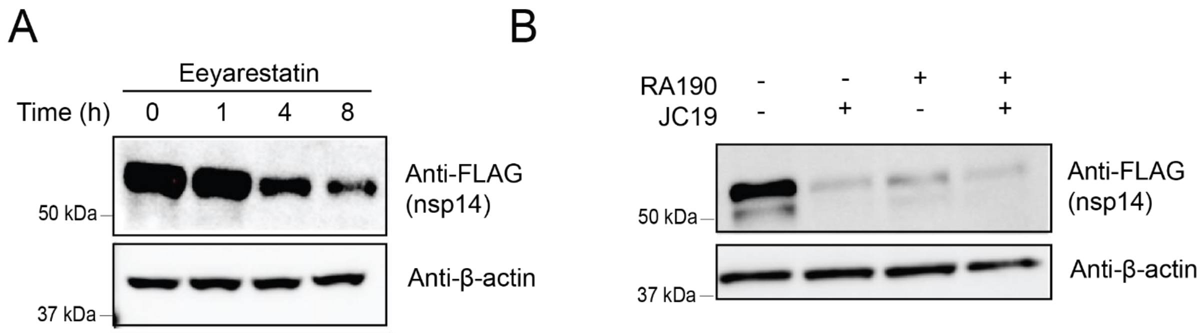


**Figure S11. Both eeyarestatin and RA190 induce depletion of nsp14.** (A) HEK293T cells transiently expressing nsp14-FLAG were treated with 50 µM eeyarestatin for the indicated time points and immunoblot used to assay nsp14 abundance. (B) HEK293T cells transiently expressing nsp14-FLAG were treated with either 2 µM RA190 for 2 hours, 50 µM JC19 for 30 minutes, or pretreated with either DMSO or 2 µM RA190 for 2 hours and then treated with either DMSO or 50 µM JC19 for 30 minutes. Immunoblot used to assay nsp14 abundance.

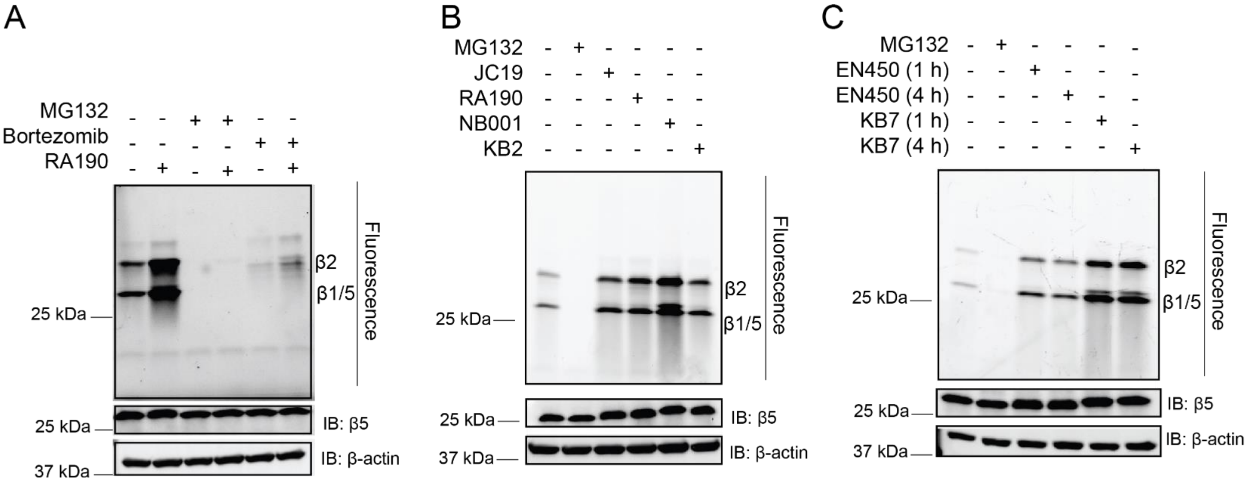


**Figure S12. Cysteine-reactive small molecules activate the proteasome.** (A) HEK293T cells were pretreated with DMSO, MG132 (10 µM for 5 hours), or bortezomib (5 µM for 5 hours), followed by treatment with either DMSO or RA190 (5 µM for 1 hour). Cells were then treated with a fluorescent proteasome activity probe (500 nM for 1 hour) and in-gel fluorescence used to assay proteasome activity. Immunoblot analysis was used to assay the abundance of PSMB5. (B) HEK293T cells were treated with DMSO, MG132 (10 µM for 5 hours), JC19 (100 µM for 1 hour), RA190 (5 µM for 1 hour), NB001 (100 µM for 1 hour), or KB2 (100 µM for 1 hour). Cells were then treated with a fluorescent proteasome activity probe (500 nM for 1 hour) and in-gel fluorescence used to assay proteasome activity. Immunoblot analysis was used to assay the abundance of PSMB5. (C) HEK293T cells were treated with DMSO, MG132 (10 µM for 5 hours), EN450 (100 µM for 1 and 4 hours), or KB7 (100 µM for 1 or 4 hours). Cells were then treated with a fluorescent proteasome activity probe (500 nM for 1 hour) and in-gel fluorescence used to assay proteasome activity. Immunoblot analysis was used to assay the abundance of PSMB5.

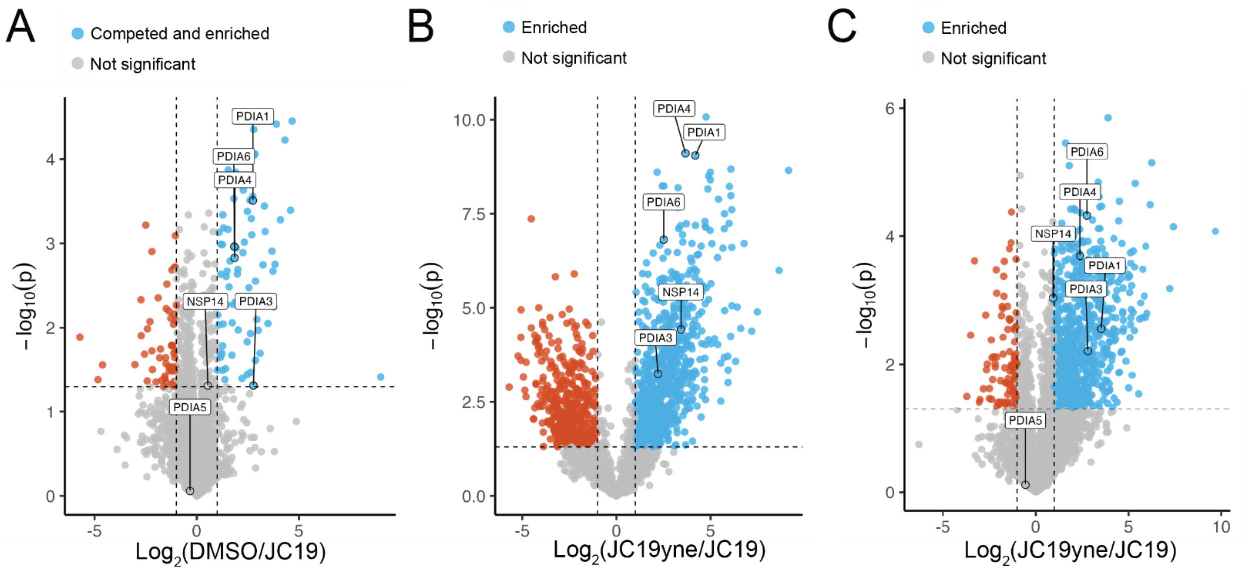


**Figure S13. Nsp14 and the protein disulfide isomerase family are directly labeled by JC19.** (A) HEK293T cells transiently expressing nsp14-FLAG were lysed, cleared by centrifugation, and the soluble lysate treated with either DMSO (*n* = 3) or 100 µM JC19 (*n* = 2) for 20 minutes. All replicates were then treated with 100 µM JC19yne for 20 minutes, conjugated to biotin-azide via CuAAC, cleaned using SP3 clean-up, enriched on streptavidin resin, and proteolytically digested prior to LC-MS/MS analysis. Label-free quantitation was used to generate protein intensities for the volcano plot. (B) HEK293T cells transiently expressing nsp14-FLAG were treated with either 100 µM JC19 (*n* = 6) for 20 minutes or JC19yne (*n* = 6) for 20 minutes. Cells were then lysed and cleared by centrifugation. The soluble lysate was then conjugated to biotin-azide via CuAAC, cleaned using SP3 clean-up, enriched on streptavidin resin, and proteolytically digested prior to LC-MS/MS analysis. Label-free quantitation was used to generate protein intensities for the volcano plot. (C) HEK293T cells transiently expressing nsp14-FLAG were lysed, cleared by centrifugation and treated with either 100 µM JC19 (*n* = 3) for 20 minutes or JC19yne (*n* = 3) for 20 minutes. The soluble lysate was then conjugated to biotin-azide via CuAAC, cleaned using SP3 clean-up, enriched on streptavidin resin, and proteolytically digested prior to LC-MS/MS analysis. Label-free quantitation was used to generate protein intensities for the volcano plot. For all analyses, a student's t-test was performed between control and treatment groups to generate p-values, and proteins with a fold-change > 1 and p-value < 0.05 were considered significantly enriched. All MS data can be found in **Table S3**.

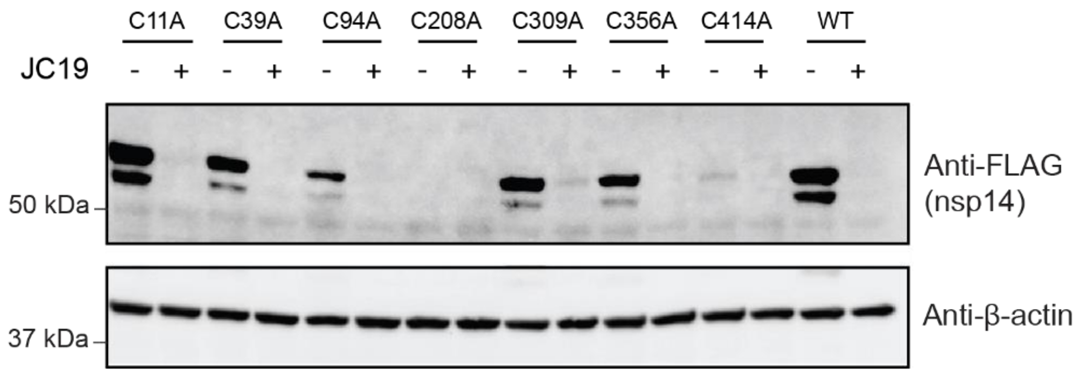


**Figure S14. Mutant screen of a focused set of nsp14 cysteine mutants.** Wild-type (WT) nsp14 or the indicated cysteine to alanine mutant was transiently transfected in HEK293T cells, treated with DMSO or JC19 (100 µM for 1 hour), and subject to immunoblot analysis to assay nsp14 abundance.

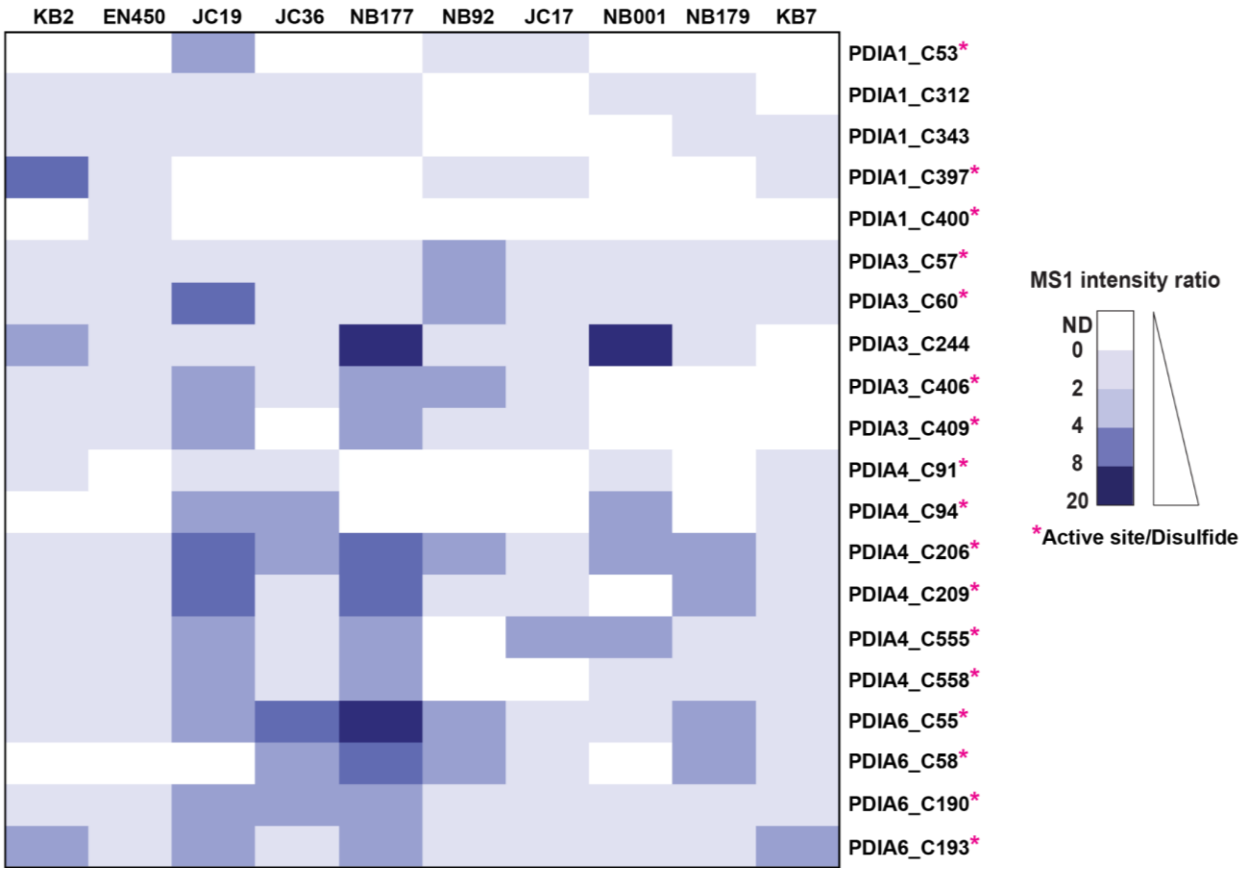


**Figure S15. IsoTOP-ABPP ratios for protein disulfide isomerase (PDI) cysteines.** Heatmap representing the unlogged isoTOP-ABPP ratios for each identified cysteine in PDIA1, PDIA3, PDIA4, and PDIA6 based on the median value from multiple biological replicates of treatment with the indicated compounds. All compound treatments were performed in biological triplicate at 100 µM for 1 hour, except for EN450, which was treated at 100 µM for 4 hours. All MS data can be found in **Table S4.**

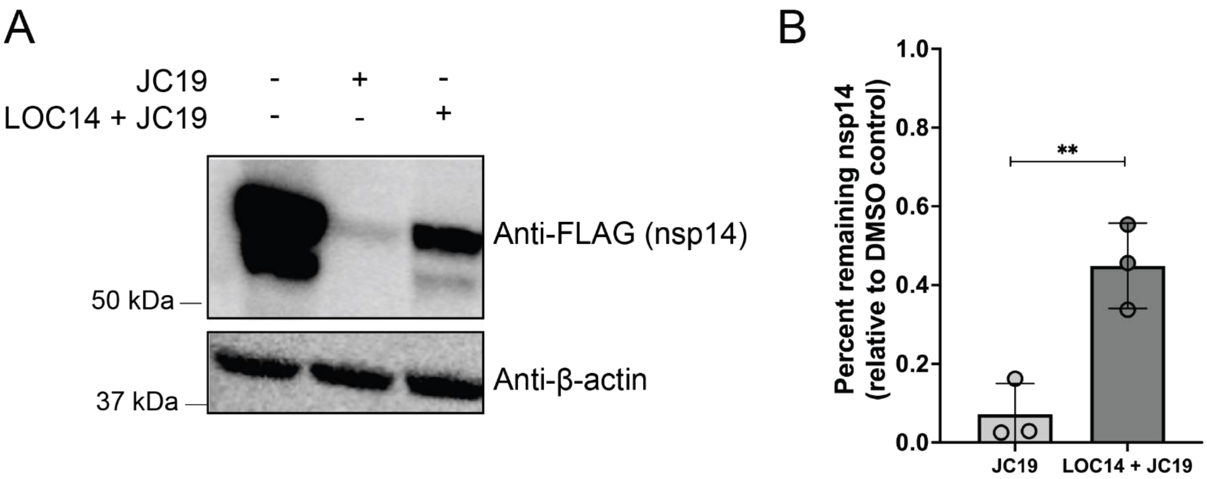


**Figure S16. LOC14 treatment partially protects nsp14 from JC19-mediated depletion. (**A) HEK293T cells transiently expressing nsp14-FLAG were pretreated with either DMSO or LOC14 (10 µM for 16 hours), then treated with either DMSO or JC19 (50 µM for 30 minutes). Immunoblot analysis was performed to detect nsp14 abundance. (B) Quantification of nsp14 abundance (ImageJ[2]) after treatment as described in (A), relative to a DMSO-treated control ($n$ = 3). Statistical significance calculated with Student's t-test, ** $p<0.005$.

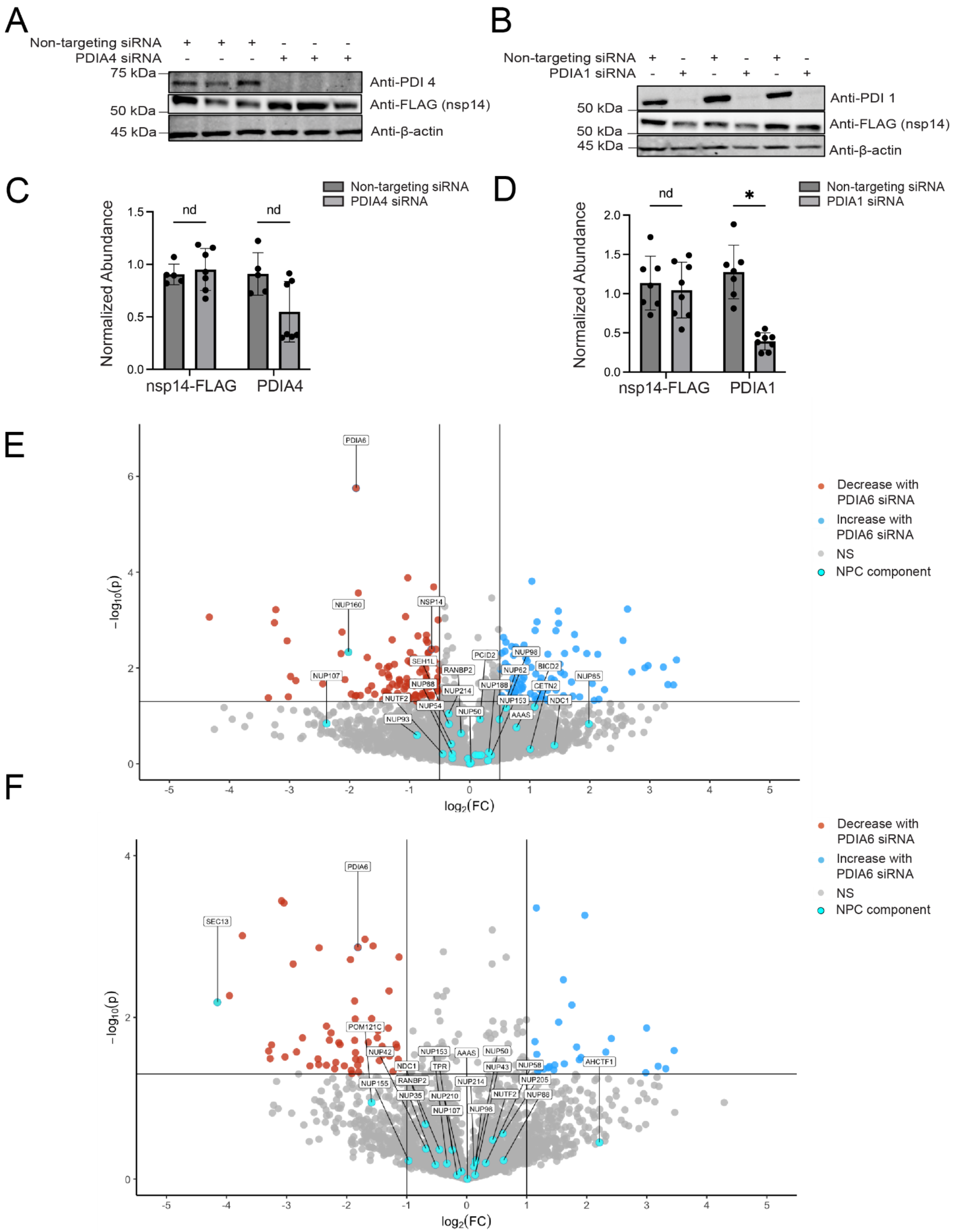


**Figure S17. Knockdown of PDIs effect on nsp14 and host proteins.** (A) Immunoblot analysis of the impact of siRNA-mediated knockdown of PDIA4 on nsp14-FLAG abundance ($n$ = 3). (B) Immunoblot analysis the impact of siRNA-mediated knockdown of PDIA1 on nsp14-FLAG abundance (*n* = 3). (C) Quantification of both PDIA4 and nsp14 abundance (ImageJ[2]) upon siRNA-mediated knockdown of PDIA4 (*n* = 5). (D) Quantification of both PDIA1 and nsp14 abundance (ImageJ[2]) upon siRNA-mediated knockdown of PDIA1 (*n* = 5). Statistical significance calculated with Student's t-tests * p<0.05. (E) Label-free quantification used to measure protein abundance in cells in which siRNA was used to knockdown PDIA6 as compared to scramble siRNA in HEK293T nsp14-overexpressing cells (*n* = 3). FC cutoff at $\log_2(0.5)$. Nucleoporins are highlighted in cyan. (F) Label-free quantification used to measure protein abundance in cells in which siRNA was used to knockdown PDIA6 as compared to scramble siRNA in HEK293T cells (*n* = 3). FC cutoff at $\log_2(1)$. Nucleoporins are highlighted in cyan. All MS data can be found in **Table S3.**

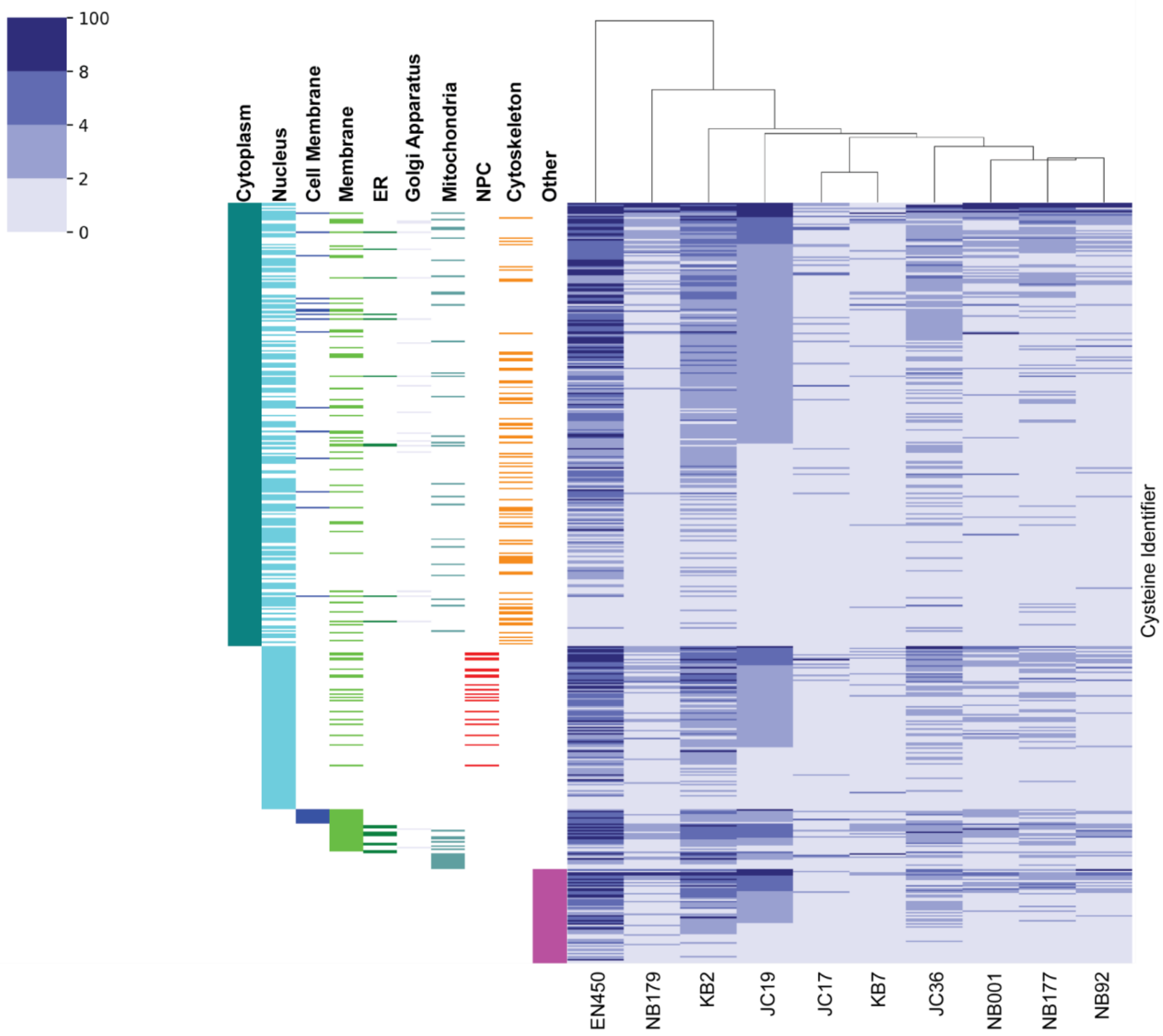


**Figure S18. High-ratio cysteines as identified by isoTOP-ABPP belong to proteins spanning various subcellular compartments.** Heatmap depicting unlogged isoTOP-ABPP cysteine ratios for high coverage cysteines belonging to proteins with multiple cysteines identified. Cell compartment annotations from Uniprot have been provided for each cysteine identifier. For generation of data, each compound was used in triplicate at 100 μM for 1 hour (exception: 4 hour treatment for EN450) in nsp14 expressing HEK293T cells. MS data can be found in **Table S4.**

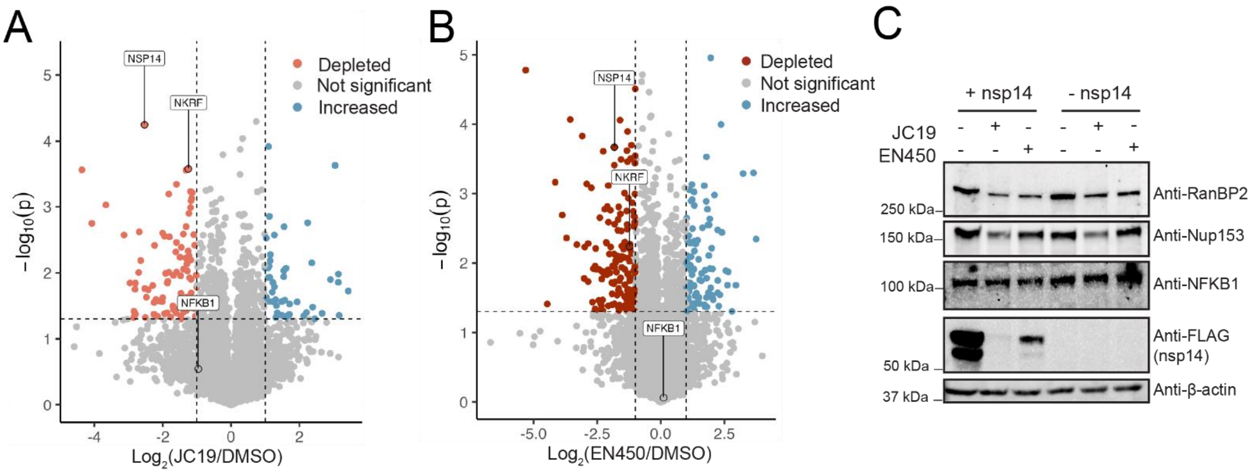


**Figure S19. Nsp14 is depleted from cells when treated with JC19 and EN450.** HEK293T cells transiently expressing nsp14-FLAG were treated with (A) DMSO (n = 3) or 100 µM JC19 (n = 3) or (B) DMSO (n = 3) or 100µM EN450 (n = 3) for 1 hour. Cells were then lysed and prepared for LC-MS/MS analysis via SP3 clean-up. Label-free quantification was used to quantify protein intensities in each replicate. For both plots, a student's t-test was performed between control and treatment groups, and the average fold-change of the label-free intensity for each protein plotted against its p-value. Proteins with a $\log_2$ fold-change < -1 and p-value < 0.05 are considered significantly depleted. (C) HEK293T cells were either mock transfected or transfected with nsp14-FLAG for 36 hours, treated with DMSO, 100 µM JC19 (1 hour), or 100 µM EN450 (4 hours). Immunoblot analysis was used to assay the abundance of nsp14, RANBP2, NUP153, and NFKB1. All MS data can be found in **Table S4**.

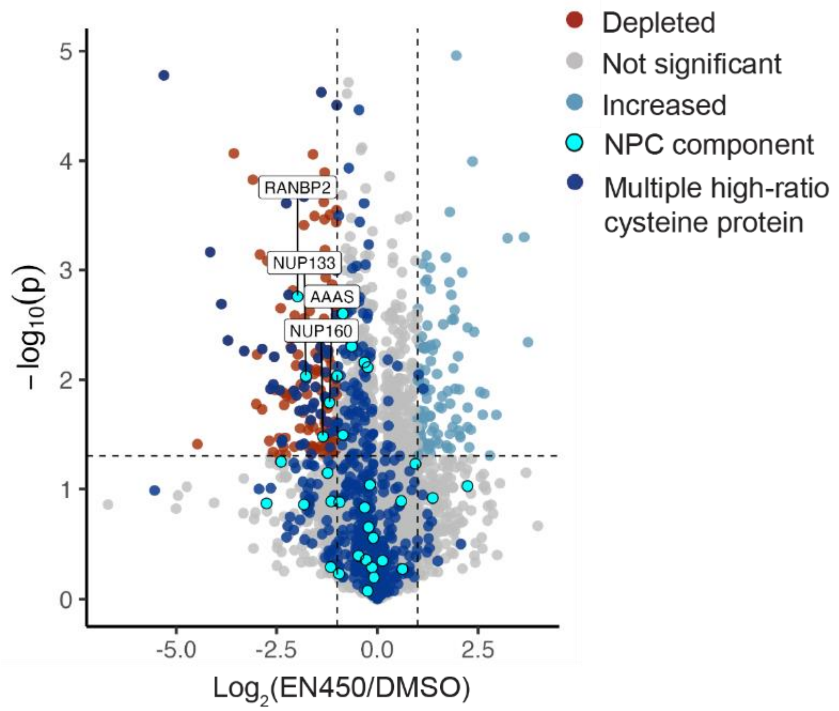


**Figure S20. EN450 induces protein depletion.** HEK293T cells transiently expressing nsp14-FLAG were treated with DMSO (*n* = 3) or 100 µM EN450 (*n* = 3) for 4 hours. Cells were then lysed and prepared for LC-MS/MS analysis via SP3 clean-up. Red points represent proteins that were depleted in response to treatment. Dark blue points represent proteins with multiple high-ratio cysteines in the EN450 isoTOP-ABPP dataset. Cyan points represent components of the nuclear pore complex (NPC). The labeled proteins are the NPC proteins that were significantly depleted in response to EN450 treatment. Label-free quantification was used to measure protein intensities in each replicate and the fold-change of the average intensity for each protein plotted against its respective p-value calculated from a student's t-test. $Log_2$(fold-change) < -1 and p-values < 0.05 are considered significantly depleted. All MS data can be found in **Table S4.**

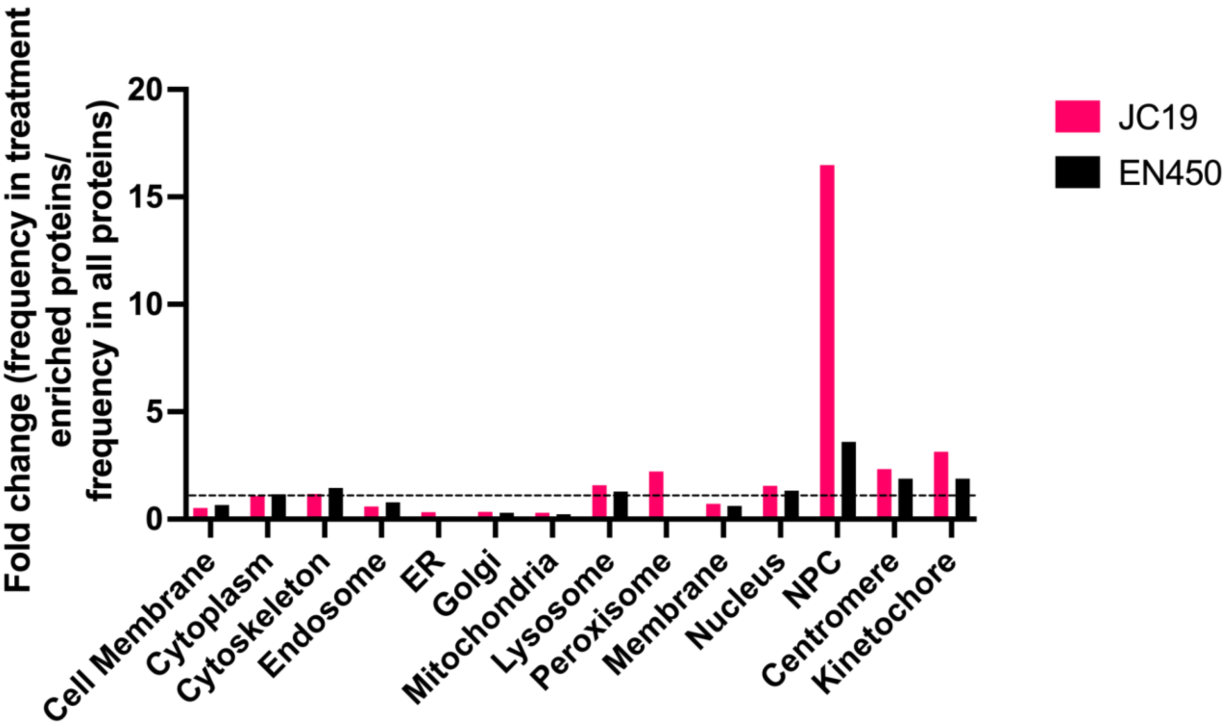


**Figure S21. The nuclear pore complex is an enriched subcellular compartment within proteins depleted by JC19 and EN450.** All proteins identified from JC19 and EN450 soluble label-free bulk proteome MS analysis annotated by subcellular location using Uniprot annotations. The total representation of subcellular location for all proteins that were identified was compared to representation of subcellular locations for the significantly depleted proteins. Fold-change of those two values were plotted. MS data can be found in **Table S4**.

A

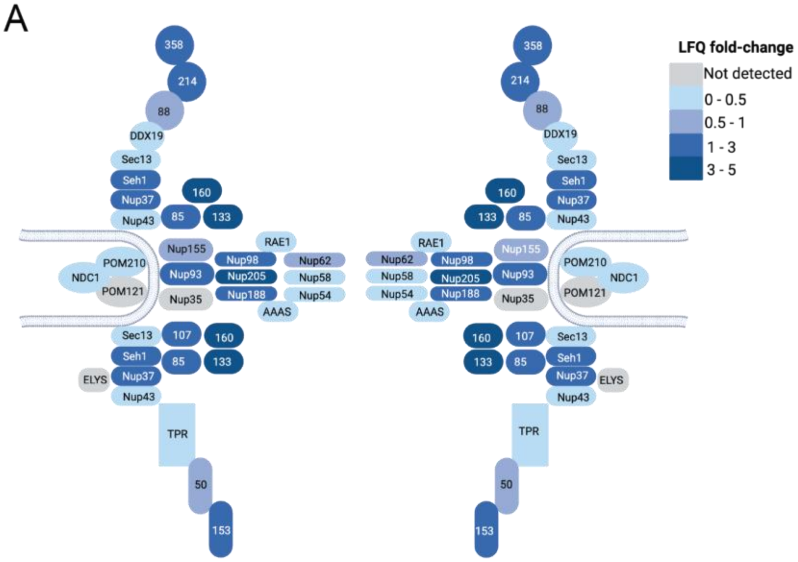


B

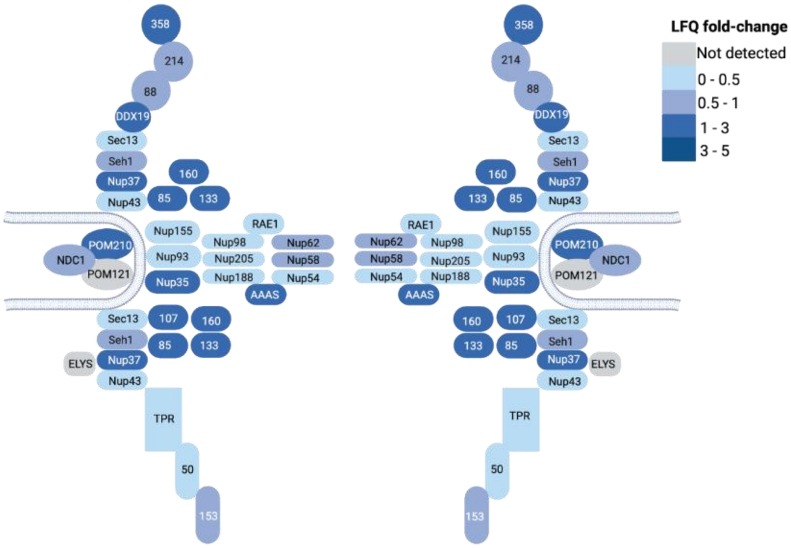


**Figure S22. Depletion of nucleoporins by JC19 and EN450 is not biased to a specific region of the NPC.** Heatmaps of the nuclear pore complex, with each nucleoporin colored based on the LFQ fold-change observed in the bulk proteome analyses in response to JC19 (A) or EN450 (B). MS data can be found in **Table S4**.

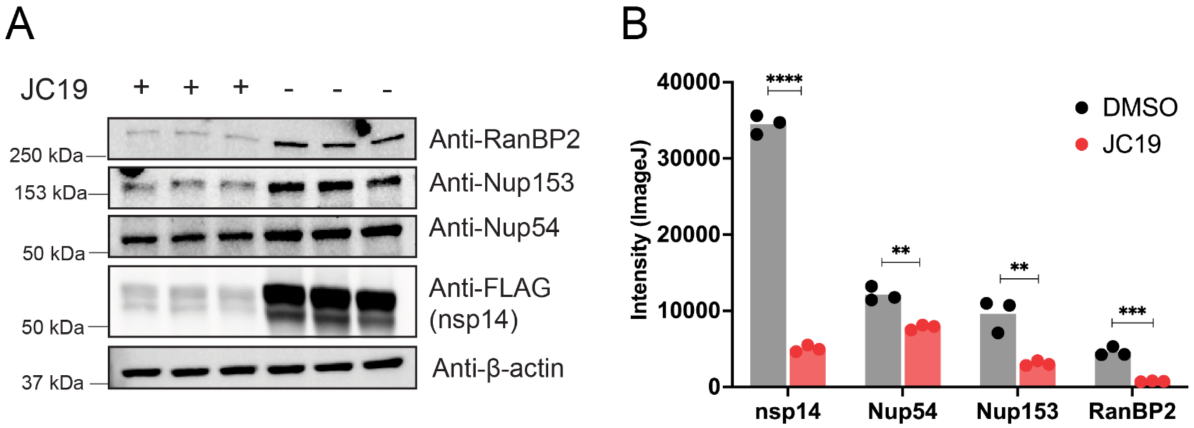


**Figure S23. Nucleoporin proteins, in addition to nsp14, are depleted by JC19.** (A) HEK293T cells transiently expressing nsp14-FLAG were treated with either DMSO or 100 µM JC19 for 1 hour (*n* = 3). Immunoblot analysis was used to assay the abundance of RANBP2, NUP153, NUP54, and nsp14. (B) Quantification of the band intensities in (A) by ImageJ [2]. A student's unpaired t-test was performed to obtain the p-values, with p-value < 0.01, < 0.001, < 0.0001 represented by **, ***, and ****, respectively.

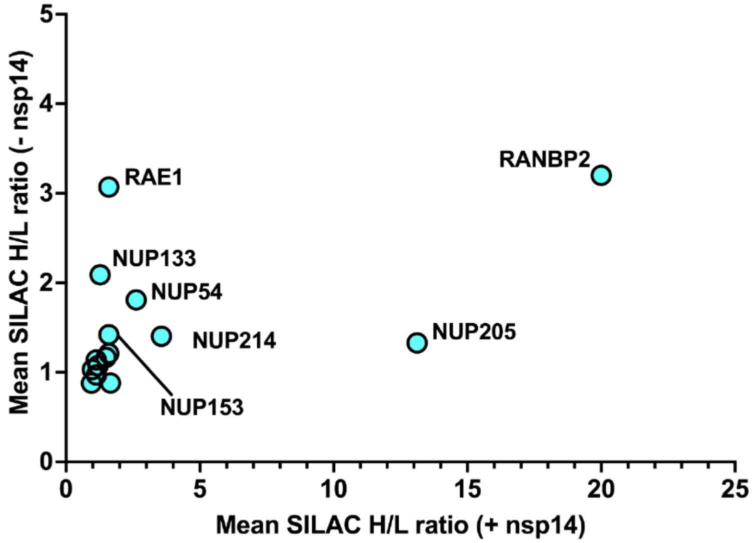


**Figure S24. Nucleoporins are depleted by cysteine-reactive electrophiles in the presence and absence of nsp14.** SILAC HEK293T cells were either mock transfected or transfected with nsp14 for 36 hours (*n* = 3 'light' and *n* = 3 'heavy' for each condition), and each set treated with either DMSO (heavy SILAC cells, *n* = 3) or 100 µM JC19 (light SILAC cells, *n* = 3) for 1 hour. Cells were then lysed, equal amounts of heavy and light soluble lysates combined, and lysates prepared for LC-MS/MS analysis via SP3 clean-up. The H/L SILAC MS1 ratio was quantified for each protein, and the average unlogged ratios for each protein in the nsp14-transfected condition was plotted against the respective ratio in the mock-transfected condition. All MS data can be found in **Table S4**.

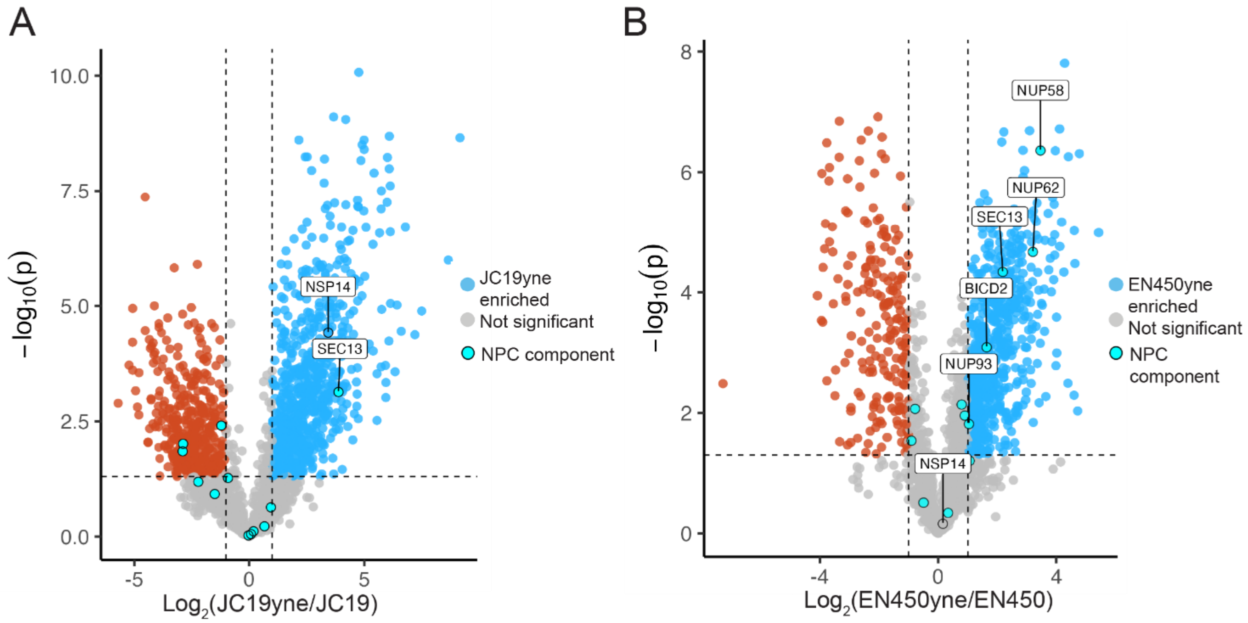


**Figure S25. Nucleoporins are largely not directly labeled by JC19 or EN450 to afford depletion.** (A) HEK293T cells transiently expressing nsp14-FLAG were treated with either 100 µM JC19 ($n$ = 6) or 100 µM JC19yne ($n$ = 6) for 20 minutes. Cells were then lysed, proteins conjugated to biotin-azide via CuAAC, enriched on streptavidin, and on-resin trypsin digest performed to release peptides for LC-MS/MS. Label-free quantification was used to measure the intensity of each protein in all replicates. A student's t-test was performed between control and treatment groups to generate a p-value, which was plotted against the average fold-change of each protein. Proteins with a $\log_2$(fold-change) > 1 and p-value < 0.05 are considered significantly enriched by JC19yne. (B) HEK293T cells transiently expressing nsp14-FLAG were treated with either 100 µM EN450 ($n$ = 4) or 100 µM EN450yne ($n$ = 4) for 4 hours. Cells were then lysed, proteins conjugated to biotin-azide via CuAAC, enriched on streptavidin, and on-resin trypsin digest performed to release peptides for LC-MS/MS. Label-free quantification was used to measure the intensity of each protein in all replicates. A student's t-test was performed between control and treatment groups to generate a p-value, which was plotted against the average fold-change of each protein. Proteins with a $\log_2$(fold-change) > 1 and p-value < 0.05 are considered significantly enriched by EN450yne. All MS data can be found in **Table S3** and **Table S4**.

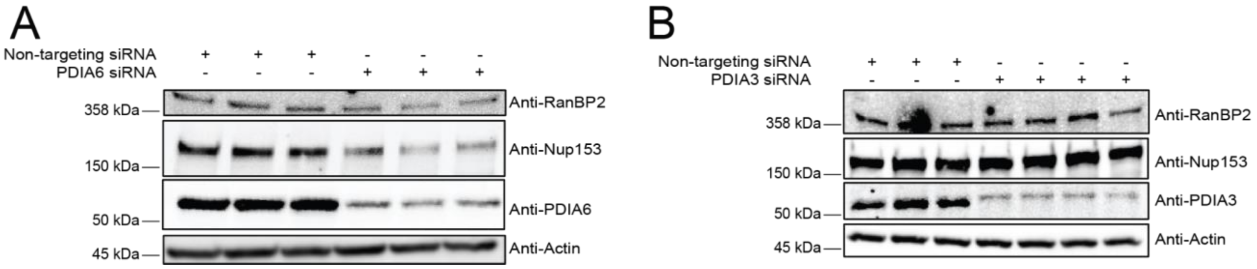


**Figure S26. Effect of PDIA6 and PDIA3 siRNA knockdowns on nucleoporins RANBP2 and NUP153. Knockdown of PDIA6 does not have a widespread effect on the abundance of nucleoporins.** (A) Western blot showing RANBP2 and NUP153 abundance in response to SiRNA knockdown PDIA6 as compared to scramble siRNA in HEK293T nsp14-overexpressing cells ($n$ = 3). (B) Western blot showing RANBP2 and NUP153 abundance in response to SiRNA knockdown PDIA3 as compared to scramble siRNA in HEK293T nsp14-overexpressing cells ($n$ = 3 or 4).

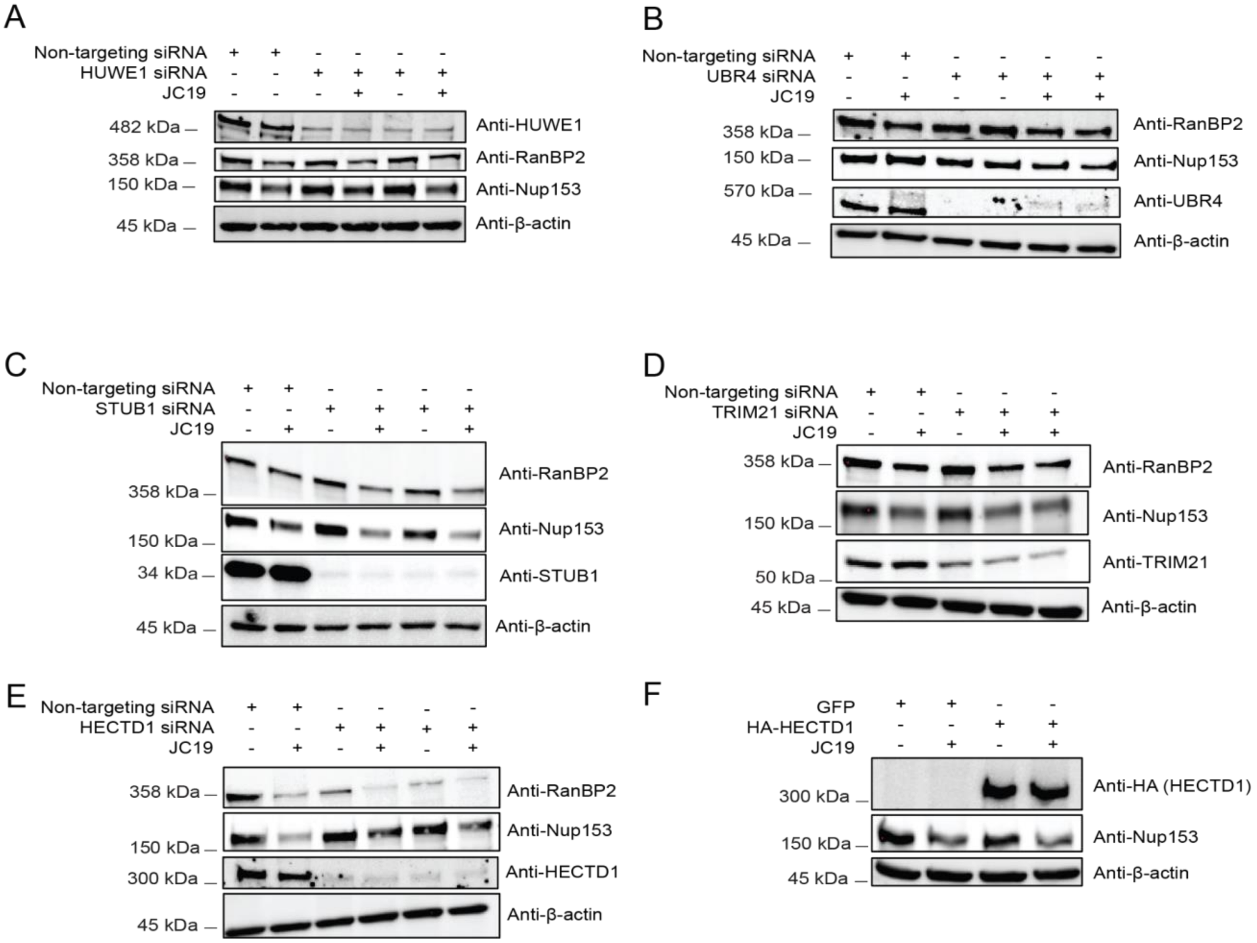


**Figure S27. Effect of E3 ligase knockdowns on nuclear pore proteins.** (A) Immunoblot of siRNA knockdown of HUWE1 and its impact on RANBP2 and NUP153 abundance, both with and without JC19 treatment (25 µM, 0.5 h). (B) Immunoblot of siRNA knockdown of UBR4 and its impact on RANBP2 and NUP153 abundance, both with and without JC19 treatment (25 µM, 0.5 h). (C) Immunoblot of siRNA knockdown of STUB1 and its impact on RANBP2 and NUP153 abundance, both with and without JC19 treatment (25 µM, 0.5 h). (D) Immunoblot of siRNA knockdown of TRIM21 and its impact on RANBP2 and NUP153 abundance, both with and without JC19 treatment (25 µM, 0.5 h). (E) Immunoblot of siRNA knockdown of HECTD1 and its impact on RANBP2 and NUP153 abundance, both with and without JC19 treatment (100 µM, 1 h). (F) HA-HECTD1 overexpression and its impact on NUP153 abundance upon JC19 treatment (100 µM, 1 h) as compared to control.

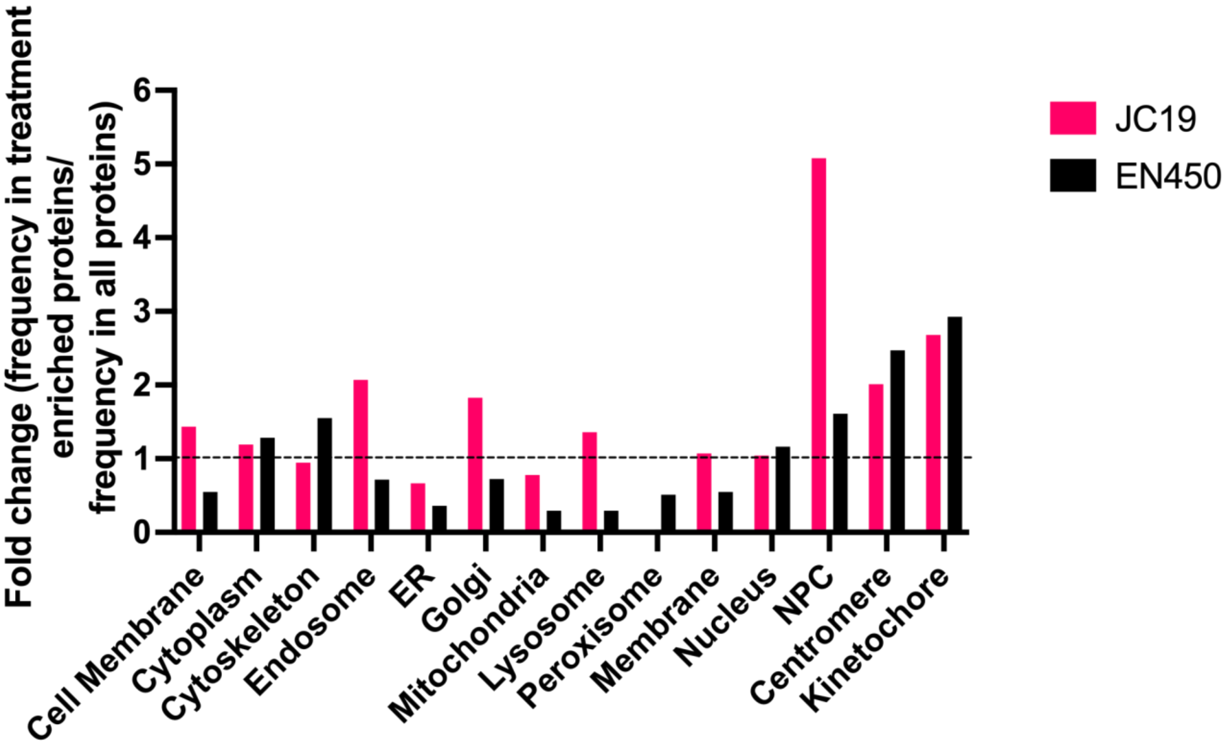
Fold change (frequency in treatment enriched proteins/ frequency in all proteins)
0
1
2
3
4
5
6
Cell Membrane
Cytoplasm
Cytoskeleton
Endosome
ER
Golgi
Mitochondria
Lysosome
Peroxisome
Membrane
Nucleus
NPC
Centromere
Kinetochore
JC19
EN450

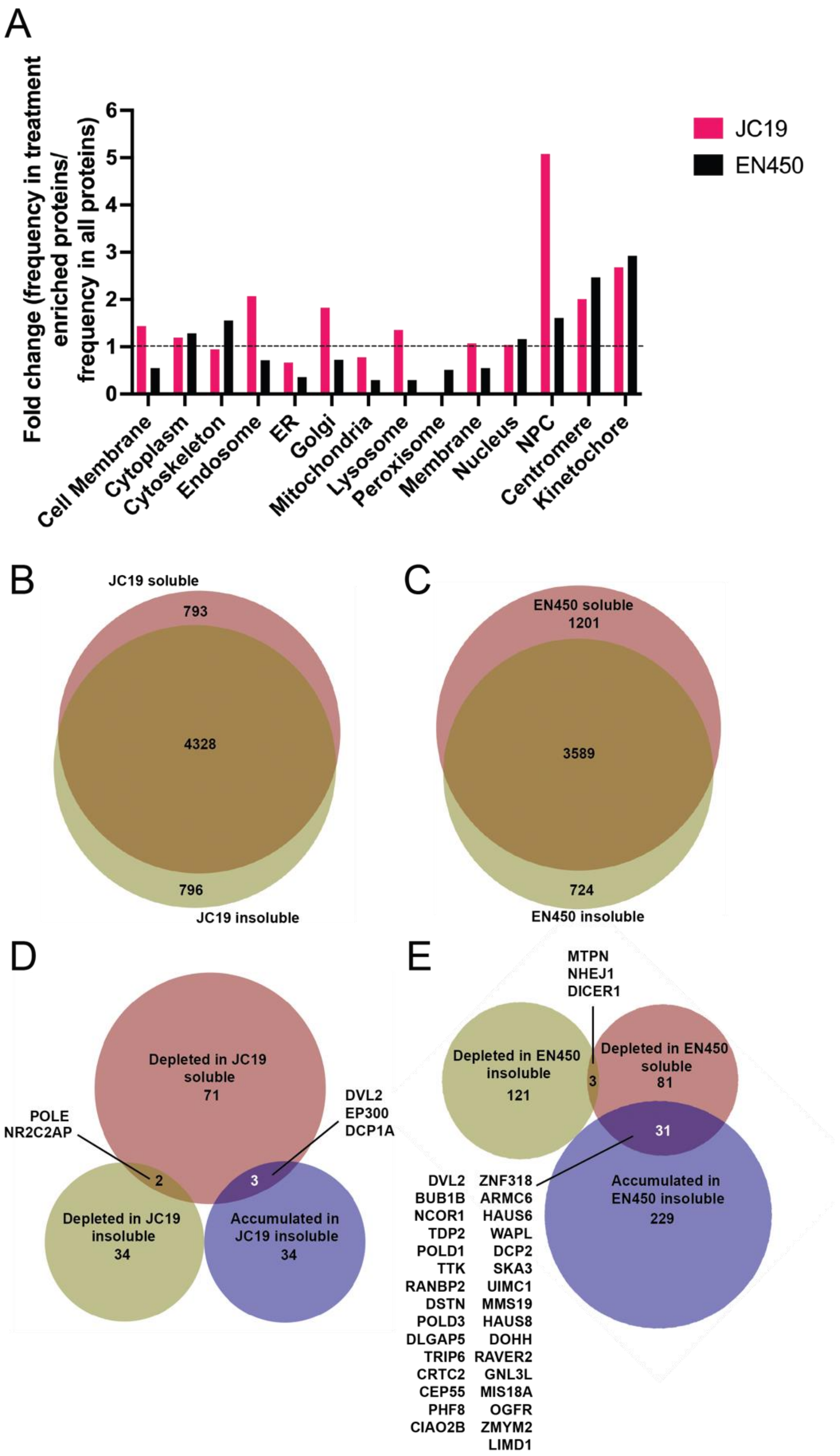


**Figure S28. The nuclear pore complex is an enriched subcellular compartment within proteins that accumulate in the insoluble fraction upon JC19 and EN450 treatment.** (A) All proteins identified from JC19 and EN450 insoluble label-free bulk proteome MS analysis annotated by subcellular location using

Uniprot annotations. The total representation of subcellular location for all proteins that were identified was compared to representation of subcellular locations for the enriched proteins. Fold-change of those two values were plotted above. (B-C) The overlap of proteins identified in the bulk soluble and insoluble proteomics datasets for JC19 (B) and EN450 (C). (D-E) Overlap of proteins that were significantly depleted from the soluble dataset and significantly depleted or accumulated in the insoluble dataset for JC19 (D) and EN450 (E). All proteins in D-E were identified in both soluble and insoluble datasets for the corresponding compound. All MS data can be found in **Table S4.**

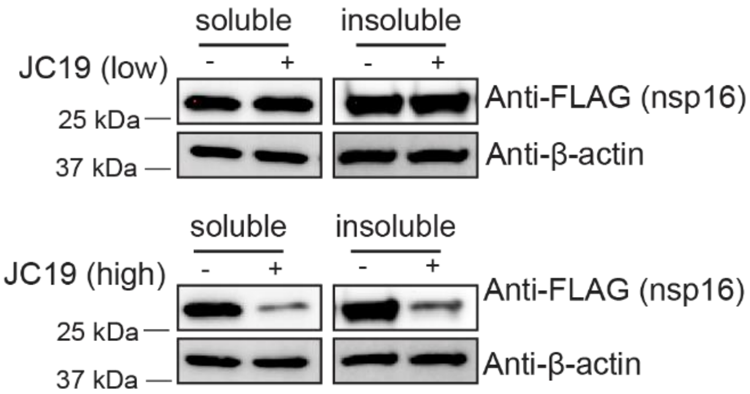


**Figure S29. Nsp16 is degraded in both the soluble and insoluble lysate fractions upon treatment with a high dose of JC19.** HEK293T cells transiently expressing nsp16-FLAG were treated with DMSO, a low dose of JC19 (25 µM for 30 minutes), or a high dose of JC19 (100 µM for 1 hour) and subject to immunoblot analysis. Cells were lysed in 0.3% CHAPS to generate the soluble lysate, and after clearance by centrifugation the insoluble debris was solubilized in 8 M urea to generate the “insoluble’ lysate.

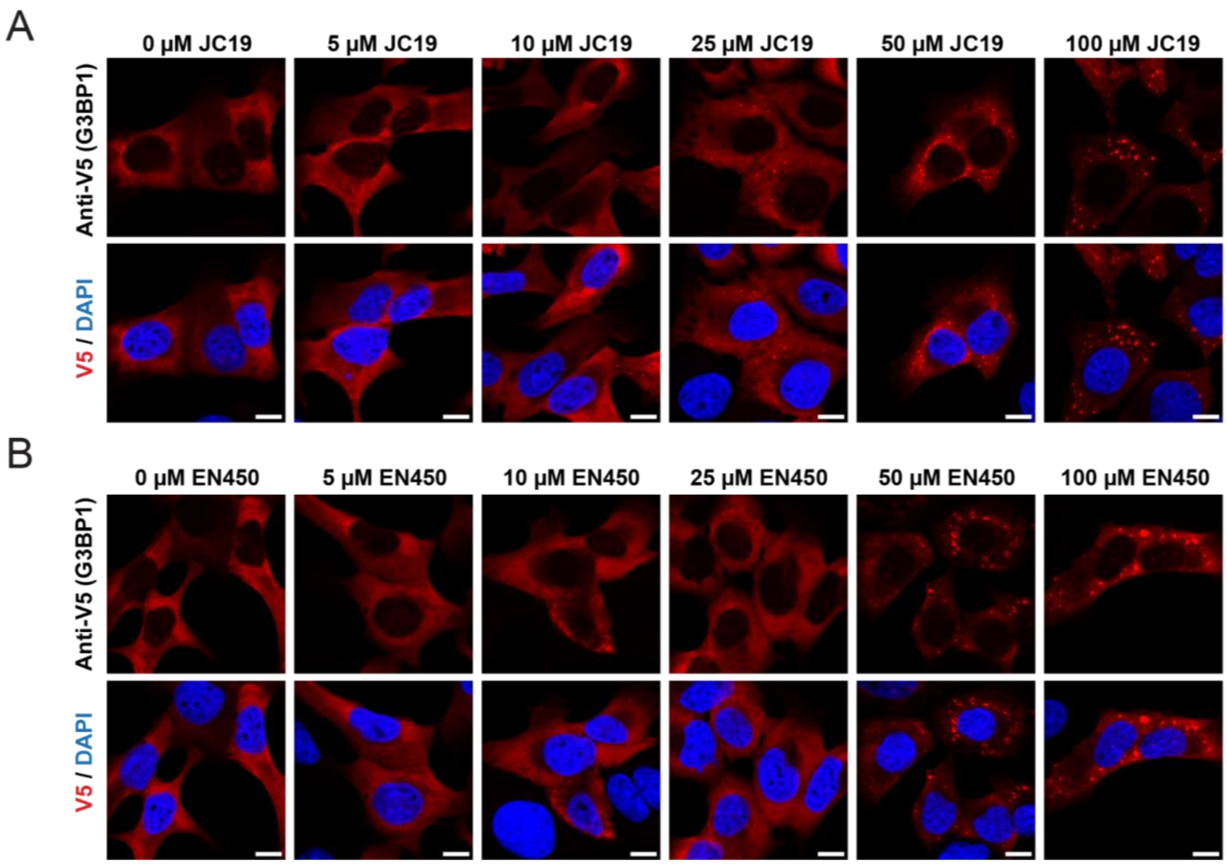


**Figure S30. Dose-dependence of cysteine-reactive electrophile induction of stress granule formation.** (A) U2OS cells stably expressing V5-tagged G3BP1 were treated with the indicated concentrations of JC19 for 30 minutes, fixed, permeabilized, and subject to immunofluorescence microscopy. Images were acquired on an LSM880 confocal microscope at 63X objective and 2X manual zoom. All scale bars = 10µm. (B) U2OS cells stably expressing V5-tagged G3BP1 were treated with the indicated concentrations of EN450 for 30 minutes, fixed, permeabilized, and subject to immunofluorescence microscopy. Images were acquired on an LSM880 confocal microscope at 63X objective and 2X manual zoom. All scale bars = 10µm.

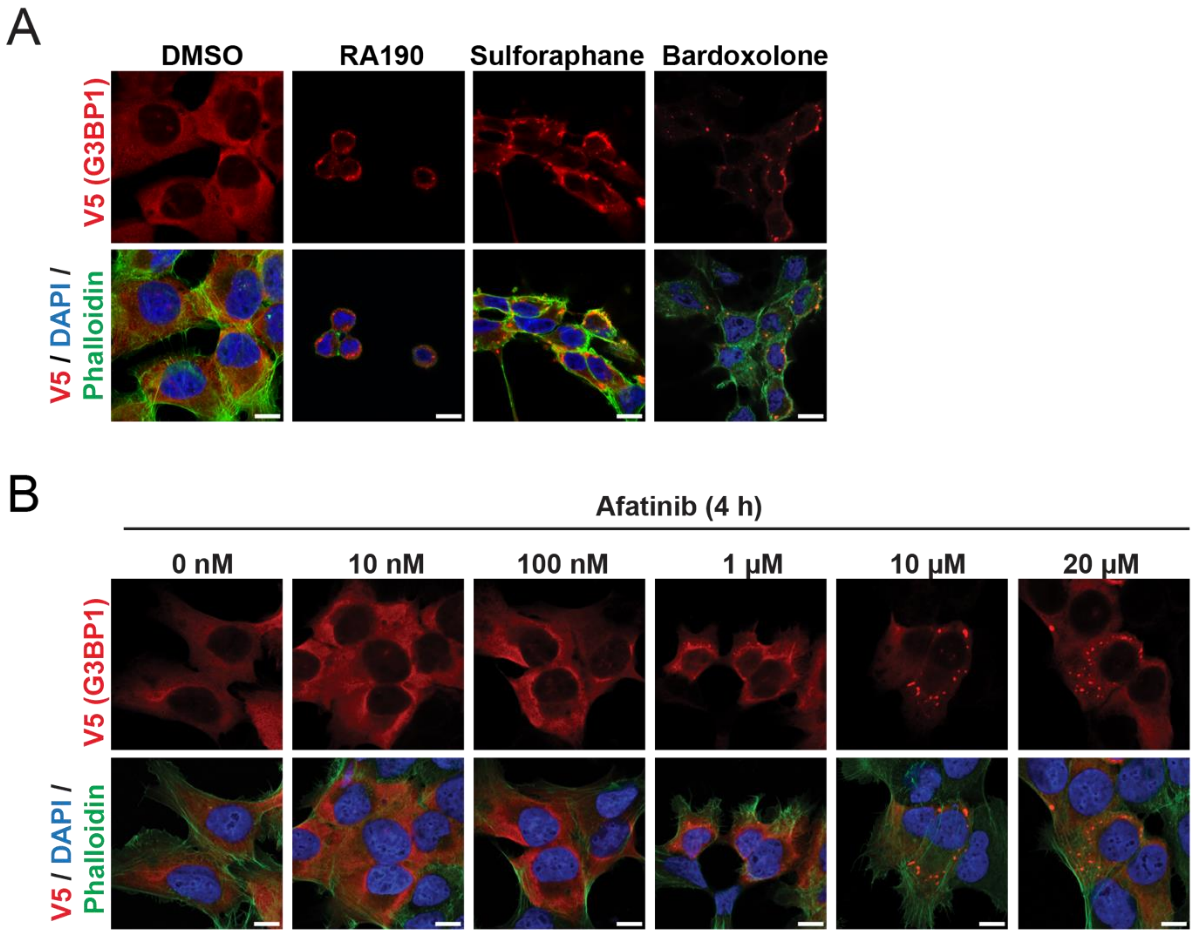


**Figure S31. Various cysteine-reactive electrophiles induce stress granule formation.** (A) U2OS cells stably expressing V5-tagged G3BP1 were treated with DMSO, RA190 (10 μM), sulforaphane (50 μM), or bardoxolone (10 μM) for 1 hour. Cells were then fixed, permeabilized, and subject to immunofluorescence microscopy. Images were acquired on an LSM880 confocal microscope at 63X objective and 2X manual zoom. All scale bars = 10μm. (B) U2OS cells stably expressing V5-tagged G3BP1 were treated with the indicated concentration of afatinib for 4 hours. Cells were then fixed, permeabilized, and subject to immunofluorescence microscopy. Images were acquired on an LSM880 confocal microscope at 63X objective and 2X manual zoom. All scale bars = 10μm.

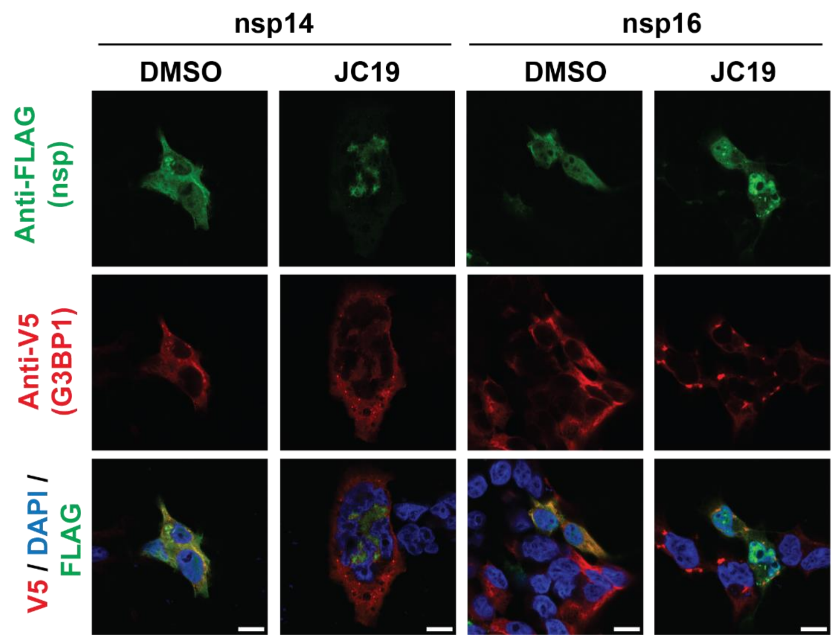


**Figure S32. Nsp14 and nsp16 are not sequestered into stress granules (SGs) in response to JC19 treatment.** HEK293T cells stably expressing G3BP1-TurboID-V5 were transiently transfected with nsp14 or nsp16. Cells were then treated with either DMSO or 100 µM JC19 for 1 hour, fixed, permeabilized, and subject to immunofluorescence microscopy to visualize nsp14, nsp16, and G3BP1. Images were acquired on an LSM880 confocal microscope at 63X objective and 2X manual zoom. All scale bars = 10 µm.

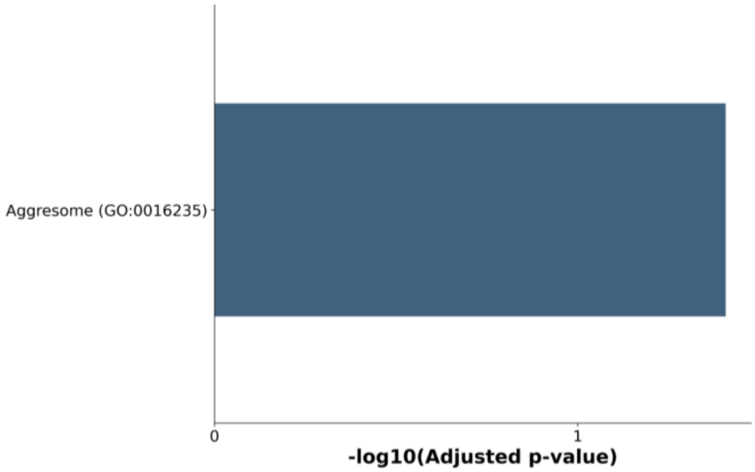


**Figure S33. The aggresome is enriched among proteins that accumulate in the insoluble fraction upon JC19 treatment.** Gene Ontology Cellular Compartment analysis (from GO Cellular Compartment 2023) on the proteins that significantly accumulated in the insoluble fraction upon JC19 treatment ($log_2$(fold-change) > 1 and p-value < 0.05 relative to DMSO control) relative to all proteins identified in the dataset. MS Data can be found in **Table S5**.

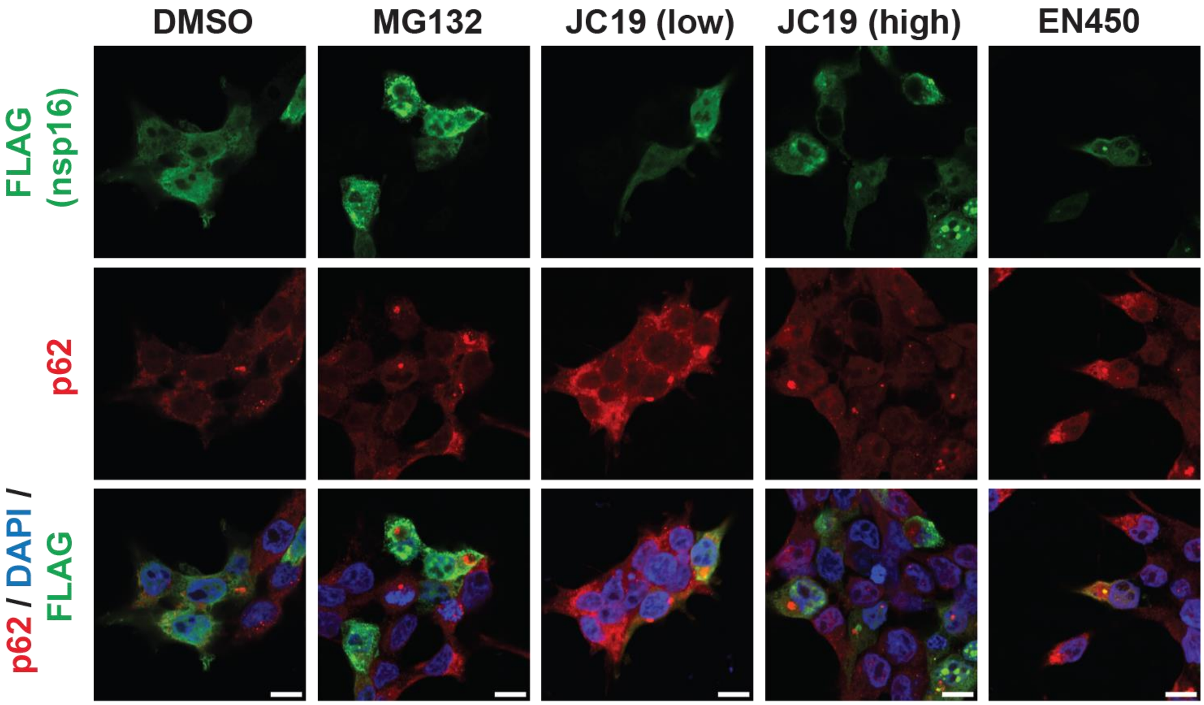


| | DMSO | MG132 | JC19 (low) | JC19 (high) | EN450 |
|---|---|---|---|---|---|
| Pearson's Coefficient | **0.53** | **0.32** | **0.45** | **0.58** | **0.85** |
| Mander's Coefficient | **0.436** | **0.525** | **0.837** | **0.865** | **0.632** |
| Yellow % (yellow area / green area) | **6.22%** | **3.94%** | **24.74%** | **30.78%** | **31.07%** |

**Figure S34. nsp16 is recruited to the aggresome upon compound treatment.** HEK293T cells were transiently transfected with nsp16-FLAG, treated with DMSO, MG132 (10 µM, 5 hours), a low dose of JC19 (25 µM, 30 minutes), a high dose of JC19 (100 µM, 1 hour), or EN450 (100 µM, 1 hour), fixed, permeabilized, and subject to immunofluorescence microscopy to visualize nsp16 and p62. Images were acquired on an LSM880 confocal microscope at 63X objective and 2X manual zoom. All scale bars = 10 µm.

#### (B) Supplementary Tables

**Table S1-5.** Datasets corresponding to each figure, provided in the attached supplementary files.

Table S1: Figure 1
Table S2: Figure 3
Table S3: Figure 4
Table S4: Figure 5
Table S5: Figure 6
Table S11: Files in Proteomics Identification Database (PRIDE) datasets

**Table S6**. Antibodies used in this study

| Name | Catalog number | Lot number | Concentration |
|---|---|---|---|
| IRDye® 800CW Goat anti-Rabbit Secondary Antibody | LI-COR (926-32211) | D20803-10 | 1:5000 (IB) |
| IRDye® 800CW Goat anti-Mouse Secondary Antibody | LI-COR (926-32210) | D1116-25 | 1:5000 (IB) |
| IRDye 680RD Donkey anti-Rabbit Secondary Antibody | LI-COR (926-68073) | D00421-09 | 1:5000 (IB) |
| IRDye 680RD Donkey anti-Mouse Secondary Antibody | LI-COR (926-68072) | D20322-25 | 1:5000 (IB) |
| Alexa Fluor™ 546 donkey anti-mouse IgG (H+L) | Invitrogen (A10036) | 2160040 | 1:1000 (IF) |
| Alexa Fluor™ 633 goat anti-rabbit IgG (H+L) | Invitrogen (A21070) | 2563844 | 1:1000 (IF) |
| Donkey anti-Rabbit IgG (H+L) Alexa Fluor™ Plus 488 | Invitrogen (A32790) | VC296619 | 1:1000 (IF) |
| DYKDDDDK Tag (D6W5B) Rabbit mAb | Cell signaling (14793S) | 7 | 1:3000 (IB); 1:1000 (IF) |
| β-Actin (8H10D10) Mouse mAb | Cell signaling (3700S) | 21 | 1:3000 |
| GAPDH (14C10) Rabbit mAb | Cell signaling (2118S) | 16 | 1:3000 |
| HA-Tag (C29F4) Rabbit mAb | Cell signaling (3724S) | 10 | 1:3000 |
| PSMB5 (D1H6B) Rabbit mAb #12919 | Cell signaling (12919S) | 1 | 1:3000 |
| 6x-His Tag Monoclonal Antibody (HIS.H8) | ThermoFisher (MA1-21315) | TK269015 | 1:3000 |

| | | | |
|---|---|---|---|
| PDI (C81H6) Rabbit mAb | Cell Signaling (3501S) | 3 | 1:3000 |
| PDIA6 Polyclonal antibody | Proteintech (18233-1-AP) | 00055732 | 1:3000 |
| PDIA6 Monoclonal antibody | Proteintech (66669-1-Ig) | | 1:3000 |
| ERp57 Rabbit pAb | Abclonal (A1085) | 0056780201 | 1:3000 |
| ERp72 Antibody | Cell Signaling (2798S) | 2 | 1:3000 |
| LC3A/B (D3U4C) XP® Rabbit mAb | Cell Signaling (12741S) | 5 | 1:2000 |
| PARP (46D11) Rabbit mAb | Cell Signaling (9532S) | 10 | 1:3000 |
| NRF2 Rabbit pAb | Abclonal (NO:A1244) | 5500014741 | 1:3000 |
| BiP (C50B12) Rabbit mAb #3177 | Cell Signaling (3177S) | 10 | 1:1000 |
| CHOP (L63F7) Mouse mAb | Cell Signaling (2895S) | 13 | 1:1000 |
| ATF-4 (D4B8) Rabbit mAb | Cell Signaling (11815S) | 5 | 1:1000 |
| STUB1 Rabbit pAb | ABclonal (A11751) | 0071170101 | 1:3000 |
| TRIM21/SS-A Rabbit mAb | ABclonal (A22724) | 6100001135 | 1:3000 |
| HUWE1 Rabbit pAb | ABclonal (A9433) | 5500008129 | 1:3000 |
| UBR4 Rabbit pAb | ABclonal (A12193) | 0092800201 | 1:3000 |
| HECTD1 Rabbit pAb | ABclonal (A9433) | 0101520101 | 1:3000 |
| RANBP2 (D-4) | Santa Cruz | I0821 | 1:1000 |

| | | | |
|---|---|---|---|
| | Biotech (sc-74518) | | |
| NUP153 (E3I6Z) | Cell Signaling (98559S) | 1 | 1:3000 |
| NUP54 Rabbit PolyAb | Proteintech (16232-1-AP) | 00059775 | 1:3000 |
| NFKB1 Rabbit Ab | Cell Signaling (3035S) | 6 | 1:3000 |
| V5-Tag (E9H80) Mouse mAb | Cell Signaling (80076S) | 3 | 1:3000 (IB); 1:1000 (IF) |
| Alexa Fluor$^{TM}$ 488 phalloidin | Invitrogen (A12379) | 2282104 | 1:800 (IF) |
| PSMB2 Rabbit PolyAb | Proteintech (I5I54-I-AP) | | 1:300 (IF) |
| Rb pAn to RanBP2 | Abcam (AB64276) | 1024132-3 | 1:300 (IF) |
| SQSTM1 (D-3) | Santa Cruz Biotech (sc-28359) | G2822 | 1:100 (IF) |

**Table S7. siRNAs used in this study**

| Target Gene | Manufacturer | Catalog Number | Lot # | Duplex Sequence |
|---|---|---|---|---|
| HECTD1 | IDT | hs.Ri.HECTD1.13.1 | 538907239 | 5'-AUGGUCAGCAAGCUACAAUAUUGAA-3'; 5'-UUCAAUAUUGUAGCUUGCUGACCAUCU-3' |
| HECTD1 | IDT | hs.Ri.HECTD1.13.2 | 538907240 | 5'-ACUGAUGCAAGCUAUCCAUCAGUCA-3'; 5'-UGACUGAUGGAUAGCUUGCAUCAGUAG-3' |
| HUWE1 | ambion | s19595 | ASO2JTF1 | 5'-CAUUGGAAAGUGCGAGUUAtt-3'; 5'-UAACUCGCACUUUCCAAU-3' |

| | | | | |
|---|---|---|---|---|
| Negative Control siRNA | IDT | 51-01-14-03 | 663345 | 5'-CGUUAAUCGCGUAUAAUACGCGUAT-3'; 5'-AUACGCGUAUUAUACGCGAUUAACGAC-3' |
| PDIA3 | ambion | s6229 | ASO2KV80 | 5'-GAACUUGAGGGAUAACUAtt-3'; 5'-UAGUUAUCCCUCAAGUUGCt-3' |
| PDIA3 | ambion | s6227 | ASO2KV8R | 5'-GGAAUAGUCCCAUUAGCAAtt-3'; 5'-UUGUAAUGGGACUAUUCCt-3' |
| PDIA4 | IDT | hs.Ri.PDIA4.13.1 | 445103958 | 5'-GGUGUUUCACUUCAUGCUCUGAATA-3'; 5'-UAUUCAGAGCAUGAAGUGAAACACCAA-3' |
| PDIA4 | IDT | hs.Ri.PDIA4.13.2 | 445103959 | 5'-ACCUGAGAGAAGAUUACAAAUUUCA-3'; 5'-UGAAAUUUGUAAUCUUCUCUCAGGUUG-3' |
| PDIA6 | IDT | hs.Ri.PDIA6.13.1 | 455645784 | 5'-GGAUGCUACAGUCAAUCAGGUUCTG-3'; 5'-CAGAACCUGAUUGACUGUAGCAUCCAC-3' |
| PDIA6 | IDT | hs.Ri.PDIA6.13.2 | 455645785 | 5'-AGGCUUUCUAUGUUGUUAAUGAACC-3'; 5'-GGUUCAUUAACAACAUAGAAAGCCUUG-3' |
| PH4B | IDT | hs.Ri.P4HB.13.1 | 448986282 | 5'-AUUUCUAUUUCACAAUCGAAUUGAA-3'; 5'-UUCAAUUCGAUUGUGAAAUAGAAAUGC-3' |
| PH4B | IDT | hs.Ri.P4HB.13.2 | 448986283 | 5'-AGAUGAACUGUAAUACGCAAAGCCA-3'; 5'-UGGCUUUGCGUAUUACAGUUCAUCUUU-3' |
| STUB1 | IDT | hs.Ri.STUB1.13.1 | 544882524 | 5'-AGGUUAUUGACGCAUUCAUCUCUGA-3'; 5'-UCAGAGAUGAAUGCGUCAAUAACCUCC-3' |
| STUB1 | IDT | hs.Ri.STUB1.13.2 | 544882525 | 5'-GACGAGCUUUUUUCUCAGGUGGATG-3'; 5'-CAUCCACCUGAGAAAAAAGCUCGUCCA-3' |

| | | | | |
|---|---|---|---|---|
| TRIM21 | IDT | hs.Ri.TRIM21.13.2 | 544882526 | 5'-GUCCACUGAAUAUUGGAUCACAAGG-3'; 5'-CCUUGUGAUCCAAUAUUCAGUGGACAG-3' |
| UBR4 | IDT | hs.Ri.UBR4.13.1 | 530799933 | 5'-GUCAACUGGAUAAAGGAUCACCUTA-3'; 5'-UAAGGUGAUCCUUUAUCCAGUUGACAU-3' |
| UBR4 | IDT | hs.Ri.UBR4.13.2 | 530799934 | 5'-GAGAAGCAGUACGAGCCAUUCCUACT-3'; 5'-AGUAGAAUGGCUCGUACUGCUUCUCAU-3' |

**Table S8.** Primers used in this study

| Gene | Forward primer | Reverse primer |
|---|---|---|
| nsp14-FLAG Cys11Ala | 5'-GAGAACGTGACCGGTCTGTTCAAGGACGCCTCTAAGGTCATCACTGGTCTGCACC-3' | 5'-GAGCCTGAGTTGGGTGCAGACCAGTGATGACCTTAGAGGCGTCCTTGAACAGACCGG-3' |
| nsp14-FLAG Cys39Ala | 5'-CCAAGTTCAAGACCGAAGGACTGGCCGTGGACATCCCCGGTATCCCTAAG-3' | 5'-GTAGGTCATGTCCTTAGGGATACCGGGGATGTCCACGGCCAGTCCTTCGG-3' |
| nsp14-FLAG Cys94Ala | 5'-GATCGGTTTCGACGTGGAAGGTGCCCACGCTACCAGAGAAGCTGTGGG-3' | 5'-GCAGGTTGGTACCCACAGCTTCTCTGGTAGCGTGGGCACCTTCCACGTC-3' |
| nsp14-FLAG Cys208Ala | 5'-GTGAAGATCGGACCCGAACGTACCTGCGCCCTGTGCGACAGACGTGCC-3' | 5'-GAGAAGCAGGTGGCACGTCTGTCGCACAGGGCGCAGGTACGTTCGGG-3' |
| nsp14-FLAG Cys208Val | 5'-GTGAAGATCGGACCCGAACGTACCTGCGCCCTGTGCGACAGACGTGCC-3' | 5'-GAGAAGCAGGTGGCACGTCTGTCGCACAGGGCGCAGGTACGTTCGGG-3' |
| nsp14-FLAG Cys309Ala | 5'-CAACGCCGCTGCCAGAAAGGTGCAGCACATGGTG-3' | 5'-GCTGCACCTTTCTGGCAGCGGCGTTGATCTTCAGTT-3' |
| nsp14-FLAG Cys356Ala | 5'-GGAGTGGAAGTTCTACGACGCTCAGCCCGCCTCCGACAAG | 5'-GTTCCTCGATCTTGTAGGCCTTGTCGGAGGCGGGCTGAGCG |

| | | |
|---|---|---|
| | GCCTAC-3’ | TAG-3’ |
| nsp14-FLAG Cys414Ala | 5'-CCTGCCCGGAGCCGACGGCGGATCCCTGTACGTGAAC-3' | 5'-CGCCGTCGGCTCCGGGCAGGTTCAGGTTGCTCAGCACACG-3' |
| nsp14-FLAG Cys216Ala | 5'-GACGTGCCACCGCCTTCTCCACCGCCTCTGACACC-3' | 5'-GGCGGTGGAGAAGGCGGTGGCACGTCTGTCGCACAGG-3' |
| nsp14-FLAG Cys272Ala | 5'-CCACGTGGCTTCTGCCGACGCTATCATGACCAGATGCCTG-3' | 5'-GTCATGATAGCGTCGGCAGAAGCCACGTGGGCGTTACCG-3' |
| nsp14-FLAG Cys279Ala | 5'-CATGACCAGAGCCCTGGCCGTGCACGAGTGC-3' | 5'-CGTGCACGGCCAGGGCTCTGGTCATGATAGCGTCGC-3' |
| nsp14-FLAG Cys340Ala | 5'-GCT ATC AAG GCC GTG CCC CAG GCC GAC GTG G-3' | 5'-GGG GCA CGG CCT TGA TAG CCT TTG GGT TTC CG-3' |
| G3BP1 (for cloning into plasmid #15) | 5’-TAAGGGAAGCTTGGGCCGCCACCATGGTGATGGAGAAGCCT-3’ | 5’-TGCTTAGCGGCCCTGCCGTGGCGCAAGCC-3’ |
| G3BP1-TurboID-V5 (for cloning into plasmid #18) | 5’-GAAGGGGGATCCCGCCACCATGGTGATGGAGAAGCCTAG-3’ | 5’-GGCGTGATCGATTTAGGTGCTGTCCAGGCCCAGCAG-3’ |
| G3BP1-TurboID-V5 (for cloning into plasmid #20) | 5’-GAAGGGTCTAGACGCCACCATGGTGATGGAGAAGCCT-3’ | 5’-GTGCGGTGTACATTAGGTGCTGTCCAGGCCCAG-3’ |

**Table S9**. Plasmids used in this study

| **Plasmid number** | **Figure** | **Plasmid** | **Source** | **Resistance** |
|---|---|---|---|---|
| 1 | 1 | pDONR223 SARS-CoV-2 NSP14_nostop | Addgene (#149316) | Spectinomycin |
| 2 | 1 | pDONR223 SARS-CoV-2 NSP9_nostop | Addgene (#149312) | Spectinomycin |

| | | | | |
|---|---|---|---|---|
| 3 | 1 | pDONR223 SARS-CoV-2 NSP10_nostop | Addgene (#149313) | Spectinomycin |
| 4 | 1 | pDONR223 SARS-CoV-2 NSP16_nostop | Addgene (#149318) | Spectinomycin |
| 5 | 1 | GateWay compatible C-terminal FLAG destination vector | Gift from T Wucherpfennig | Chloramphenicol and ampicillin |
| 6 | 1-6 | nsp14-FLAG expression vector | GateWay cloned with plasmid #1 (donor) and #5 (destination) | Ampicillin |
| 7 | 1 | nsp9-FLAG expression vector | GateWay cloned with plasmid #2 (donor) and #5 (destination) | Ampicillin |
| 8 | 1 | nsp10-FLAG expression vector | GateWay cloned with plasmid #3 (donor) and #5 (destination) | Ampicillin |
| 9 | 1, 6 | nsp16-FLAG expression vector | GateWay cloned with plasmid #4 (donor) and #5 (destination) | Ampicillin |
| 10 | 3 | pRK5-HA-Ubiquitin-WT | Addgene (#17608) | Ampicillin |
| 11 | 3 | Nsp14_K_R ENTR (pMB1) | Twist Biosciences (custom sequence) | Kanamycin |
| 12 | 3 | Nsp14_K_R expression vector | GateWay cloned with plasmid #11 | Ampicillin |

| | | | | |
|---|---|---|---|---|
| | | | (donor) and #5 (destination) | |
| 13 | 4 | PDIA3-bio-His | Addgene (#52070) | Ampicillin |
| 14 | 6 | pENTR4_G3BP1 | Addgene (#127104) | Kanamycin |
| 15 | 6 | C1(1-29)-TurboID-V5_pCDNA3 | Addgene (#107173) | Ampicillin |
| 16 | 6 | G3BP1-TurboID-V5 | Restriction cloned G3BP1 from plasmid #14 into plasmid #15 | Ampicillin |
| 17 | S27 | HA-HECTD1 | Gift from Dr. Irene Zohn at Children's National | Ampicillin |
| 18 | 6 | Empty Piggybac | Gift from Melody Li at UCLA | Ampicillin |
| 19 | 6 | Piggybac transposase | Gift from Melody Li at UCLA | Ampicillin |
| 20 | S32 | pLenti CMV GFP Hygro (656-4) | Addgene (#17446) | Ampicillin |
| 21 | 6 | G3BP1-TurboID-V5_Piggybac | Restriction cloned G3BP1 from plasmid #16 into plasmid #18 | Ampicillin |
| 22 | 6 | G3BP1-TurboID-V5_pCMVHygro | Restriction cloned G3BP1 from plasmid #16 into plasmid #20 | Ampicillin |

**Table S10.** Conditions of Liquid-chromatography (LC)

| Experiment Type | Parameter | Condition |
| --- | --- | --- |
| isoTOP-ABPP<br>-yne enrichment<br>Competition<br>SILAC<br>XRNAX | Column | 100 μM ID fused silica capillary packed in-house with bulk C18 reversed phase resin (particle size, 1.9 μm; pore size, 100 Å; Dr. Maisch GmbH) |
| | Mobile phase | Buffer A: water with 3% DMSO and 0.1% formic acid<br>Buffer B: 80% acetonitrile with 3% DMSO and 0.1% formic acid |
| | Gradient and flow rate | 0 – 5 min, 3 – 20% B, 220 nL/min |
| | | 5 – 64 min, 20 – 47% B, 220 nL/min |
| | | 64 – 70 min, 47 – 95% B, 250 nL/min |
| | Run time | 70 minutes |
| | Injection volume | 5 uL |
| EN450 LFQ<br>Insoluble LFQ | Column | 100 μM ID fused silica capillary packed in-house with bulk C18 reversed phase resin (particle size, 1.9 μm; pore size, 100 Å; Dr. Maisch GmbH) |
| | Mobile phase | Buffer A: water with 3% DMSO and 0.1% formic acid<br>Buffer B: 80% acetonitrile with 3% DMSO and 0.1% formic acid |
| | Gradient and flow rate | 0 – 3 min, 1 – 6% B, 220 nL/min |
| | | 3 – 133 min, 6 - 35% B, 220 nL/min |
| | | 133 – 153 min, 35 - 40% B, 220 nL/min |

| | | |
|---|---|---|
| | | 153 – 160 min, 40 – 95% B, 250 nL/min |
| | Run time | 160 minutes |

### (C) Biology Methods

**Cell lines, culture conditions.** Cell culture reagents including Dulbecco's phosphate-buffered saline (DPBS), Dulbecco's Modified Eagle Medium (DMEM) media, and penicillin/streptomycin (Pen/Strep) were purchased from Fisher Scientific. Fetal Bovine Serum (FBS) was purchased from Avantor Seradigm (lot #214B17). All cell lines were obtained from ATCC and were maintained at a low passage number (<20 passages). U2OS cells were provided by Dr. Melody Li and used author's with permission. HEK293T (ATCC: CRL-3216), HeLa (ATCC: CCL-2), and U2OS cells were cultured in DMEM supplemented with 10 % FBS and 1 % antibiotics (Penn/Strep, 100 U/mL). Stable-Isotope Labeling by Amino Acids in Cell Culture (SILAC) cells were maintained in DMEM for SILAC supplemented with 10% dialyzed FBS, 1 % antibiotics (Penn/Strep, 100 U/mL), and isotopically labeled arginine and lysine. All media was filtered (0.22 µm) prior to use. Cells were maintained in a humidified incubator at 37 °C with 5 % CO2. Cell lines were tested monthly for mycoplasma using the Mycoplasma Detection Kit (InvivoGen).

**Cloning of plasmids.** List of plasmids with detailed information used in this study can be found in Table S9. pDONR223 SARS-CoV-2 vectors containing sequences of nsp9, nsp10, nsp14, and nsp16 were gifts from Fritz Roth (Addgene plasmid #149312, 149313, 149316, 149318, respectively)[3] and subcloned using GateWay cloning into C-terminal FLAG destination vector, which was a kind gift from T Wucherpfennig**.** For GateWay Cloning, 300 ng of both the donor and destination vector were combined in an microcentrifuge tube, following addition of TE buffer up to 8 µL. 2 µL of Gateway LR Clonase II enzyme mix (Invitrogen; 11791020) was added to the mixture, vortexed briefly, and allowed to incubate for 2 hours at room temperature before being transferred to 4°C for 12-16 hours. The enzymes were then inactivated at 65°C for 15 minutes, and the reaction mixture transformed into competent TOP10 cells, colonies grown up in LB media containing 100 µg/mL ampicillin, and successful cloning confirmed by sequencing. Plasmid for stable overexpression of the G3BP1-TurboID-V5 construct was generated via two rounds of restriction cloning. First, the G3BP1 sequence was PCR amplified from pENTR4_G3BP1 (a gift from Thomas Tuschl, Addgene plasmid #127104) using Phusion polymerase (Berkeley) and restriction cloned via HindIII and NotI restriction sites into C1(1-29)-TurboID-V5_pCDNA3 (a gift from Alice Ting, Addgene plasmid #107173)[4] using T4 DNA ligase. Then, the G3BP1-TurboID-V5 sequence was PCR amplified from this construct using Phusion polymerase (Berkeley) and restriction cloned into both the empty Piggybac transposon vector (a kind gift from Dr. Melody Li's lab) and the pLenti CMV GFP Hygro vector (a gift from Eric Campeau and Paul Kaufman) (Addgene plasmid #17446)[5] for generation of constructs that can be used for doxycycline-inducible stable expression or constitutive stable expression, respectively. HA-HECTD1 in pCMV plasmid was gifted by Irene Zohn at Children's National.

**Generation of U2OS cell line stably expressing doxycycline-inducible G3BP1-TurboID-V5.** U2OS cells were transfected at 70-80% confluency. For transfection of a single well of a 6-well plate, plasmid (0.5 ug of G3BP1-TurboID-V5_Piggybac transposon vector + 0.5ug of the ePiggybac transposase vector), reduced-serum OptiMEM (up to 100uL) and Lipofexin (Lambda Biotech, TS310) (3 µL) were mixed and incubated for 15 min at room temperature, followed by adding dropwise to the cells. 6-8 hours after transfection, media was replaced with fresh culture media and cells were incubated for 48 hours. After 48 hours, media was replaced with fresh media containing 1 µg/mL of puromycin (Fisher, AAJ672368EQ) for selection of stably transfected cells. Selection media was replaced every 24-48 hours until colonies of transfected cells were visible. Cells were expanded to larger plates and cryogenically frozen in FBS containing 10% DMSO for future use.

**Generation of HEK293T cell line stably expressing G3BP1-TurboID-V5.** For preparation of lentiviruses, HEK293T cells in 10 cm plates were transfected at ~70-80% confluency with lentiviral vector pLenti_CMV_GFP_Hygro (656-4) containing G3BP1-TurboID-V5 (10 µg; Addgene #17446) with the lentiviral packaging plasmids pVSVG (4 µg; Addgene #8454) and Δ8.9 (8 µg; Addgene #2221) and 66 µL of Turbo DNAfectin3000 (Lamda Biotech, TS310) in antibiotic-free media for 6 hours. After 6 hours, the DNAfectin-containing media was replaced with fresh antibiotic-free media and the cells were left to incubate for 48 hours for lentiviral generation. The media was collected (containing virus) and stored at 4°C and cells were incubated for another 24 hours in fresh media. After 24 hours, the lentivirus-containing media was collected, and added to the previously harvested media. 1/3 volume of Lenti-X concentrator (Takara Bio, Cat# 631231) was added to the total harvested media and incubated 16 hours at 4°C. The lentivirus was pelleted at 1500 *g* for 45 minutes at 4°C and resuspended in 500 µL plain DMEM and stored in 100 µL aliquots at -80°C. 300 µL of reconstituted virus was added dropwise to HEK293T cells at 70-80% confluency in antibiotic-free media containing 8 µg/mL polybrene and allowed to incubate for 48 hours. Media was then replaced with DMEM containing 400 µg/mL hygromycin and allowed to incubate for 48 hours for selection of transduced cells. Selection media was replaced every 24-48 hours until the appearance of visible colonies of transduced cells. Cells were expanded to larger plates and cryogenically frozen in FBS containing 10% DMSO for future use.

**Mutagenesis.** Point mutations (C11A, C39A, C94A, C208A, C208V, C216A, C272A, C279A, C309A, C356A, C414A, C340A) were created by PCR-based site-directed mutagenesis as previously described[6]. All primers used to generate mutant constructs can be found in Table S8. Mutant plasmids were then transformed into competent TOP10 cells, colonies selected and grown in SOC media at 37°C for >2 hours, and successful mutagenesis confirmed by sequencing.

**Transfections.** Cells were transfected at 70-80% confluency. For a 10-cm plate, plasmid (5 µg), serum-free DMEM (350 µL) and PEI MAX- Transfection Grade Linear

Polyethylenimine Hydrochloride (MW 40,000) (Polysciences, Inc., 24765-1) (35 µL of 1 mg/mL, pH 7.4) were mixed and incubated for 15 minutes at room temperature. Transfection cocktail was then added dropwise to the cells and incubated for ~36 hours. Transfection reagents were scaled according to volume of media in plates of various sizes.

**Cell treatments.** All compounds were made up as 50 mM stock solutions in DMSO. For cells transiently expressing a protein of interest, treatments were conducted 24 hours post-transfection. Otherwise, cells were grown to ~80% confluency prior to treatment. Cells were treated with either DMSO (vehicle) or the compound concentration indicated in the respective figure legend, and incubated at 37˚C for the indicated time. For isoTOP-ABPP analyses, all fragments were screened at 100 µM *in situ* for 1 hour, except for EN450, which was screened at 100 µM for 4 hours.

**Cell harvesting and lysis.** Cells were washed with cold PBS, scraped into cold PBS and harvested by centrifugation at 1800 *g* for 3 minutes. Cell pellets were then washed twice with cold PBS. Unless otherwise indicated, cells were lysed in 0.3% 3-[(3-Cholamidopropyl) dimethylammonio]-1-propanesulfonate (CHAPS) in PBS (pH 7.4) with 1X protriethylaminese inhibitor cocktail (Sigma, 11836170001) and incubated on ice for 30 minutes (protriethylaminese inhibitor cocktail was excluded from any lysates that required CuAAC 'click' conjugation). After lysis, cellular debris was clarified by centrifugation at 21,100 *g* for 5 minutes and the soluble fraction transferred to a fresh microcentrifuge tube. If applicable, the insoluble fractions were re-solubilized with 8M urea in PBS. Protein concentrations were determined using a BioRad DC protein assay kit from BioRad Life Science (Hercules, CA) and the lysate diluted to 2-5 mg/mL, with all samples of the same experiment normalized to the same concentration.

**Western Blots.** Lysate was normalized to 2-5 mg/mL, 25 µL of each lysate was added to 8 µL of 4X SDS loading dye (BioRad), and the samples were denatured at 95°C for 5 minutes. At least 60 µg of protein was resolved on a 4-20% SDS-PAGE gel (BioRad). Gels were transferred to nitrocellulose or activated PVDF membranes (BioRad) using a semi-dry transfer system (BioRad) and membranes blocked in 5% (w/v) milk in Tris-buffered saline (TBS) for 30 minutes by rocking at room temperature. Membranes were rocked with primary antibodies (**Table S6**) in 5% (w/v) milk in TBS overnight (14-16 hours) at 4°C, then washed 3 times with TBS for 5 minutes. Membranes were then rocked with secondary antibodies (**Table S6**) in 5% (w/v) milk in TBST (Tris-buffered saline with 0.1% Tween20) for 1 hour at room temperature, then washed 3 times with TBS. Membrane was imaged on a BioRad ChemiDoc Imaging System. Antibodies used were listed in Table S6. ImageJ[2] was used to normalize and quantify band intensities.

**Calculation of apparent DC50.** HEK293T cells were transfected with nsp14-FLAG for 36 hours, then treated with varying concentrations of JC19 (final concentrations 0 µM, 5 µM, 10 µM, 25 µM, 50 µM, and 100 µM) for 30 minutes at 37°C in biological triplicate. Cells were harvested, lysed, and lysates normalized to 2 mg/mL. 25 µL of each lysate was added to 8 µL of 4X SDS loading dye (BioRad) and the samples were denatured at 95°C for 5 minutes. Samples were resolved on 4-20% SDS-PAGE gels (BioRad) and transferred to a nitrocellulose membrane. The nsp14-FLAG signal was detected by an anti-FLAG primary antibody (**Table S6**). Membranes were imaged using a BioRad ChemiDoc Imaging System. Nsp14 bands of each sample were quantified using ImageJ[2] and plotted in GraphPad Prism 7 to obtain an apparent $DC_{50}$ value. Briefly, the intensity of the nsp14 band for each replicate of each JC19 treatment condition was quantified by ImageJ[2], and intensity values were normalized to the DMSO control in GraphPad Prism. Normalized values were then transformed to logarithms in GraphPad Prism and the IC50 calculated using a nonlinear regression curve fit.

**Immunoprecipitation for Western Blot.** HEK293T cells were transfected with the specified constructs for 36 hours and treated with the specified compounds indicated on the respective figures (**Figure 3B and Figure 3G**). Cells were harvested, lysed, and normalized as described in 'Cell harvesting and lysis.' To prepare anti-FLAG EZView resin (Sigma, F2426) for immunoprecipitation, 10 µL of anti-FLAG EZView resin per sample was washed in 600 µL TBS 2 times, then resuspended in 10 µL TBS/sample. Lysates were immunoprecipitated by rotating 500 µL of lysates with 10uL of anti-FLAG EZView resin per sample at 4°C for 2 hours. At least 50uL of the original lysates was kept for detection of input samples. Resin was spun down at 8200 *g* for 1.5 minutes and supernatant aspirated, followed by 3 washes with 500 µL of ice cold TBS (pH 7.4). Resin was then boiled in 30uL of 4X SDS loading dye (absent of Beta-mercaptoethanol) for 10 minutes to release immunoprecipitated proteins. 80 µg of each input lysate and 15uL of each immunoprecipitated sample was loaded and separated on 4-20% SDS-PAGE gels (BioRad). Gels were transferred to nitrocellulose membranes and taken through the standard western blotting procedure as described in the 'Western Blots' section.

**siRNA Knockdowns.** 10 µM siRNA stocks (**Table S7**) resuspended in nuclease-free duplex buffer (IDT, Cat. No. 11-01-03-01) were combined with 140 µL of DharmaFECT Transfection Reagent #1 (Horizon Discovery, Cat. No. T-2001-01) at a dilution of 1:100 with HBSS (Gibco) for a final concentration of 25 nM of siRNA per well for a 6-well cell culture plate. The siRNA and Transfection Reagent #1 mixture in HBSS was incubated at ambient temperature for 20 mins. HEK293T cells at a density of ~4.5 $x10^5$ cells/well were then reverse transfected with the incubated siRNA/Transfection Reagent #1 mix. Cells were left in the 37˚C incubator for 48 hours before either harvesting or transfecting. For transfections, cells were transfected with nsp14-FLAG plasmid (as described above) for 24 hours, treated with compounds (if stated), and then harvested as described in 'Cell harvesting and lysis.' All samples were normalized and prepared for analysis by SDS-PAGE and western blot analysis as described in 'Western blots.' For analysis by mass spectrometry, samples were prepared as stated below in the "Protein abundance sample preparation" section. All siRNAs used in this study are listed in Table S7.

**IsoTOP-ABPP sample preparation.** HEK293T cells were transfected with nsp14-FLAG for 36 hours (unless otherwise indicated) and treated with either DMSO or electrophilic fragment in at least biological duplicate. Cells were then harvested, lysed, and lysate concentration normalized to 2 mg/mL as described in 'Cell harvesting and lysis'. 200 µL of normalized lysates were then treated with 200 µM iodoacetamide alkyne (IAA) for 1 hour in the dark at ambient temperature. Samples were then subjected to bioorthogonal

copper(I)-catalyzed azide-alkyne cycloaddition (CuAAC) ‘click’ conjugation with either heavy (DMSO treated samples) or light (compound treated samples) biotin-azide tags containing a TEV cleavable linker[7] for 1 hour at ambient temperature.
Click reaction was performed by adding 24 µL of a premixed cocktail of click reagents (TEV tags (4µl of 5 mM stock per sample, final concentration= 100 µM), TCEP (4 µl of fresh 50 mM stock in water per sample, final concentration 1 mM), TBTA (12 µl of 1.7 mM stock in DMSO/t-butanol 1:4 per sample, final concentration = 100 µM), and CuSO4 (4 µl of 50 mM stock in water per sample, final concentration = 1 mM) to each 200 µL lysate sample. Protein was then solubilized by addition of SDS (final concentration = 1%), and treated with benzonase (0.5 µL) at 37˚C for 30 minutes. Equal volumes (200 µL) of heavy and light samples were then combined and subject to Single-Pot Solid-Phase-enhanced sample-preparation (SP3) clean-up. For preparation of SP3 beads, 40 µL Sera-Mag SpeedBeads Carboxyl Magnetic Beads per sample, hydrophobic (GE Healthcare, 65152105050250, 50 µg/µL, total 1 mg) and 40 µL Sera-Mag SpeedBeads Carboxyl Magnetic Beads per sample, hydrophilic (GE Healthcare, 45152105050250, 50 µg/µL, total 1 mg) were mixed and washed with water three times, then resuspended in 80 µL water per sample. 80 µL of the bead slurry was then transferred to the lysate samples (800 µg protein input) and incubated for 5 minutes at room temperature with shaking (1000 rpm). 600 µL of absolute ethanol was added to each sample and incubated for 10 minutes at room temperature with shaking (1000 rpm). The beads were then washed (3x, 500 µL 80% ethanol) using a magnetic rack (Sergi Lab Supplies, Cat. No. 1005a). Beads were then resuspended in 200 µL 2M urea in 0.5% SDS/PBS. DTT was added to a final concentration of 10 mM to each sample and incubated at 65°C for 15 minutes. Iodoacetamide was added to each sample to a final concentration of 20 mM and incubated at 37°C for 30 minutes. Next, absolute ethanol (500 µL) was added to each sample, and the samples were incubated for 5 min at room temperature with shaking (1000 rpm). The beads were then washed (3x, 500 µL 80% ethanol) using a magnetic rack. Next, beads were resuspended in 150 µL 2M urea in PBS and 3 µL of 1 mg/mL trypsin solution was added. Lysates were digested for 12-16 hours at 37ºC with shaking (200 rpm). After digestion, 3.8 mL acetonitrile (> 95% of the final volume) was added to each sample and the mixtures were incubated for 10 min at room temperature with shaking (1000 rpm). The beads were then washed (3x, 1 mL acetonitrile) with a magnetic rack. Peptides were eluted from SP3 beads with 100 µL of 2% DMSO in molecular biology-grade water for 30 min at 37°C with shaking (1000 rpm). The elution was repeated again with 100 µL of 2% DMSO in molecular biology-grade water and the elution fractions from the same sample combined. The samples were then enriched on Pierce streptavidin agarose beads in PBS (Thermo Scientific, Cat. No. 20353). To prepare streptavidin resin, 50 mL of resin per sample was washed in 10 mL PBS and collected by centrifugation at 1800 *g* for 3 minutes. PBS was then aspirated and the resin resuspended in 1 mL PBS/per sample. 1 mL of the resin slurry was then added to each sample and samples incubated for 2 hours with rotation at ambient temperature. Samples were then spun down at 1800 *g* for 3 minutes and supernatant aspirated. Streptavidin resin was then washed 2x with 1 mL PBS, then washed 2x with 1 mL water, collecting resin by centrifuging at 1800 *g* for 3 minutes in between each wash. Final wash was aspirated and the beads were resuspended in 75 µL of 1X TEV solution (50 mM Tris, pH 8, 0.5 mM EDTA, 1 mM DTT). 1.5 µl TEV protease (2 mg/mL; MacroLab, UC Berkeley) was added. Samples were then

incubated with rotation at 30˚C for 7 hours. Samples were then spun at 1800 *g* for 3 minutes and the supernatant was collected. The beads were then washed with 20 µL of water, spun, and the wash was collected and combined with the supernatant. Trifluoroacetic acid (TFA) was added to the samples to a final concentration of 0.5%. Samples were then cleaned up with Pierce C18 spin tips (Thermo Fisher, Cat. No. 87784) according to the manufacturer's protocol. The samples were dried (SpeedVac) and reconstituted with mass spectrometry buffer (5% acetonitrile and 1% formic acid in molecular biology-grade water) and analyzed by LC-MS/MS.

**AP-MS sample preparation.** Stable-Isotope Labeling by Amino Acids in Cell Culture (SILAC) HEK293T cells were used for affinity purification-mass spectrometry (AP-MS) sample preparation (**Figure 3D, 3E, 3F**). 6 10-cm plates of 'heavy' labeled HEK293T cells and 6 10-cm plates of 'light' labeled HEK293T cells were transfected with nsp14-FLAG for 36 hours. All 6 'light' labeled plates were then treated with DMSO for 1 hour, while 3 'heavy' labeled plates were treated with 100 µM JC19 for 1 hour and 3 'heavy' labeled plates were treated with 100 µM NB001 for 1 hour. All cells were harvested, lysed, and lysate concentrations normalized to 2.5 mg/mL. 120 µL of anti-FLAG EZView resin (Sigma, F2426) (10 µL/sample) was washed 2 times with 1X TBS and then resuspended in 120 µL of 1X TBS. 10 µL of washed EZView resin was added to each 350 µL lysate sample and rotated at 4°C for 2 hours. Resin was then spun down at 8200 *g* for 1.5 minutes and supernatant aspirated, followed by 3 washes with 500 µL of ice-cold TBS (pH 7.4). Bound proteins were eluted in 40 µL 8M urea in PBS at 37°C and 400 rpm for 30 minutes. Resin was spun down and eluant transferred to a new microcentrifuge tube. Resin was washed in an additional 20 µL of 8M urea in PBS and combined with previous eluant fraction. 50 µL of each 'heavy' lysate was then combined with a corresponding 50 µL of 'light' lysate. DTT was added to each sample to a final concentration of 10 mM and incubated at 65°C for 15 minutes. Iodoacetamide was added to each sample to a final concentration of 20 mM and incubated at 37°C for 30 minutes. Samples were diluted with PBS such that final urea concentration was 2M. 7 µL of 1 mg/mL trypsin was added to each sample and incubated at 37°C and 200 rpm for 12-16 hours to digest proteins. Trifluoroacetic acid (TFA) was added to the samples to a final concentration of 0.5%. Samples were then cleaned up with Pierce C18 spin tips (Thermo Fisher, Cat. No. 87784) according to the manufacturer's protocol. The samples were dried (SpeedVac) and reconstituted with mass spectrometry buffer (5% acetonitrile and 1% formic acid in molecular biology-grade water) and analyzed by LC-MS/MS.

**Proteasome activation assay.** HEK293T cells were seeded in a 6-well plate and grown to 90% confluence. Cells were pretreated as indicated in figure legends, then treated with 500 nM fluorescent proteasome probe Me4BodipyFL-Ahx3Leu3VS for 1 hour. Cells were then harvested and lysed as described in 'Cell harvesting and lysis.' Normalized cell lysates were then run on SDS-PAGE gel (BioRad) and in-gel fluorescence used to visualize proteasome activity.

**Click probe non-competition and competition enrichment sample preparation.** HEK293T cells were transfected with nsp14-FLAG for 36 hours. For non-competition enrichment experiments, cells or lysates were either treated with 100 µM electrophilic compound (*n* = 6 for in-cell labeling, *n* =3 for lysate labeling) or 100 µM alkynylated electrophilic probe (*n* = 6 for in-cell labeling, *n* =3 for lysate labeling) for 20 minutes. For competition-based enrichment experiments, cells were either treated with DMSO (*n* = 6 for in-cell labeling, *n* = 3 for lysate labeling) or 100 µM electrophilic compound (*n* = 6 for in-cell labeling, *n* = 2 for lysate labeling) for 20 minutes, then all conditions treated with 100 µM alkynylated electrophilic compound for 20 minutes. Cells were then harvested, lysed, and lysate concentration normalized to 3 mg/mL. 200 µL each lysate was then subject to CuAAC 'click' biotinylation by treatment with 24 uL of premixed cocktail of click reagents (12uL 1.7mM TBTA per sample, 4uL 50mM copper sulfate per sample, 4uL 50mM biotin-azide per sample, 4uL 50mM TCEP per sample). Samples were then cleaned via SP3 clean-up. 20 µL Sera-Mag SpeedBeads Carboxyl Magnetic Beads per sample, hydrophobic (GE Healthcare, 65152105050250, 50 µg/µL, total 1 mg) and 20 µL Sera-Mag SpeedBeads Carboxyl Magnetic Beads per sample, hydrophilic (GE Healthcare, 45152105050250, 50 µg/µL, total 1 mg) were mixed and washed with water three times and resuspended in 40 µL of water per sample. 40 µL of the bead slurry was then transferred to the lysate samples (500 µg protein input) and incubated for 5 minutes at room temperature with shaking (1000 rpm). 500uL of absolute ethanol was added to each sample and incubated for 10 minutes at room temperature with shaking (1000 rpm). The beads were then washed (3x, 400uL 80% ethanol) with a magnetic rack. Proteins were eluted from SP3 beads with 100 µL of 0.2% SDS in PBS for 30 min at 37°C with shaking (1000 rpm). The elution was repeated again with 100 µL of 0.2% SDS in PBS and the elution fractions from the same sample combined. Streptavidin resin (Thermo Scientific, Cat. No. 20353) was then prepared by washing 50 µL of streptavidin resin per sample in 10mL PBS and collecting resin by centrifugation at 1800 *g* for 5 minutes, PBS aspirated, and resin resuspended in 1 mL PBS per sample. 1mL of streptavidin slurry was added to each sample and samples rotated at room temperature for 1.5 hours. Streptavidin resin was collected via centrifugation at 1500 *g* for 2 minutes and washed 1 time with 1 mL 0.2% SDS in PBS, 3 times with 1 mL PBS, and 3 times with 1 mL molecular biology-grade water. Resin was then resuspended in 200 µL 6M urea. DTT was then added to each sample to a final concentration of 10 mM and incubated at 65°C for 15 minutes. Iodoacetamide was then added to each sample to a final concentration of 20 mM and incubated at 37°C for 30 minutes. Urea was diluted to ~2M, resin collected via centrifugation at 1500 *g* for 2 minutes, supernatant aspirated, and resin resuspended in 150 µL 2M urea in PBS. Then, 3 µL of 1 mg/mL trypsin was added to each sample and samples were subject to digestion for 12-16 hours at 37°C and 200 rpm. Resin was then spun down and supernatant transferred to a new tube (containing digested peptides). Resin was washed with an additional 50 µL of molecular biology-grade water, spun down, and supernatant added to the previous peptide fraction. Trifluoroacetic acid (TFA) was added to the samples to a final concentration of 0.5%. Samples were then cleaned up with Pierce C18 spin tips (Thermo Fisher, Cat. No. 87784) according to the manufacturer's protocol. The samples were dried (SpeedVac) and reconstituted with mass spectrometry buffer (5% acetonitrile and 1% formic acid in molecular biology-grade water) and analyzed by LC-MS/MS.

**Bulk proteomics sample preparation.** HEK293T cells were transfected with nsp14-FLAG for 36 hours and treated with either DMSO or electrophilic compound in biological triplicate. Cells were then harvested, lysed, and lysate concentration normalized to 3 mg/mL. Samples were then cleaned using SP3 clean-up. 20 µL Sera-Mag SpeedBeads Carboxyl Magnetic Beads per sample, hydrophobic (GE Healthcare, 65152105050250, 50 µg/µL, total 1 mg) and 20 µL Sera-Mag SpeedBeads Carboxyl Magnetic Beads per sample, hydrophilic (GE Healthcare, 45152105050250, 50 µg/µL, total 1 mg) were mixed and washed with water three times and resuspended in 40 µL water per sample. 40 µL of the bead slurry was then transferred to the lysate samples (500ug protein input) and incubated for 5 minutes at room temperature with shaking (1000 rpm). 500 µL of absolute ethanol was added to each sample and incubated for 10 minutes at room temperature with shaking (1000 rpm). The beads were then washed (3x, 400uL 80% ethanol) with a magnetic rack. Beads were then resuspended in 200 µL 2M urea in 0.5% SDS/PBS. DTT was added to each sample to a final concentration of 10 mM to each sample and incubated at 65°C for 15 minutes. Iodoacetamide was added to each sample to a final concentration of 20 mM and incubated at 37°C for 30 minutes. Absolute ethanol (400 µL) was then added to each sample, and the samples were incubated for 5 minutes at room temperature with shaking (1000 rpm). Beads were then washed (3x, 400uL 80% ethanol) with a magnetic rack. Next, beads were resuspended in 200 µL 2M urea in PBS and 3 µL of 1 mg/mL trypsin solution was added. Proteins were digested for 12-16 hours at 37ºC with shaking (200 rpm). After digestion, 3.8 mL acetonitrile (> 95% of the final volume) was added to each sample and the mixtures were incubated for 10 minutes at room temperature with shaking (1000 rpm). The beads were then washed (3x, 1mL acetonitrile) with a magnetic rack. Peptides were eluted from SP3 beads with 100 µL of 2% DMSO in molecular biology-grade water for 30 min at 37°C with shaking (1000 rpm). The elution was repeated again with 100 µL of 2% DMSO in molecular biology-grade water and the elution fractions from the same sample combined. The samples were dried (SpeedVac), then reconstituted with 5% acetonitrile and 1% formic acid in molecular biology-grade water and analyzed by LC-MS/MS.

**Pathway analysis.** For KEGG and Gene Ontology analyses, the subset of proteins to be analyzed was searched using the Enrichr search algorithm[8,9,10], and plotted using the GSEApy package[10]. The entirety of the proteins in the dataset used as the 'background' dataset for statistical comparison. Pathway groups with an adjusted p-value <0.05 were considered significantly enriched over background.

**Cellular Compartment analysis.** For the cell compartment analysis done on the isoTOP-ABPP samples, the isoTOP data was first filtered based on proteins containing multiple identified cysteines across the aggregate dataset. The bulk multi-cysteine containing protein dataset was crossed with Uniprot keyword annotations[11] and the proteins were grouped based on each keyword annotations for cellular compartment. Then, for each compound, proteins were grouped as having more than 2 cysteines with high ratios or as having less than 2 cysteines with high ratios. The frequency at which proteins from each cellular compartment were identified from the whole dataset versus the dataset with more than 2 high ratio cysteines was calculated and, finally, a fold change ratio was calculated to assess enrichment of proteins from a particular subcellular compartment.

**Immunocytochemistry.** Sterilized coverslips were placed in each well of a 24 well plate and coated with poly-d-lysine for 30 minutes at 37°C. Cells were seeded on coverslips overnight, then treated with the indicated compounds. For nsp14 and nsp16 experiments, viral proteins were transfected into cells for at least 24 hours prior to compound treatment. For doxycycline-inducible U2OS cells, cells were cultured with media containing 1 µg/mL doxycycline for at least 24 hours prior to fixation for expression of G3BP1-TurboID-V5. Media was aspirated and each well washed 1 time with 500 µL DPBS, followed by fixation with 3.7% formaldehyde in DPBS for 15 minutes at room temperature. Each well was then washed 2 times with 500 µL DPBS and permeabilized with 500 µL 0.1% TritonX in DPBS for 6 minutes at room temperature. Cells were then washed 3 times with 500 µL of DPBS and blocked in 1% FBS in DPBS for 30 minutes at room temperature. Cells were then incubated with 300 µL primary antibodies (Table S6) in 0.1% FBS in PBS overnight at 4°C. Cells were then washed 2 times with 500 µL 0.1% Tween20 in PBS and 1 time with 500 µL PBS. Cells were then incubated with 300 µL secondary antibodies (**Table S6**) for 1 hour at room temperature. Cells were then incubated with 300 µL 1:400 phalloidin-488 stain (Invitrogen, A12379) for 30 minutes at room temperature, followed by staining with 500 µL 1 µg/mL DAPI stain for 5 minutes. Cells were then washed 2 times with 500 µL PBS and mounted onto glass slides using Aqua-Poly/Mount mounting media (Polysciences, Inc., 18606-20). Samples were left to sit in the dark for 24 hours. Slides were then imaged on a Zeiss LSM880 confocal microscope at 63X oil objective with 2X manual zoom. Quantification of fluorescence intensity was done on ImageJ[2] software using particle picker. Unpaired student's t-test was performed to generate p-value between control and treatment groups. Colocalization analyses were done on ImageJ using the Coloc2 plugin.

**Glutathione assay.** Cells were grown to 80% confluence and treated with either DMSO (vehicle), 100 µM JC19 for 1 hour, or 500 µM Buthionine sulfoximine (BSO) for 24 hours in duplicate. Total glutathione content in cells was measured using Calorimetric Glutathione Assay (G Biosciences, 786-075) according to the manufacturer's Cell Lysate Preparation protocol using the end-point method. Only deviation from protocol is that cell lysates were prepared at 4X the concentration suggested in the protocol.

**IncuCyte Live Cell Imaging.** HeLa cells were seeded in a 96-well plate at a density of 100,000 cells/well and allowed to adhere for 6 hours in a tissue culture incubator at 37° C with 5% $CO_2$. 30 mins before first live cell time point, addition of media containing Cytotox Green reagent (Cat. No. 4633) (final dilution 1:4000) and compound was performed. An IncuCyte SX5 Live-Cell Imaging and Analysis System (Sartorius) was used for data acquisition. Over the course of 13 hours, at each 1 hour time point, four images per well were taken using both the 10X phase contrast and green lenses with the adherent cell-by-cell option selected. Analysis was performed using the IncuCyte 2019B Rev2 software. In brief, the green integrated intensity per well was normalized to 0d0h0m per condition and each condition was plated in triplicate.

**XRNAX.** Stable-Isotope Labeling by Amino Acids in Cell Culture (SILAC) HEK293T cells were used for XRNAX experiments. 'Light' labeled cells were treated with DMSO for 1 hour in duplicate and 'heavy' labeled cells were treated with 100 µM JC19 for 1 hour in duplicate. Cells were then exposed to 254 nm light at 200 mJ/cm$^2$ on ice to initiate crosslinking of RNA to RNA-binding proteins. Cells were then harvested and cells counted on a cell counter. Equal number of heavy and light labeled cells (~6.0x10$^6$ cells) were then combined and collected by centrifugation at 1800 *g* for 5 minutes. Cells were lysed in 1 mL TRIzol (Invitrogen, 15596026) until homogenous and let to sit for 10 minutes at room temp. 200 µL of chloroform was added to each sample and samples shook vigorously to mix, then let to incubate at room temperature for 5 minutes. Samples were centrifuged at 7000 *g* for 10 minutes at 4°C. Samples separate into three phases; the top layer was discarded and the middle interphase layer (light pink/white jelly-like) transferred to a new tube. Wash interphase layer 2x with 1 mL TE buffer + 0.1% SDS. Disintegrate pellet with 1 mL TE buffer + 0.1% SDS and spin down at 5000 *g* for 5 minutes and transfer supernatant to new tube. Repeated this disintegration step 3 additional times to get 4 fractions for each sample. To each fraction, added 60 µL 5M NaCl, 1 µL glycoblue (Fisher, AM9515), and 1 mL isopropanol, mix, and incubate on ice for 15 minutes to encourage precipitation. Spun each tube at 18,000 *g* and -9°C for 15 minutes and removed supernatant from all fractions. Used 1 mL 80% ethanol to combine pellets from the 4 fractions from each sample and spin down pellets at 18,000 *g* for 1 minute. Removed ethanol supernatant and added 900 µL RNAse-free water to reconstitute pellets. Incubated each sample at 65°C for 5 minutes and let cool to room temperature. Added 100 µL 10X DNAse buffer (100 mM Tris-HCl, 1 mM $CaCl_2$, 25 mM $MgCl_2$, pH 7.5) to each sample followed by addition of 1 µL DNAse to each sample and incubated at 37°C for 1.5 hours with shaking (600 rpm). Transferred 1 mL samples to 2 mL microcentrifuge tubes and added 60 µL 5M NaCl and 1 mL isopropanol to each sample, then incubated samples on ice for 30 minutes. Spun down samples at 18,000 *g* for 15 minutes at -5°C, then removed supernatant and resuspended pellet in 1 mL 80% ethanol and transferred back to microcentrifuge tubes. Spun down samples at 21.1k *g* for 5 minutes to pellet RNA and removed ethanol supernatant carefully so as not to disturb pellets and let pellets dry for ~5-10 minutes. Resuspended pellets in 100 µL PBS and let sit at room temp for about an hour to allow full resuspension. Samples were then cleaned using SP3 clean-up. 20 µL Sera-Mag SpeedBeads Carboxyl Magnetic Beads per sample, hydrophobic (GE Healthcare, 65152105050250, 50 µg/µL, total 1 mg) and 20 µL Sera-Mag SpeedBeads Carboxyl Magnetic Beads per sample, hydrophilic (GE Healthcare, 45152105050250, 50 µg/µL, total 1 mg) were mixed and washed with water three times and resuspended in 40 µL water per sample. 40 µL of the bead slurry was then transferred to the lysate samples (500ug protein input) and incubated for 5 minutes at room temperature with shaking (1000 rpm). 500 µL of absolute ethanol was added to each sample and incubated for 10 minutes at room temperature with shaking (1000 rpm). The beads were then washed (3x, 400uL 80% ethanol) with a magnetic rack. Beads were then resuspended in 200 µL 2M urea in 0.5% SDS/PBS. DTT was added to each sample to a final concentration of 10 mM to each sample and incubated at 65°C for 15 minutes. Iodoacetamide was added to each sample to a final concentration of 20 mM and incubated at 37°C for 30 minutes. Absolute ethanol (400 µL) was then added to each sample, and the samples were incubated for 5 minutes at room temperature with shaking (1000 rpm). Beads were then washed (3x, 400uL 80% ethanol) with a magnetic rack. Next, beads were resuspended in 200 µL 2M urea in PBS and 3 µL of 1 mg/mL trypsin solution was added. Proteins were digested for 12-16 hours at 37ºC with shaking (200 rpm). After digestion, 3.8 mL acetonitrile (> 95%

of the final volume) was added to each sample and the mixtures were incubated for 10 minutes at room temperature with shaking (1000 rpm). The beads were then washed (3x, 1mL acetonitrile) with a magnetic rack. Peptides were eluted from SP3 beads with 100 μL of 2% DMSO in molecular biology-grade water for 30 min at 37°C with shaking (1000 rpm). The elution was repeated again with 100 μL of 2% DMSO in molecular biology-grade water and the elution fractions from the same sample combined. The samples were dried (SpeedVac), then reconstituted with 5% acetonitrile and 1% formic acid in molecular biology-grade water and analyzed by LC-MS/MS.

**Liquid-chromatography tandem mass-spectrometry (LC-MS/MS) acquisition.** The samples were analyzed by liquid chromatography tandem mass spectrometry using a Thermo Scientific™ Orbitrap Eclipse™ Tribrid™ mass spectrometer. Peptides were fractionated online using a 18 cm long, 100 μM inner diameter (ID) fused silica capillary packed in-house with bulk C18 reversed phase resin (particle size, 1.9 μm; pore size, 100 Å; Dr. Maisch GmbH). The 70-minute water/acetonitrile gradient was delivered using a Thermo Scientific™ EASY-nLC™ 1200 system at different flow rates (Buffer A: water with 3% DMSO and 0.1% formic acid and Buffer B: 80% acetonitrile with 3% DMSO and 0.1% formic acid). The detailed gradient for each experiment is outlined in **Table S10**. Data was collected with charge exclusion (1, 8,>8). Data was acquired using a Data-Dependent Acquisition (DDA) method consisting of a full MS1 scan (Resolution = 120,000) followed by sequential MS2 scans (Resolution = 15,000) to utilize the remainder of the 1 second cycle time. Precursor isolation window was set as 1.6 and normalized collision energy was set as 30%. Some datasets were collected with a high-field asymmetric waveform ion mobility (FAIMS) device with current voltages of -35, -45, and -55. Files of MS data with corresponding experiments can be found in **Table S11**.

**Proteomic data availability.** The MS data have been deposited to the ProteomeXchange Consortium via the PRIDE partner repository with the dataset identifier PXD046278 and PXD046393. File details can be found in **Table S11**.

**Data Compilation and Statistics.** Raw data collected by LC-MS/MS were searched with MSFragger and FragPipe (version 17-19)[12–14]. The proteomic workflow and its collection of tools was set as default. Precursor and fragment mass tolerance was set as 20 ppm. Missed cleavages were allowed up to 1. Peptide length was set 7 - 50 and peptide mass range was set 500 - 5000. For SILAC based experiments, MS1 labeling quant was enabled with Light set as K;R+0 and Heavy set as K+8.0142 and R+10.0083. MS1 intensity ratio of heavy and light labeled cysteine peptides and compiled for individual proteins were reported. For isoTOP-ABPP experiments, MS1 labeling quant was enabled with Light set as C+521.3074 and Heavy set as C+527.3213. MS1 intensity ratio of heavy and light labeled cysteine peptides were reported. For Glygly searches, a variable modification of +114.0429 was included on lysine. Calibrated and deisotoped spectrum files produced by FragPipe were retained and reused for this analysis. For protein-level analyses quantified using label-free quantification, identified proteins were filtered by Perseus[15,16] to retain only proteins identified in at least 2 replicates (for experiments with 3 replicates/condition) or 3 replicates (for experiments with 6 replicates/condition) of at least 1 experimental condition per experiment. Missing values were then imputed by Perseus based on the normal distribution. For experiments utilizing MS1 quantitation (e.g.

isoTOP-ABPP and SILAC), custom python scripts were implemented to compile labeled peptide and protein datasets. Unique proteins, unique cysteines, and unique peptides were quantified for each dataset. Unique proteins were established based on UniProt protein IDs. Unique peptides were found based on sequences containing a modified cysteine residue. Unique cysteines were classified by an identifier consisting of a UniProt protein ID and the amino acid number of the modified cysteine (ProteinID_C#); residue numbers were found by aligning the peptide sequence to the corresponding UniProt protein sequence. When there are multiple cysteines in one peptide, all the cysteine residue numbers will be reported as ProteinID_C#_C# though it is undistinguishable on which cysteine the modification is. Statistical values including the exact n, statistical test, and significance are reported in the Figure Legends. Statistical significance was defined as p-value < 0.05 and unless indicated otherwise determined by an unpaired 2-tailed Student's t-test.

**Statistics.** Statistical analysis was performed using GraphPad Prism (v9.4.1) for Mac (GraphPad Software). Statistical values including the exact n, statistical test, and significance are reported in the Figure Legends. Statistical significance was defined as p-value < 0.05 and unless indicated otherwise determined by an unpaired 2-tailed Student's t-test.

#### (D) Chemistry Methods

**General Methods**. All reactions were performed in dried glassware under an atmosphere of dry $N_2$ unless otherwise stated. Silica gel P60 (SiliCycle) was used for column chromatography. Plates were visualized by fluorescence quenching under UV light or by staining with iodine, $KMnO_4$, or bromocresol green. Other reagents were purchased from Sigma-Aldrich (St. Louis, MO), Alfa Aesar (Ward Hill, MA), EMD Millipore (Billerica, MA), Fisher Scientific (Hampton, NH), Oakwood Chemical (West Columbia, SC), Combi-blocks (San Diego, CA) and Cayman Chemical (Ann Arbor, MI) and used without further purification. $^1H$ NMR and $^{13}C$ NMR spectra for characterization of new compounds and monitoring reactions were collected in $CDCl_3$, $CD_3OD$, $D_2O$ or DMSO-$d_6$ (Cambridge Isotope Laboratories, Cambridge, MA) on a Bruker AV 500 MHz spectrometer or Bruker AV 400 MHz in the Department of Chemistry & Biochemistry at The University of California, Los Angeles. All chemical shifts are reported in the standard notation of parts per million using the peak of residual proton signals of the deuterated solvent as an internal reference. Coupling constant units are in Hertz (Hz). Splitting patterns are indicated as follows: br, broad; s, singlet; d, doublet; t, triplet; q, quartet; m, multiplet; dd, doublet of doublets; dt, doublet of triplets. Low-resolution mass spectrometry was performed on an Agilent Technologies InfinitiyLab LC/MSD single quadrupole LC/MS (ESI source). High-resolution mass spectrometry was performed on a Waters LCT Premier with ACQUITY LC and autosampler (ESI source). All compounds named with prefix **KB** were purchased from MilliporeSigma or prepared according to our previous procedures[17].

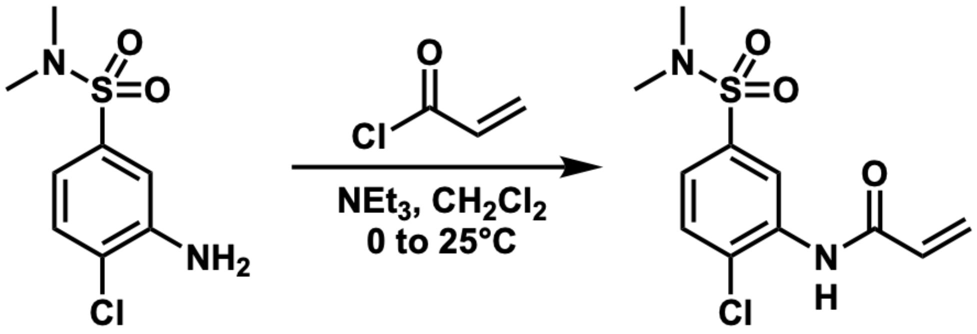


**N-(2-chloro-5-(N,N-dimethylsulfamoyl)phenyl)acrylamide (EN450)**

To a solution of 3-amino-4-chloro-N,N-dimethylbenzenesulfonamide (100 mg, 1 Eq, 426 µmol) in DCM (15 mL) was added triethylamine (86.2 mg, 2 Eq, 852 µmol) dropwise. The reaction was then brought to 0°C and stirred for 5 minutes. Acryloyl chloride (57.8 mg, 1.5 Eq, 639 µmol) was then added and the reaction was brought to 230 and left to stir for 16 hours. Mixture was quenched with H2O (5mL), extracted with DCM (3 x 5mL), organic layer was washed with brine (3mL), and dried with NaSO4 and concentrated in vacuo. Column chromatography (eluting with 30% EtOAC in hexanes) afforded a white solid (62.0 mg, 50%). 1H NMR (400 MHz, CDCl3): δ (d, J = 2.1 Hz, 1H), 7.56 (d, J = 8.3 Hz, 1H), 7.52 (d, J = 2.1 Hz, 1H), 6.50 (dd, J = 16.9, 1.0 Hz, 1H), 6.32 (dd, J = 16.9, 10.2 Hz, 1H), 5.89 (dd, J = 10.2, 1.0 Hz, 1H), 2.78 (s, 6H). All analyses were consistent with previously reported data [18].

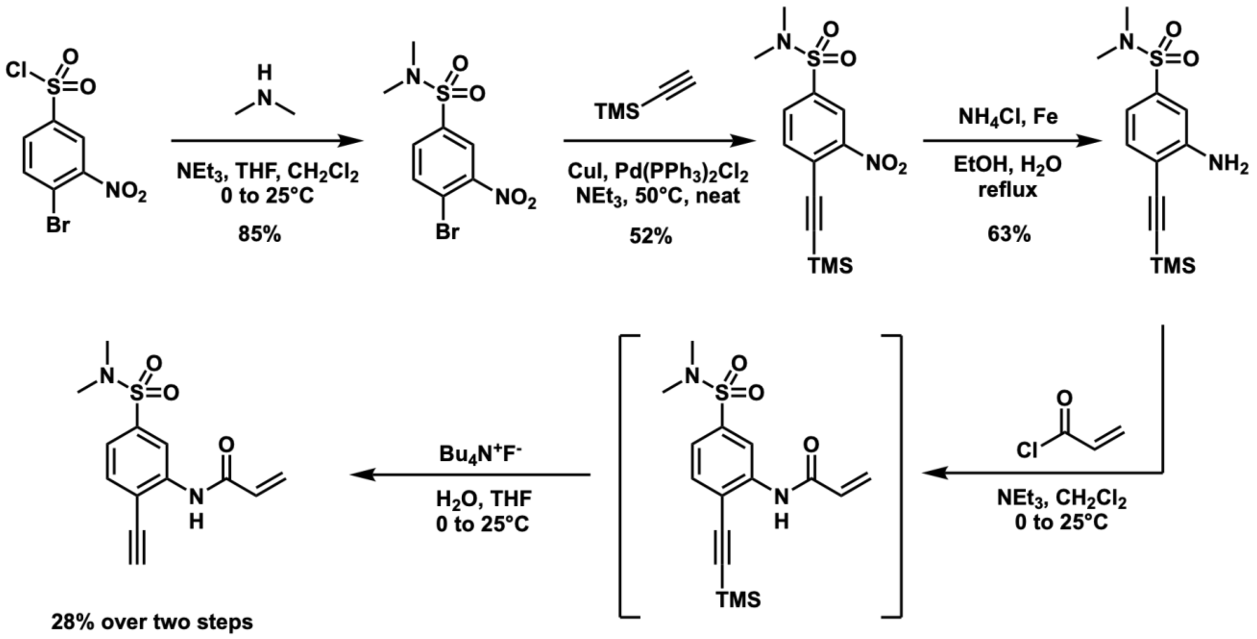


**Synthesis of EN450yne**

**4-bromo-N,N-dimethyl-3-nitrobenzenesulfonamide (S1)**

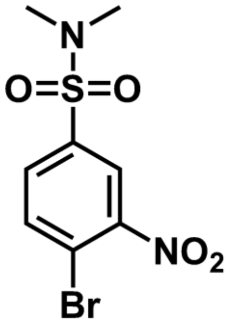

To an oven dried 25mL round-bottom flask was added dimethylamine (27.2 mg, 302 µL, 2 molar, 1.21 Eq, 604 µmol) in $CH_2Cl_2$ (0.98 mL) and cooled to 0°C. Then with strong stirring, triethylamine (61.1 mg, 1.21 Eq, 604 µmol) was added drop wise and stirred at 0°C for 50 min. 4-bromo-3-nitrobenzenesulfonyl chloride (150 mg, 1 Eq, 499 µmol) in THF (0.43 mL) and $CH_2Cl_2$ (0.85 mL) were then added, reaction was brought to room temperature and stirred for 5 minutes. Mixture was then diluted with $CH_2Cl_2$ (3mL), quenched with $H_2O$ (1mL), extracted with $CH_2Cl_2$ (3 x 1mL), dried with $NaSO_4$, and concentrated in vacuo. Column chromatography (eluting with 30% EtOAC in hexanes) afforded a light-yellow solid (262 mg, 85%).

$^1$H NMR (400 MHz, $CDCl_3$) δ 8.20 (d, J = 2.1 Hz, 1H), 7.95 (d, J = 8.3 Hz, 1H), 7.80 (dd, J = 8.4, 2.1 Hz, 1H), 2.79 (s, 6H).

$^{13}$C NMR (101 MHz, $CDCl_3$) δ 137.36, 136.31, 131.45, 124.63, 119.61, 37.94.

HRMS (ESI-MS) *m/z*: Calculated $[M+H]^+$ = 308.9536 , Found $[M+H]^+$ = 308.9539

**N,N-dimethyl-3-nitro-4-((trimethylsilyl)ethynyl)benzenesulfonamide (S2)**

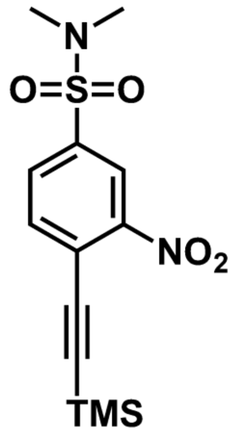


To an oven dried round-bottom flask was added $Pd(PPh_3)_2Cl_2$ (3.4 mg, 0.05 equiv, 4.9 µmol), CuI (0.92 mg, 0.05 equiv, 4.9 µmol), and 4-bromo-N,N-dimethyl-3-nitrobenzenesulfonamide (30 mg, 1 equiv, 97 µmol). The reaction flask was purged with argon followed by addition of dry triethylamine (0.15 mL). After a few seconds of stirring the reaction mixture turned yellow. Trimethylsilylacetylene (19 mg, 27 µL, 2 equiv, 0.19 mmol) was then added dropwise turning the mixture black. Upon complete addition the reaction mixture was heated to 50 °C and let stir. After 2 hours the reaction mixture was cooled to ambient temperature and filtered over cotton and celite. The resulting flask and solids were rinsed with hexanes and the combined filtrate was concentrated down. The crude mixture was purified using flash column chromatography (0 to 70% ethyl acetate in hexanes) to yield the desired product as a beige solid (16mg, 52%).

$^1$H NMR (400 MHz, $CDCl_3$) δ 8.37 (d, J = 1.8 Hz, 1H), 7.92 (dd, J = 8.2, 1.8 Hz, 1H), 7.81 (dd, J = 8.1, 0.4 Hz, 1H), 2.77 (s, 6H), 0.30 (s, 9H).
$^{13}$C NMR (75 MHz, $CDCl_3$) δ 150.16, 136.68, 135.94, 130.91, 123.64, 122.37, 108.88, 97.95, 37.84.
HRMS (ESI-MS) *m/z*: Calculated $[M+H]^+$ = 327.0829 , Found $[M+H]^+$ = 327.0838

**3-amino-N,N-dimethyl-4-((trimethylsilyl)ethynyl)benzenesulfonamide (S3)**

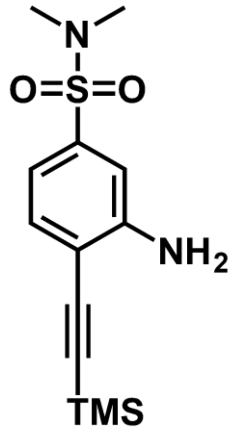


To an oven dried round bottom flask was added ammonium chloride (31 mg, 6 equiv, 0.59 mmol), Iron Powder (33 mg, 6 equiv, 0.59 mmol), EtOH (0.75 mL), Water (0.75 mL), and heated to 60°C for 30 minutes. After 30 minutes, N,N-dimethyl-3-nitro-4-((trimethylsilyl)ethynyl)benzenesulfonamide (32 mg, 1 equiv, 98 µmol) dissolved in EtOH (0.75 mL) and Water (0.75 mL) was added and mixture was refluxed at 80 °C. After 24 hours the mixture was quenched with $H_2O$ (5mL), extracted with $CH_2Cl_2$ (3 x 5mL). The combined organic layers were then washed with brine (1x3mL), dried with $NaSO_4$ and concentrated in vacuo. Crude material was purified using flash column chromatography (20 to 50% ethyl acetate in hexanes) to afford the desired product as a white solid (18.0 mg, 62%).

$^1$H NMR (400 MHz, $CDCl_3$) δ 7.41 (d, J = 8.1 Hz, 1H), 7.07 (s, 1H), 7.01 (dd, J = 8.0, 1.5 Hz, 1H), 2.69 (s, 6H), 0.28 (s, 9H).

$^{13}C$ NMR (101 MHz, $CDCl_3$) δ 136.09, 132.79, 116.35, 112.72, 38.00.
HRMS (ESI-MS) *m/z*: Calculated $[M+H]^+$ = 297.1087 , Found $[M+H]^+$ = 297.1146

**N-(5-(N,N-dimethylsulfamoyl)-2-ethynylphenyl)acrylamide (EN450yne)**

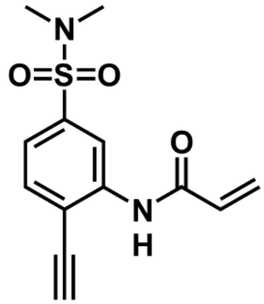

To an oven dried round-bottom flask containing 3-amino-N,N-dimethyl-4- ((trimethylsilyl)ethynyl)benzenesulfonamide (19mg, 1 equiv, 63μmol) was added dry $CH_2Cl_2$ (1mL). The flask was then capped and purged with argon followed by cooling to 0°C and addition of triethylamine (35μL, 4 equiv, 0.252 mmol). Next, acryloyl chloride (15μL, 3 equiv, 0.189mmol) was added and the reaction mixture allowed to warm to ambient temperature and stir for 16h. Upon completion the reaction mixture was quenched with water (1mL) and extracted with $CH_2Cl_2$ (3x2mL). The combined organic layers were dried over sodium sulfate, concentrated down, and filtered through a silica plug using 30% ethyl acetate in hexanes. The crude product was used directly for the next step.

To the crude N-(5-(N,N-dimethylsulfamoyl)-2-((trimethylsilyl)ethynyl)phenyl) acrylamide in a dry round-bottom flask was added THF (0.7 mL) and water (2.3 mg, .0023 mL, 2.0 equiv, 0.13 mmol). The flask was capped, purged with argon, and cooled to 0 °C followed by dropwise addition of TBAF (16 mg, .063 mL, 1 molar, 1.0 equiv, 63 µmol). Reaction mixture was then allowed to stir at 0°C for 10 minutes followed by warming to ambient temperature for 5 minutes. After 5 minutes, the mixture was quenched with $H_2O$ (2mL), extracted with EtOAc (3 x 5mL), organic layer, and dried with $NaSO_4$ and concentrated in vacuo. The crude material was purified by flash column chromatography (0 to 30% ethyl acetate in hexanes) to afford the desired compound as an off-white solid. (5mg, 28% over two steps)

$^1H$ NMR (400 MHz, CDCl3) δ 8.92 (d, J = 1.7 Hz, 1H), 8.09 (s, 1H), 7.62 (d, J = 8.0 Hz, 1H), 7.49 (dd, J = 8.1, 1.8 Hz, 1H), 6.48 (dd, J = 16.9, 1.1 Hz, 1H), 6.30 (dd, J = 16.9, 10.3 Hz, 1H), 5.87 (dd, J = 10.2, 1.1 Hz, 1H), 3.70 (s, 1H), 2.78 (s, 6H).
$^{13}C$ NMR (101 MHz, CDCl3) δ 163.50, 139.92, 137.30, 132.65, 130.73, 129.06, 122.53, 118.43, 114.89, 87.46, 77.94, 38.04.
HRMS (ESI-MS) *m/z*: Calculated $[M+H]^+$ = 279.0798 , Found $[M+H]^+$ = 279.0830

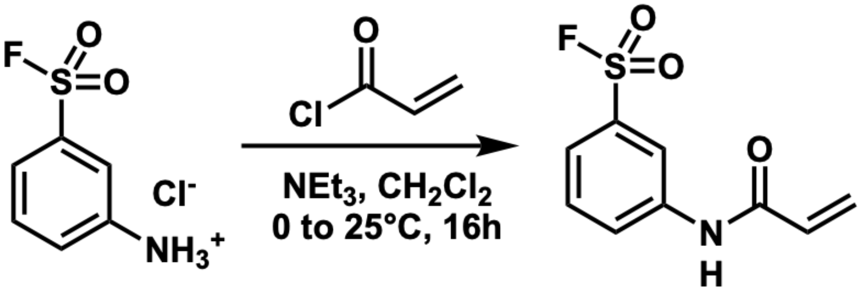


**3-acrylamidobenzenesulfonyl fluoride (JC17)**

Following the previously described procedure[19], to a solution of 3-aminobenzenesulfonyl fluoride hydrochloride (100 mg, 0.47 mmol, 1 equiv.) in 2 mL DCM was added triethylamine (198 µL, 1.42 mmol, 3 equiv.), acryloyl chloride (77 µL, 0.94mmol, 2 equiv.) in an ice bath. Next the reaction was allowed to react for 3h at RT. The reaction was quenched with 30 mL saturated NaHCO3 (aq), extracted with DCM (2 × 20 mL), and was concentrated and purified by silica column chromatography (2:1 hexanes/ EtOAc) to afford product (88.9 mg, 82%) as a colorless solid.All analyses were consistent with previously reported data.

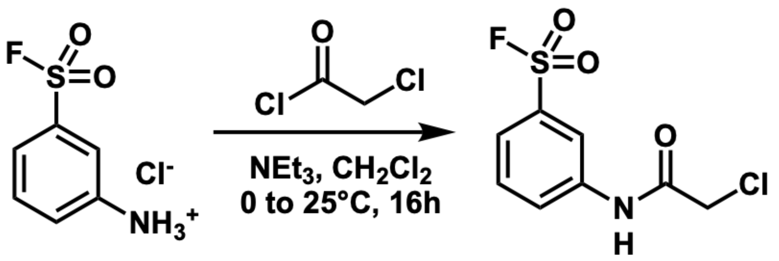


**3-(2-chloroacetamido)benzenesulfonyl fluoride (JC19)**

Following the previously described procedure[19], to a solution of 3-aminobenzenesulfonyl fluoride hydrochloride (100 mg, 0.47 mmol, 1 equiv.) in 2 mL DCM was added Triethylamine (198 µL, 1.42 mmol, 3 equiv.), 2- chloroacetyl chloride (75 µL, 0.94mmol, 2 equiv.) in an ice bath. Next the reaction was allowed to react for 3h at RT. The reaction was quenched with 30 mL saturated NaHCO3 (aq), extracted with DCM (2 × 20 mL), and was concentrated and purified by silica column chromatography (2:1 hexanes/ EtOAc) to afford product (108.1 mg, 91%) as a colorless solid.All analyses were consistent with previously reported data.

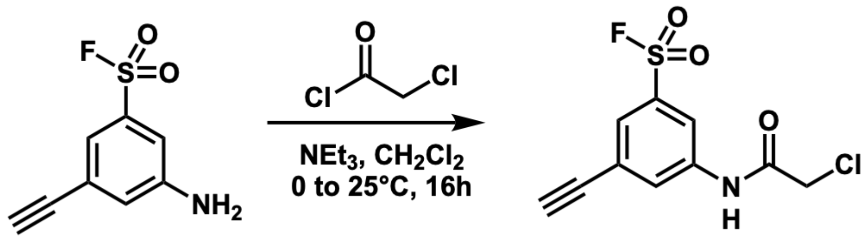


**3-(2-chloroacetamido)-5-ethynylbenzenesulfonyl fluoride (JC19yne)**

To a solution of 3-amino-5-ethynylbenzenesulfonyl fluoride hydrochloride (50 mg, 0.25 mmol, 1 equiv.) in 2 mL DCM was added triethylamine (104 µL, 0.75 mmol, 3 equiv.), 2-chloroacetyl chloride (40 µL, 0.50mmol, 2 equiv.) in an ice bath. Next the reaction was allowed to react for 3h at RT. The reaction was quenched with 30 mL saturated NaHCO3 (aq), extracted with DCM (2 × 20 mL), and was concentrated and purified by silica column chromatography (2:1 hexanes/ EtOAc) to afford product (63mg, 92%) as an off-white solid.

$^{1}$H NMR (400 MHz, Chloroform-*d*) δ 8.54 (s, 1H), 8.23 (t, *J* = 2.0 Hz, 1H), 8.10 (t, *J* = 1.8 Hz, 1H), 7.87 (t, *J* = 1.6 Hz, 1H), 4.24 (s, 2H), 3.27 (s, 1H).

$^{13}$C NMR (101 MHz, $CDCl_3$) δ 164.60, 138.39, 134.54, 134.29, 129.47, 127.89, 125.46, 119.31, 81.37, 80.39, 42.83.

$^{19}$F NMR (376 MHz, $CDCl_3$) δ 65.67.

HRMS (ESI-MS) *m/z*: Calculated $[M-H]^-$ = 273.9741 , Found $[M-H]^-$ = 273.9761

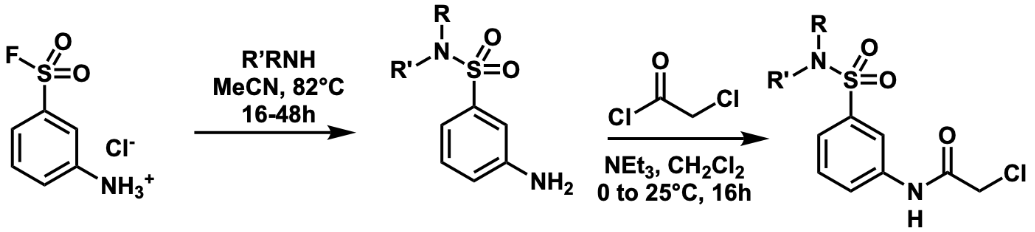


**General procedure for the synthesis of JC19 sulfonamide analogues (NB92, NB177, NB179, and NB001):**

Following a previously published procedure[20], to a µwave vial was added 3-aminobenzenesulfonyl fluoride hydrochloride (1 Eq) and DMAP (0.05 Eq). The vial was capped and purged with argon followed by addition of dry acetonitrile (0.36M), triethylamine (2.1 Eq), and the specified primary or secondary amine (1.05 Eq). The reaction mixture was then left to stir at reflux (82°C) for 16-48h. Upon completion as determined by TLC (30% ethyl acetate in hexanes) the reaction mixture was concentrated down and ran through a silica plug (1:1 ethyl acetate:hexanes). These compounds were verified by LC-MS analysis and used directly for the following step.

The resulting anilines were then subjected to chloroacetylation using the described procedure above for **JC19** with the only deviation being the use of 2.0 Eq of triethylamine.

***N*-(3-(*N*-benzylsulfamoyl)phenyl)-2-chloroacetamide (NB177)**

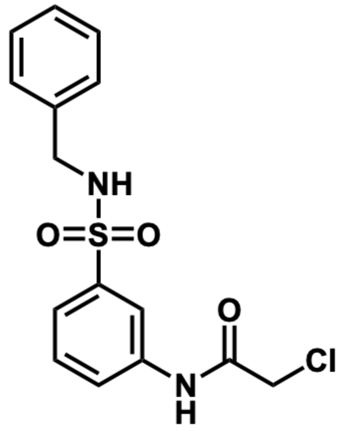

Prepared according to the general procedure with benzylamine and obtained as a yellow oil (7.5mg, 7% over two steps).

$^{1}$H NMR (400 MHz, Chloroform-*d*) δ 8.46 (br, 1H), 7.97 (t, *J* = 2.0 Hz, 1H), 7.89 (ddd, *J* = 8.2, 2.2, 1.0 Hz, 1H), 7.65 (ddd, *J* = 7.9, 1.8, 1.0 Hz, 1H), 7.49 (t, *J* = 8.0 Hz, 1H), 7.31 – 7.15 (m, 5H), 5.00 (t, *J* = 6.1 Hz, 1H), 4.19 (s, 2H), 4.17 (d, *J* = 6.1 Hz, 2H).

$^{13}$C NMR (101 MHz, $CDCl_3$) δ 164.36, 141.13, 137.77, 136.21, 130.20, 128.86, 128.11, 124.11, 123.52, 118.54, 47.53, 42.94.

HRMS (ESI-MS) *m/z*: Calculated $[M-H]^-$ = 337.0414 , Found $[M-H]^-$ = 337.0389

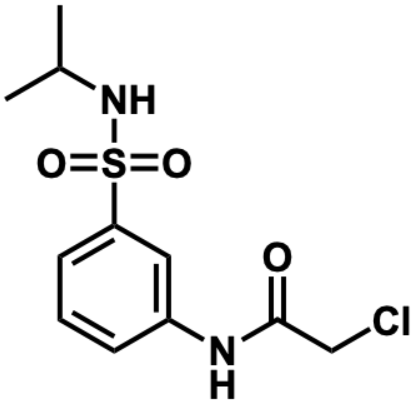

**2-chloro-*N*-(3-(*N*-isopropylsulfamoyl)phenyl)acetamide (NB179)**

Prepared according to the general procedure with isopropylamine and obtained as a yellow oil (11mg, 7% over two steps)

$^{1}$H NMR (400 MHz, Chloroform-*d*) δ 8.39 (br, 1H), 8.04 (t, *J* = 2.0 Hz, 1H), 7.86 (ddd, *J* = 8.1, 2.2, 1.0 Hz, 1H), 7.68 (ddd, *J* = 7.8, 1.8, 1.0 Hz, 1H), 7.51 (t, *J* = 8.0 Hz, 1H), 4.33 (d, *J* = 7.7 Hz, 1H), 4.22 (s, 2H), 3.51 (dp, *J* = 7.6, 6.5 Hz, 1H), 1.11 (d, *J* = 6.5 Hz, 6H).

$^{13}$C NMR (101 MHz, $CDCl_3$) δ 130.19, 123.86, 123.50, 118.43, 46.49, 42.93, 23.99.

HRMS (ESI-MS) *m/z*: Calculated $[M-H]^-$ = 289.0414 , Found $[M-H]^-$ = 289.0524

**N-(3-(*N*-benzyl-*N*-methylsulfamoyl)phenyl)-2-chloroacetamide (NB001)**

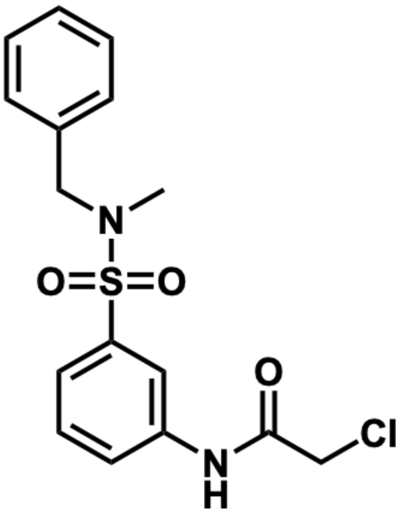

Prepared according to the general procedure with N-methyl-N-benzylamine and obtained as a white solid (55mg, 70% over two steps).

$^{1}$H NMR (400 MHz, Chloroform-*d*) δ 8.58 (br, 1H), 7.97 (t, *J* = 1.9 Hz, 1H), 7.94 (ddd, *J* = 8.0, 2.2, 1.1 Hz, 1H), 7.61 (dt, *J* = 7.9, 1.3 Hz, 1H), 7.53 (t, *J* = 7.9 Hz, 1H), 7.34 – 7.27 (m, 5H), 4.21 (s, 2H), 4.16 (s, 2H), 2.62 (s, 3H).
$^{13}$C NMR (101 MHz, $CDCl_3$) δ 164.59, 138.35, 137.91, 135.48, 130.19, 128.80, 128.47, 128.11, 124.28, 123.74, 118.89, 54.30, 42.98, 34.56.

HRMS (ESI-MS) *m/z*: Calculated $[M-H]^-$ = 351.0570 , Found $[M-H]^-$ = 351.0521

**2-chloro-*N*-(3-(*N*-methylsulfamoyl)phenyl)acetamide (NB92)**

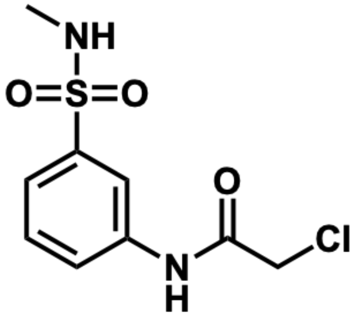

Prepared according to the general procedure with methylamine hydrochloride and 3.1 Eq of triethylamine and obtained as an orange solid (123mg, 40% over two steps).

$^{1}$H NMR (400 MHz, Chloroform-*d*) δ 8.41 (br, 1H), 8.02 (t, *J* = 2.0 Hz, 1H), 7.87 (ddd, *J* = 8.1, 2.2, 1.0 Hz, 1H), 7.66 (ddd, *J* = 7.8, 1.8, 1.0 Hz, 1H), 7.53 (t, *J* = 8.0 Hz, 1H), 4.47 (d, *J* = 5.6 Hz, 1H), 4.22 (s, 2H), 2.70 (d, *J* = 5.4 Hz, 3H).
$^{13}$C NMR (101 MHz, $CDCl_3$) δ 164.35, 140.04, 137.73, 130.22, 124.21, 123.80, 118.68, 42.93, 29.57.

HRMS (ESI-MS) *m/z*: Calculated $[M-H]^-$ = 261.0101 , Found $[M-H]^-$ = 261.0344

#### (E) NMR Spectra

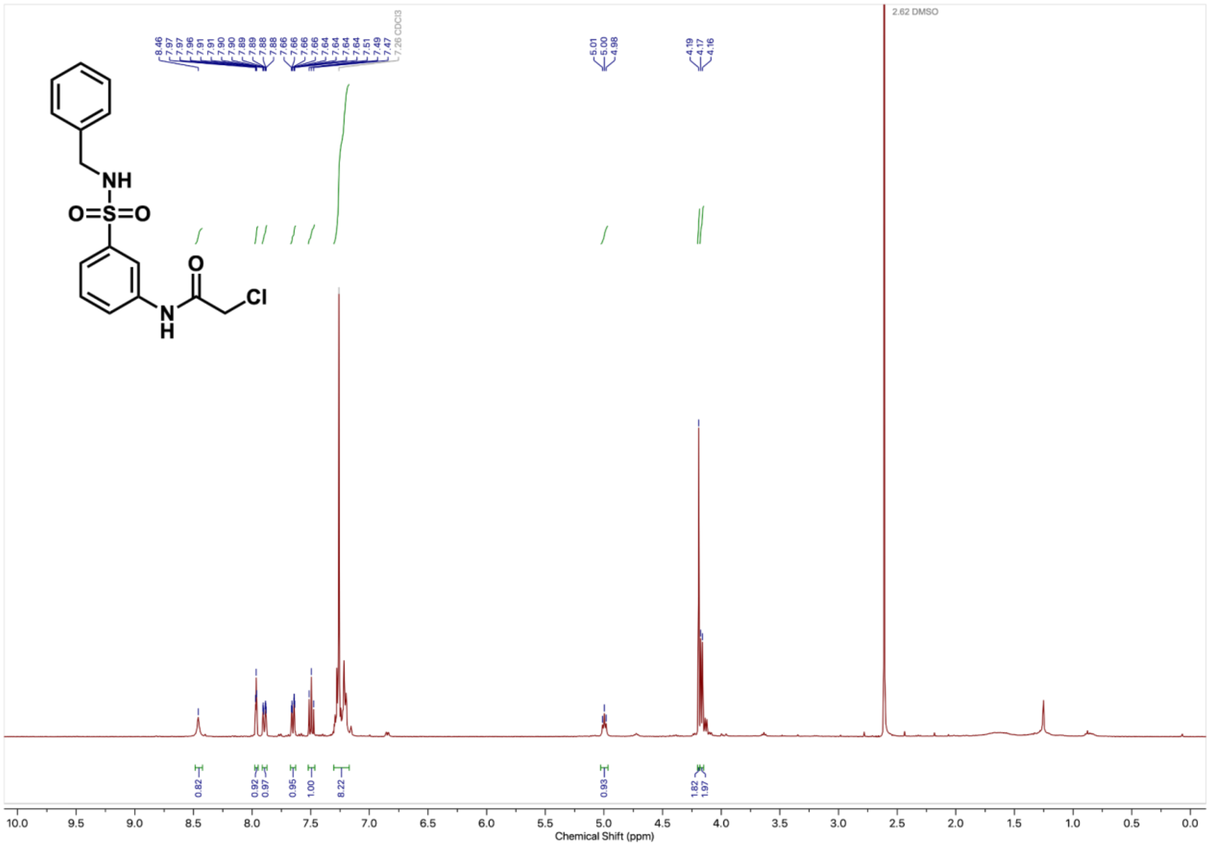

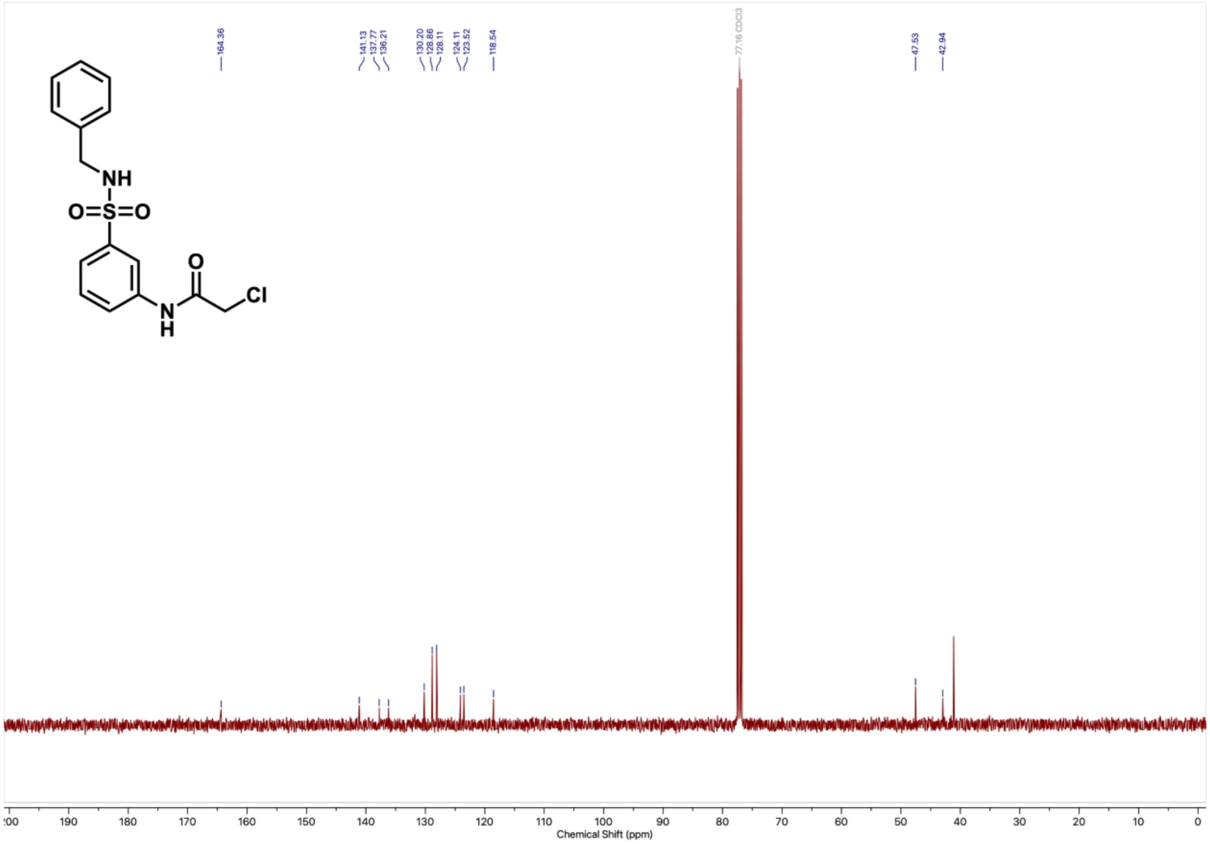
164.36
141.13
137.77
136.21
130.20
128.86
128.11
124.11
123.52
118.54
77.16 CDCl3
47.53
42.94
NH
O=S=O
O
Cl
N
H
200
190
180
170
160
150
140
130
120
110
100
90
80
70
60
50
40
30
20
10
0
Chemical Shift (ppm)

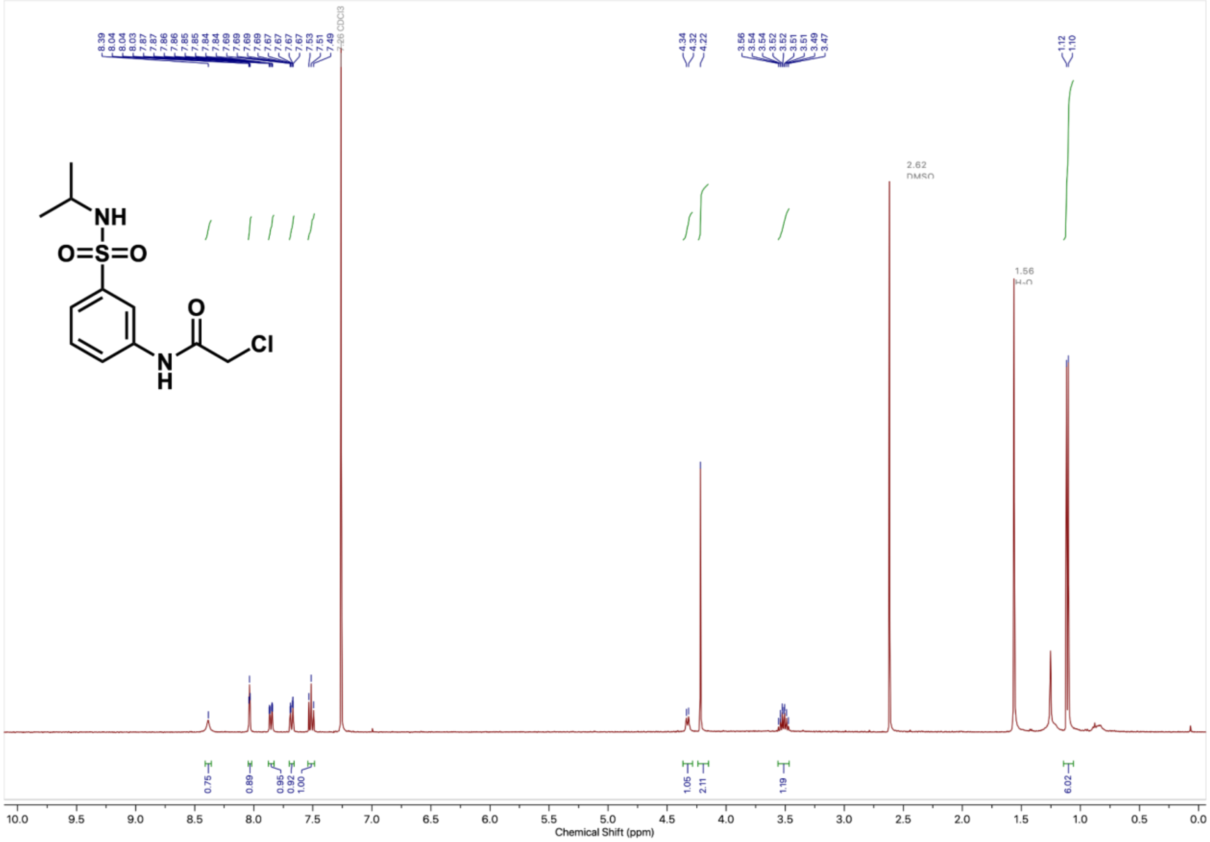

NH
O=S=O
O
N
H
Cl
8.39 8.04 8.04 8.03 7.87 7.87 7.86 7.86 7.85 7.85 7.84 7.84 7.69 7.69 7.69 7.69 7.67 7.67 7.67 7.67 7.53 7.51 7.49
7.26 CDCl3
4.34 4.32 4.22
3.56 3.54 3.54 3.52 3.52 3.51 3.51 3.49 3.47
1.12 1.10
2.62 DMSO
1.56 H2O
0.75 0.89 0.95 0.92 1.00 1.05 2.11 1.19 6.02
10.0 9.5 9.0 8.5 8.0 7.5 7.0 6.5 6.0 5.5 5.0 4.5 4.0 3.5 3.0 2.5 2.0 1.5 1.0 0.5 0.0
Chemical Shift (ppm)

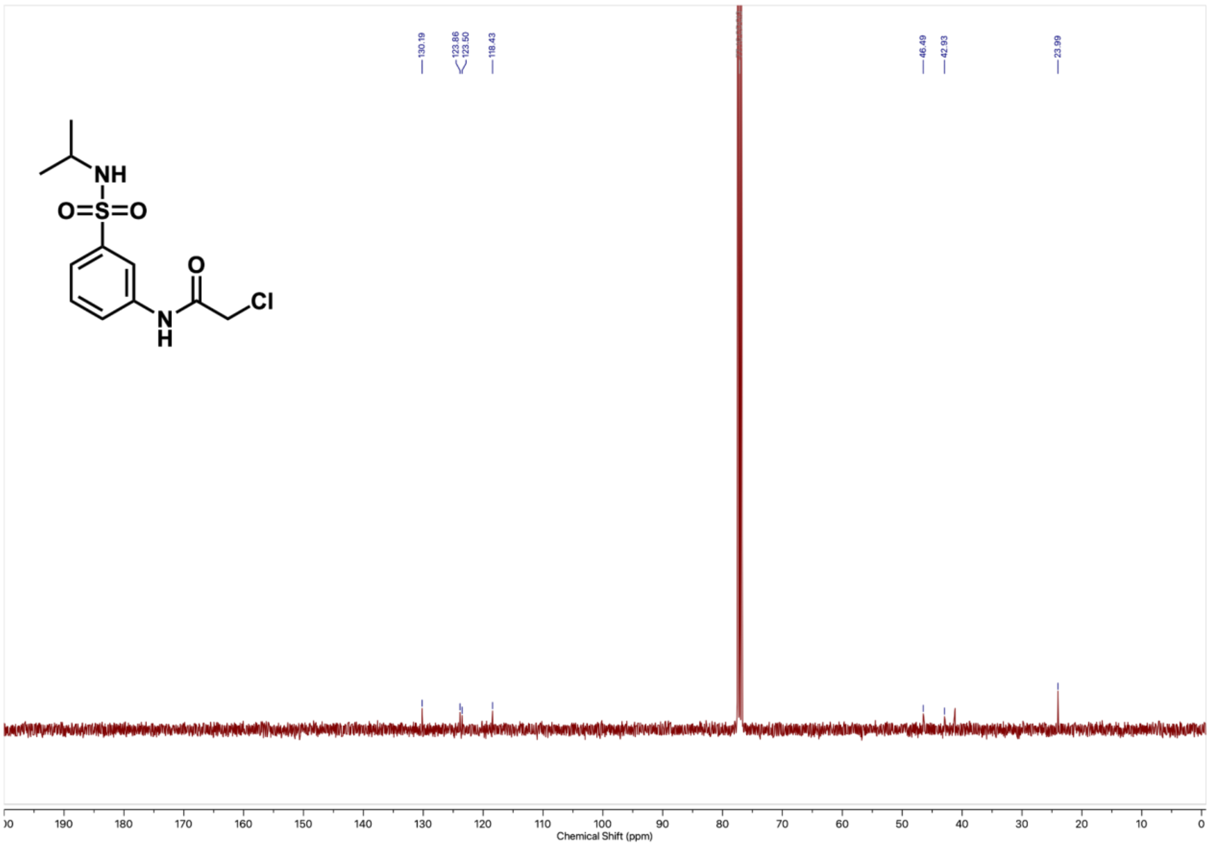

130.19
123.86
123.50
118.43
46.49
42.93
23.99
NH
O=S=O
O
Cl
N
H
190 180 170 160 150 140 130 120 110 100 90 80 70 60 50 40 30 20 10 0
Chemical Shift (ppm)

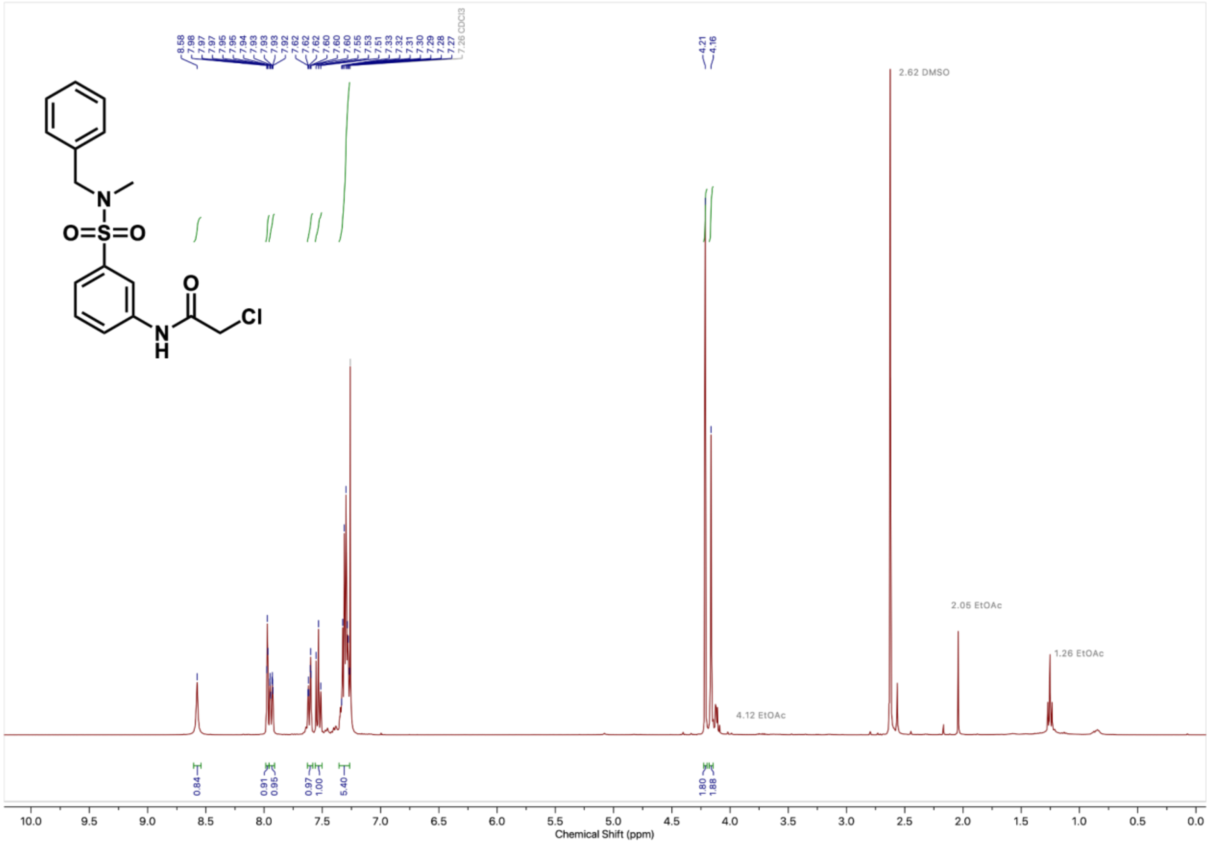
2.62 DMSO
2.05 EtOAc
1.26 EtOAc
4.12 EtOAc
7.26 CDCl3
Chemical Shift (ppm)

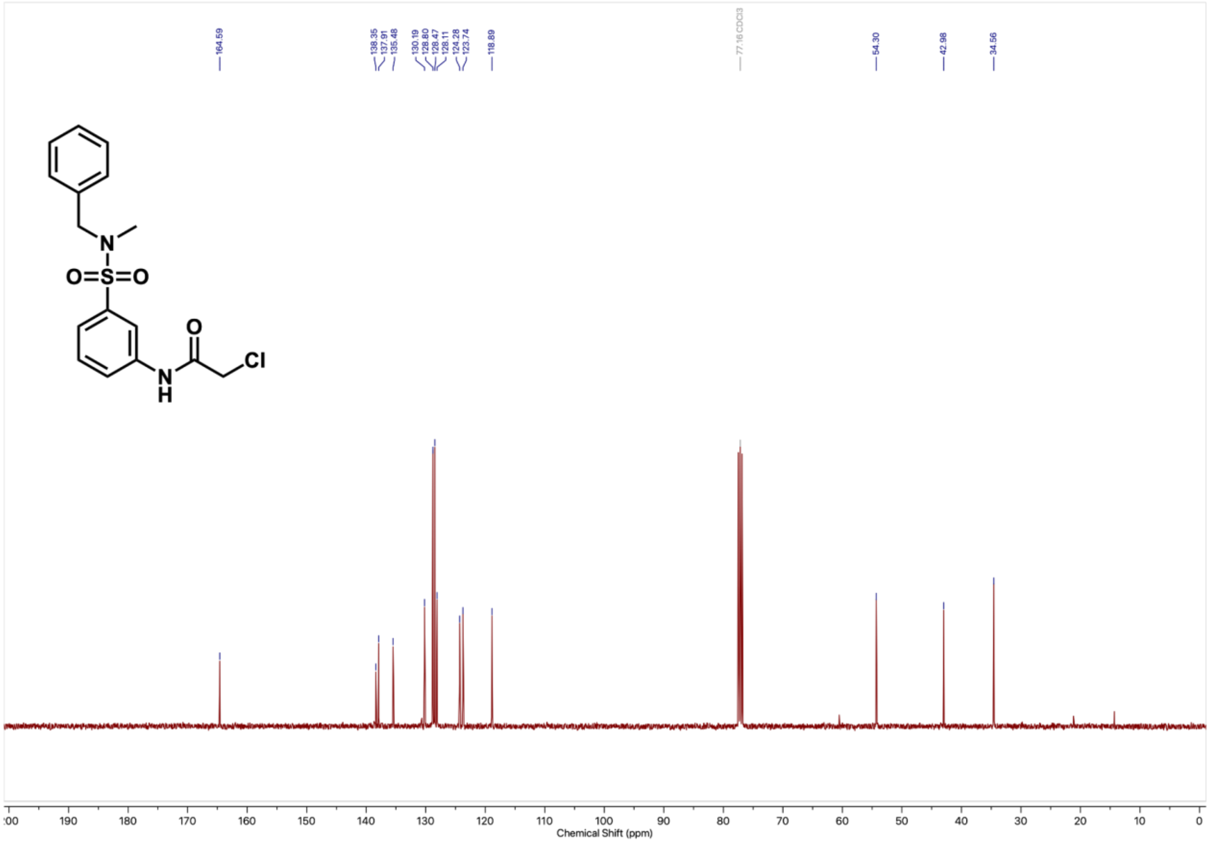
164.59
138.35
137.91
135.48
130.19
128.80
128.47
128.11
124.28
123.74
118.89
77.16 CDCl3
54.30
42.98
34.56
O
Cl
N
H
O=S=O
Chemical Shift (ppm)

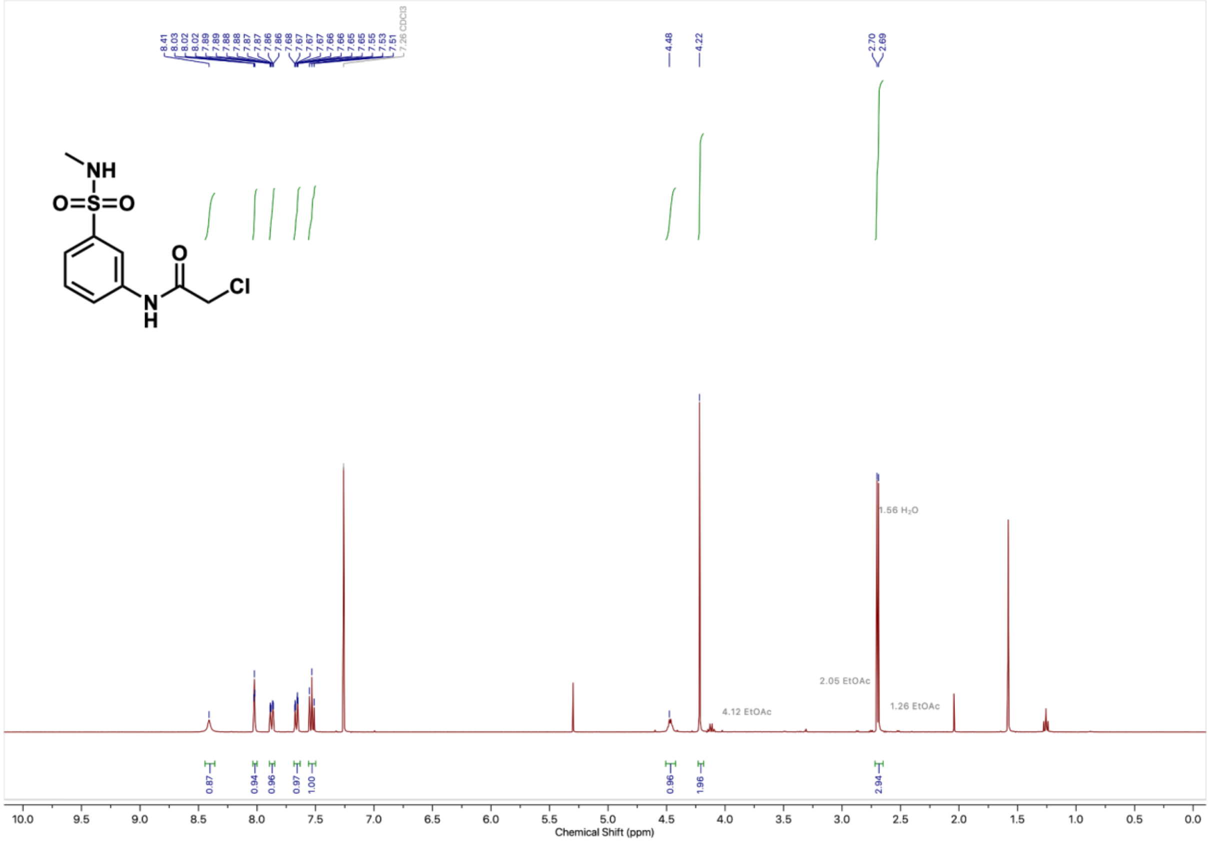
8.41
8.03
8.02
8.02
7.89
7.89
7.88
7.88
7.87
7.87
7.86
7.86
7.68
7.67
7.67
7.67
7.66
7.66
7.65
7.65
7.55
7.53
7.51
7.26 CDCl3
4.48
4.22
2.70
2.69
NH
O=S=O
O
N
H
Cl
1.56 H₂O
2.05 EtOAc
1.26 EtOAc
4.12 EtOAc
0.87
0.94
0.96
0.97
1.00
0.96
1.96
2.94
10.0
9.5
9.0
8.5
8.0
7.5
7.0
6.5
6.0
5.5
5.0
4.5
4.0
3.5
3.0
2.5
2.0
1.5
1.0
0.5
0.0
Chemical Shift (ppm)

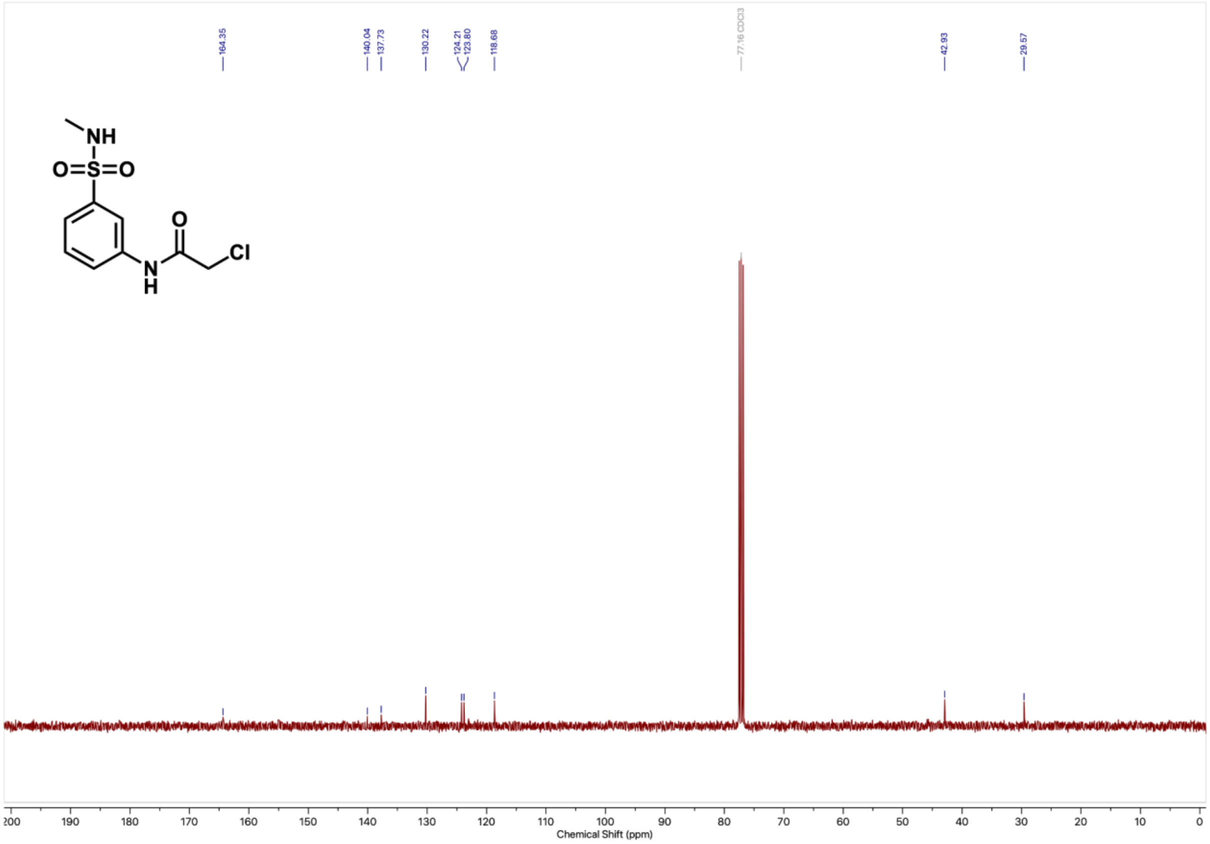

164.35
140.04
137.73
130.22
124.21
123.80
118.68
77.16 CDCl3
42.93
29.57
NH
O=S=O
O
N
H
Cl
200
190
180
170
160
150
140
130
120
110
100
90
80
70
60
50
40
30
20
10
0
Chemical Shift (ppm)

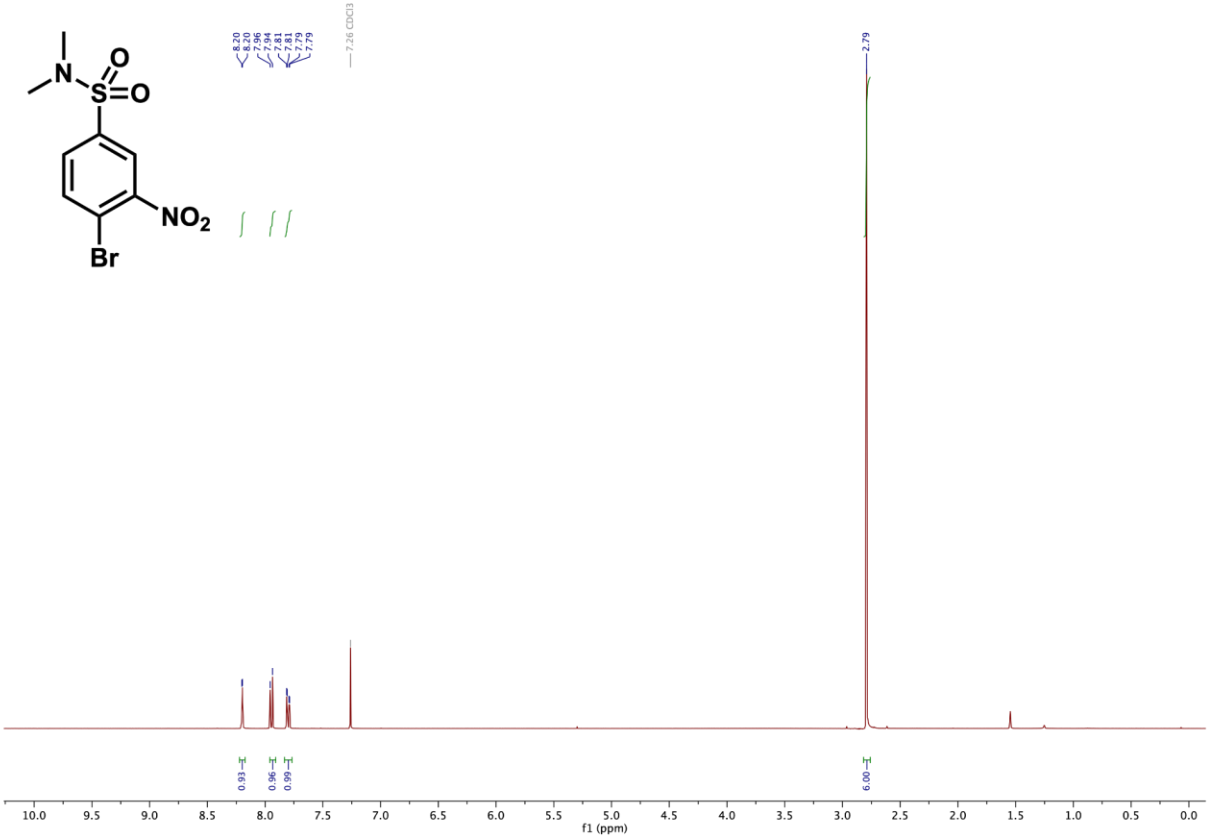
8.20
8.20
7.96
7.94
7.81
7.81
7.79
7.79
7.26 CDCl3
2.79
0.93
0.96
0.99
6.00
f1 (ppm)

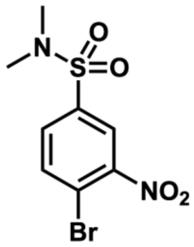
N
O
S
O
NO2
Br

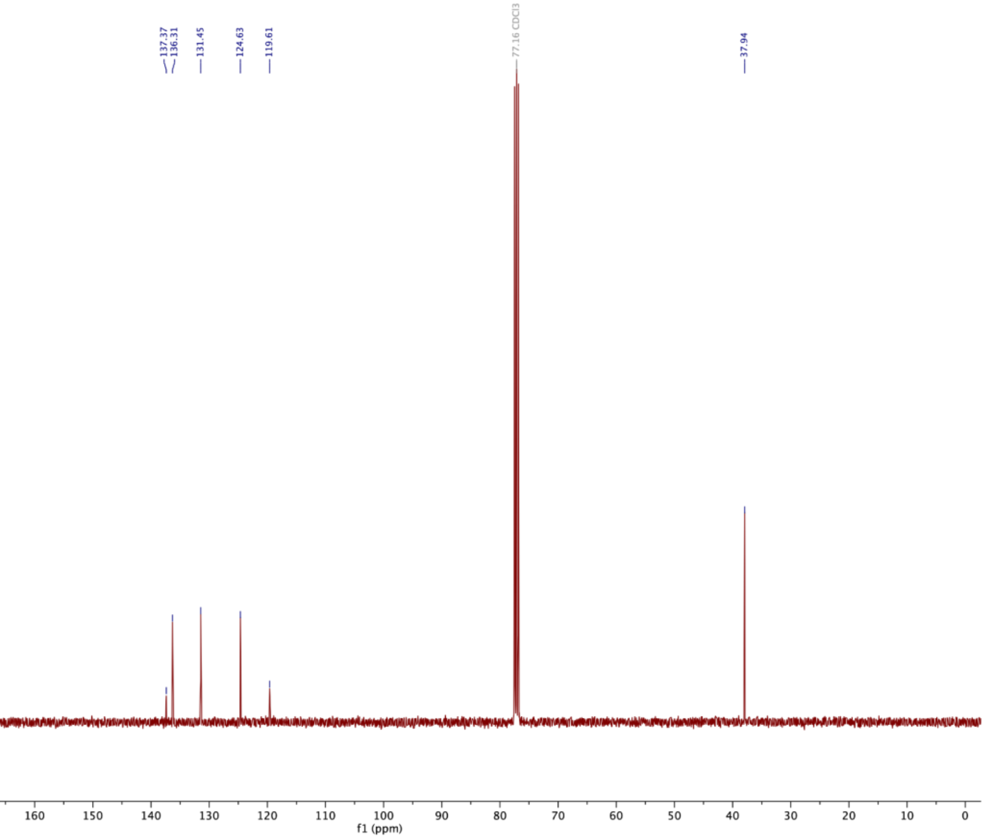
137.37
136.31
131.45
124.63
119.61
77.16 CDCl3
37.94
200 190 180 170 160 150 140 130 120 110 100 90 80 70 60 50 40 30 20 10 0
f1 (ppm)

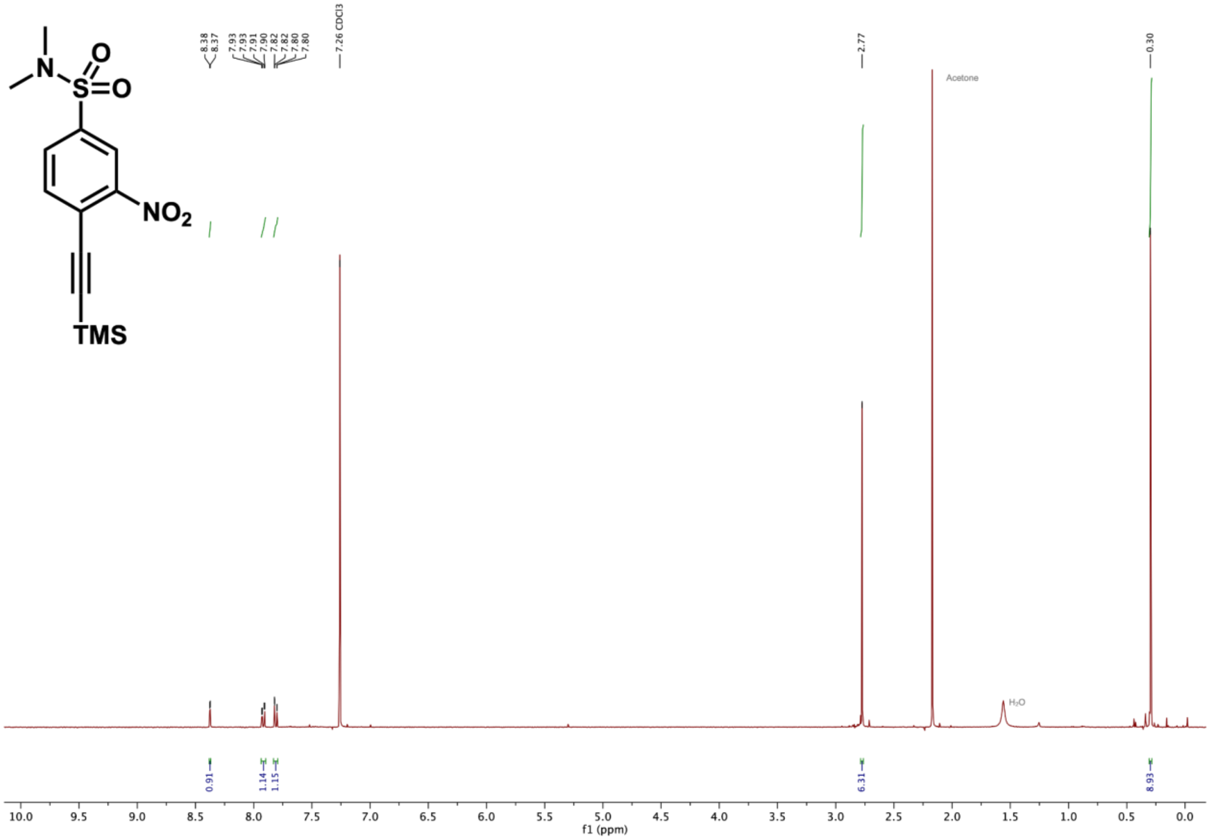

O
N
S
O
NO2
TMS
8.38
8.37
7.93
7.93
7.91
7.90
7.82
7.82
7.80
7.80
7.26 CDCl3
2.77
0.30
Acetone
H2O
0.91
1.14
1.15
6.31
8.93
10.0 9.5 9.0 8.5 8.0 7.5 7.0 6.5 6.0 5.5 5.0 4.5 4.0 3.5 3.0 2.5 2.0 1.5 1.0 0.5 0.0
f1 (ppm)

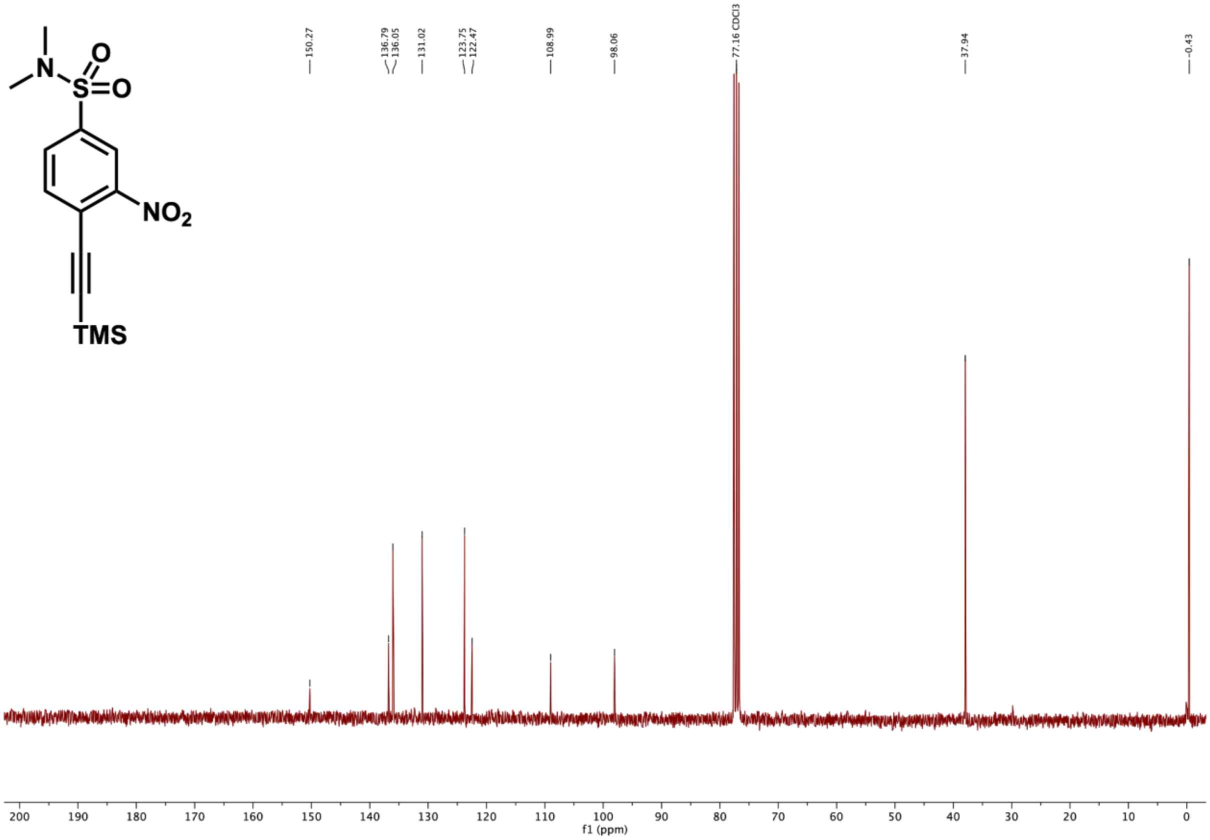
NO2
TMS
O
O
N
S
150.27
136.79
136.05
131.02
123.75
122.47
108.99
98.06
77.16 CDCl3
37.94
-0.43
200 190 180 170 160 150 140 130 120 110 100 90 80 70 60 50 40 30 20 10 0
f1 (ppm)

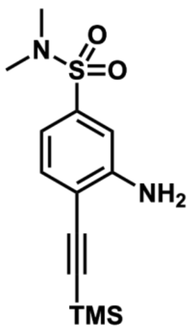

O
N
S
O
NH2
TMS


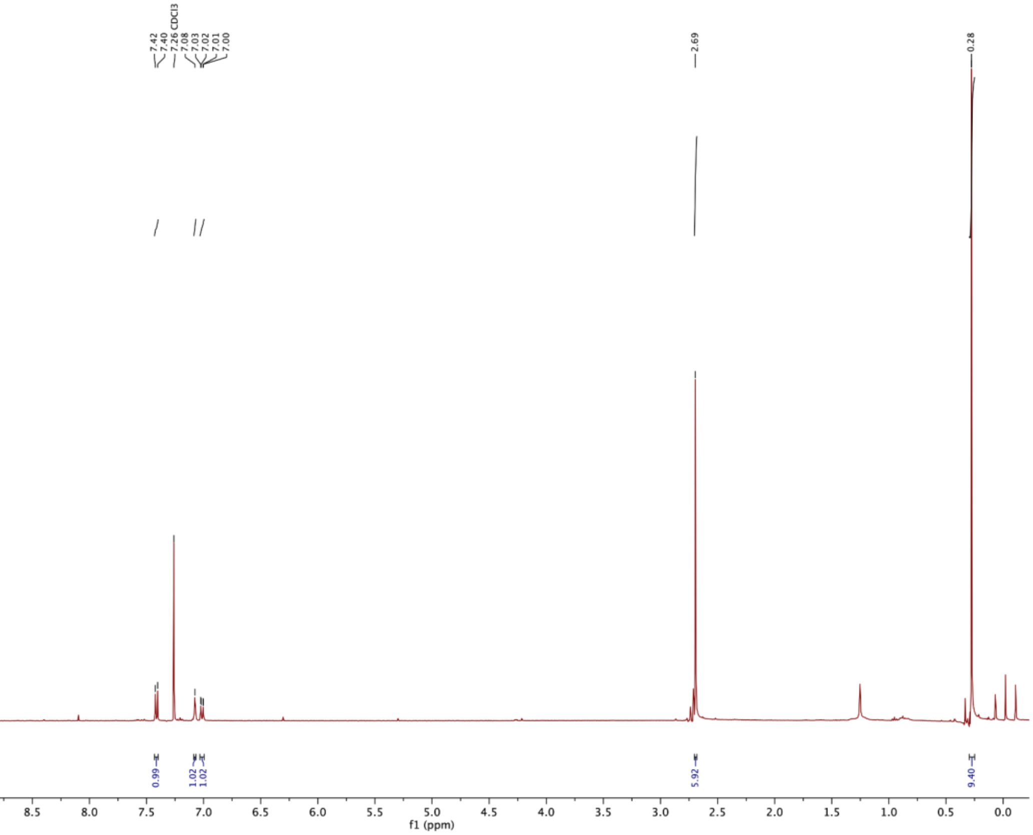

7.42
7.40
7.26 CDCl3
7.08
7.03
7.02
7.01
7.00
2.69
0.28
0.99
1.02
1.02
5.92
9.40
10.0
9.5
9.0
8.5
8.0
7.5
7.0
6.5
6.0
5.5
5.0
4.5
4.0
3.5
3.0
2.5
2.0
1.5
1.0
0.5
0.0
f1 (ppm)

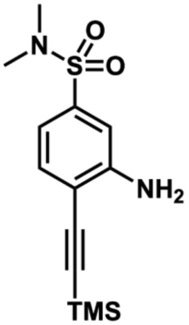

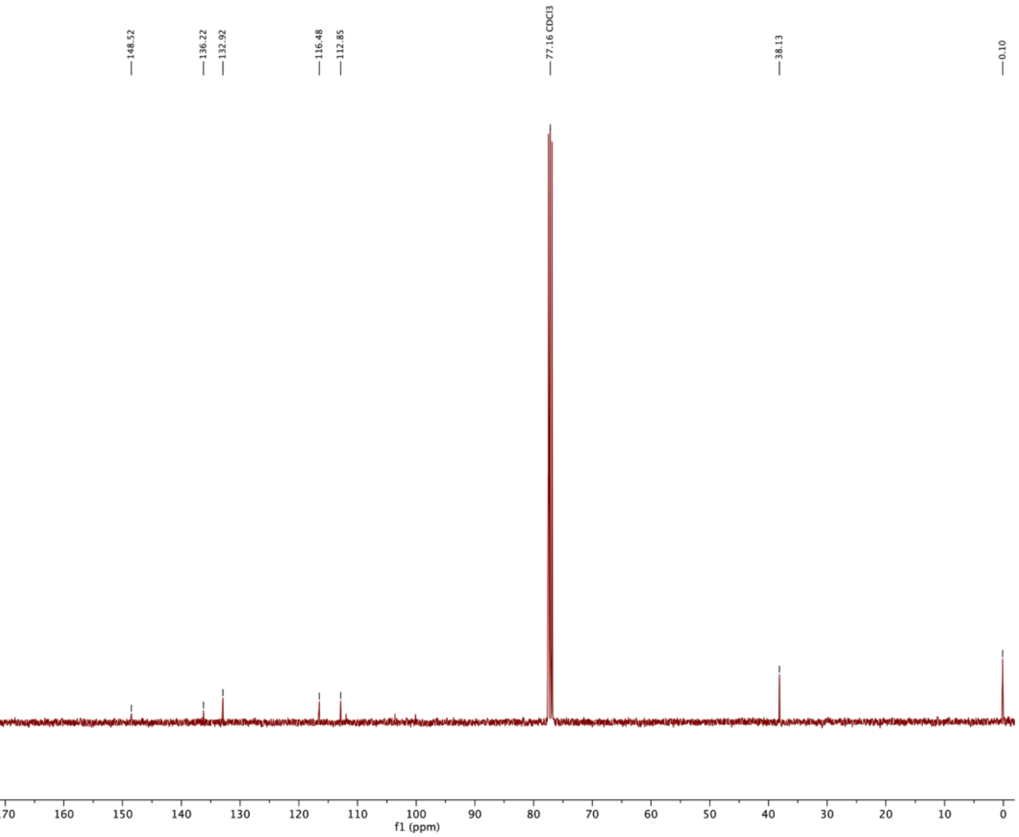

148.52
136.22
132.92
116.48
112.85
77.16 CDCl3
38.13
0.10
200
190
180
170
160
150
140
130
120
110
100
90
80
70
60
50
40
30
20
10
0
f1 (ppm)

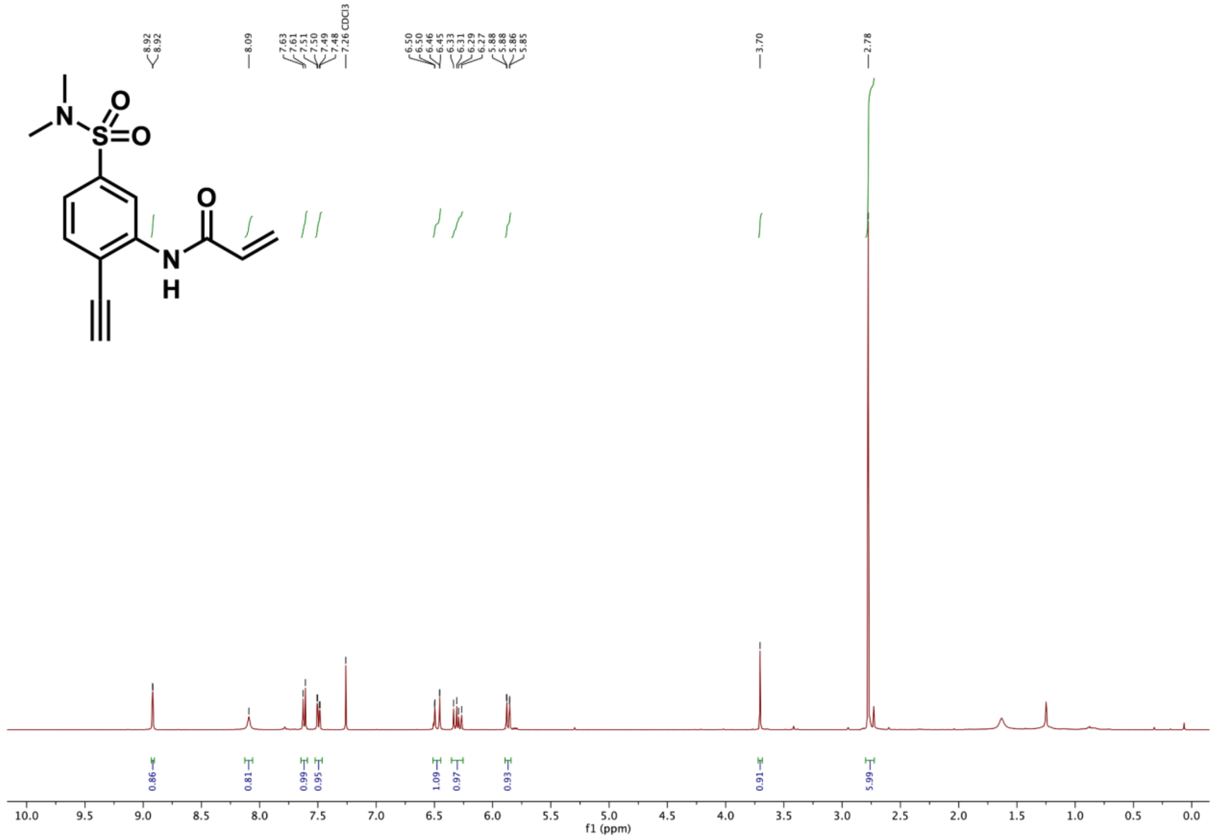

8.92
8.92
8.09
7.63
7.61
7.51
7.50
7.49
7.48
7.26 CDCl3
6.50
6.50
6.46
6.45
6.33
6.31
6.29
6.27
5.88
5.88
5.86
5.85
3.70
2.78
0.86
0.81
0.99
0.95
1.09
0.97
0.93
0.91
5.99
f1 (ppm)

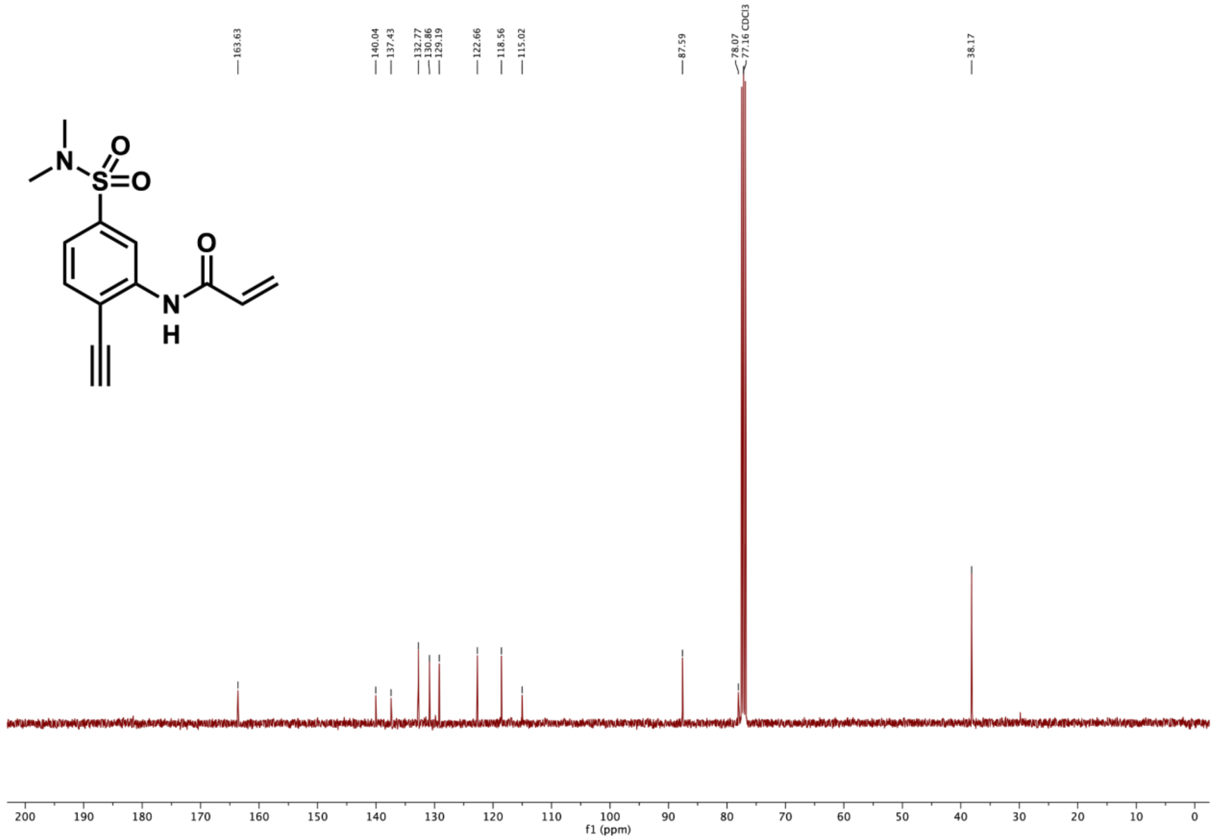
163.63
140.04
137.43
132.77
130.86
129.19
122.66
118.56
115.02
87.59
78.07
77.16 CDCl3
38.17
O
N
S
O
O
N
H
200 190 180 170 160 150 140 130 120 110 100 90 80 70 60 50 40 30 20 10 0
f1 (ppm)

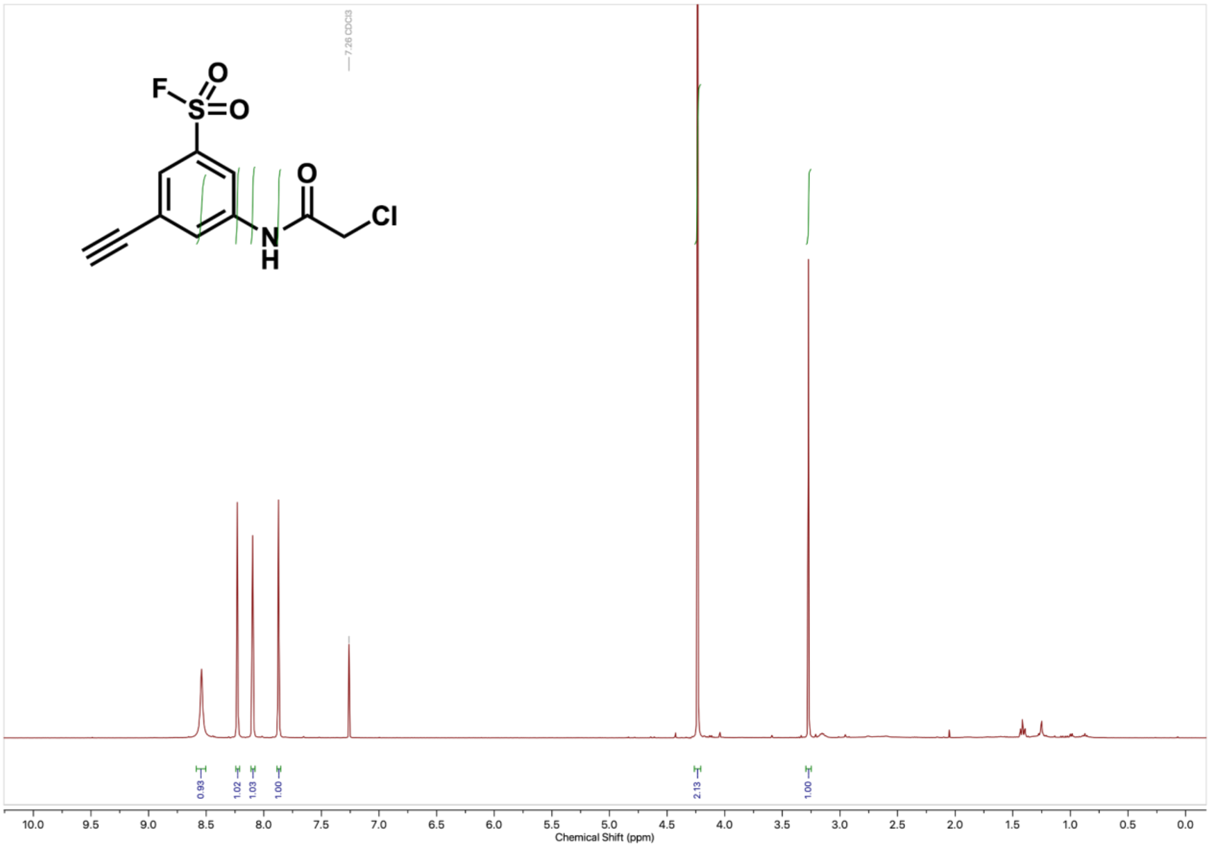

7.26 CDCl3
0.93
1.02
1.03
1.00
2.13
1.00
10.0
9.5
9.0
8.5
8.0
7.5
7.0
6.5
6.0
5.5
5.0
4.5
4.0
3.5
3.0
2.5
2.0
1.5
1.0
0.5
0.0
Chemical Shift (ppm)

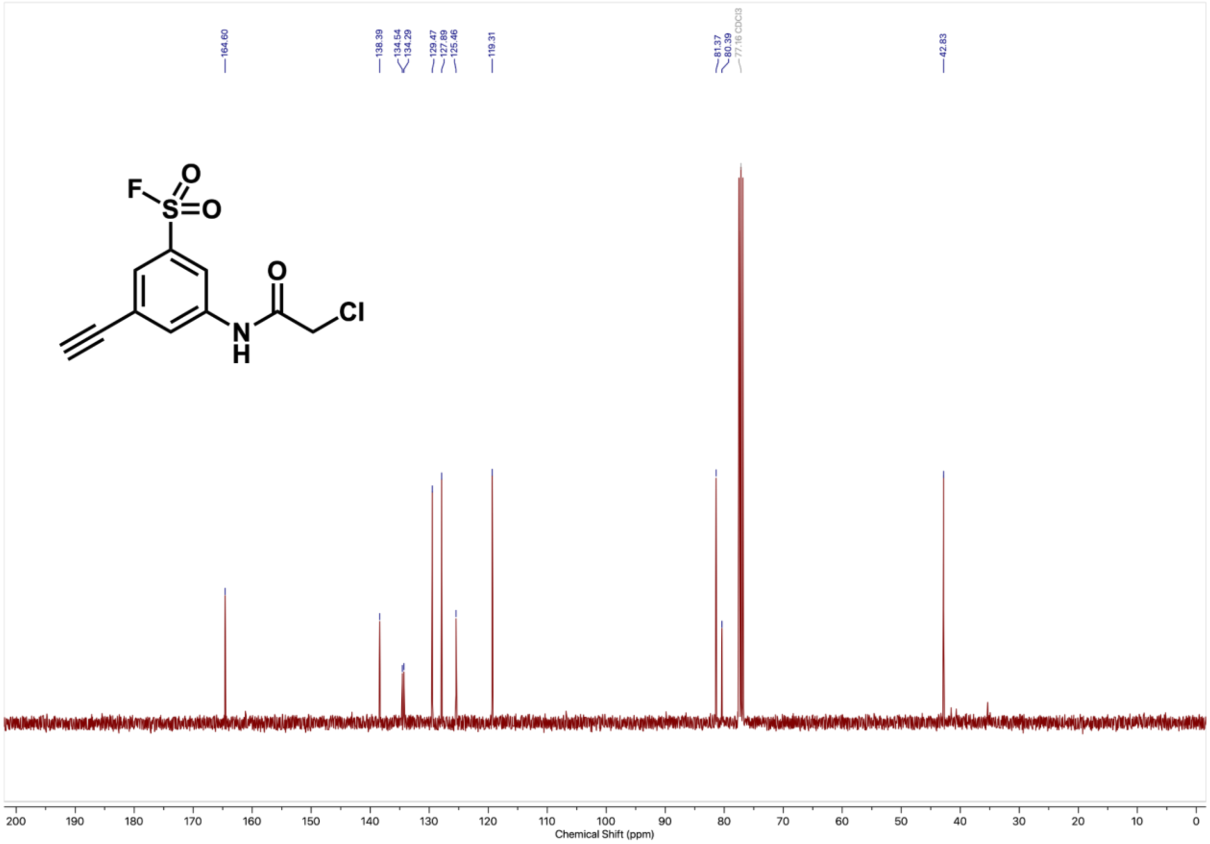

164.60
138.39
134.54
134.29
129.47
127.89
125.46
119.31
81.37
80.39
77.16 CDCl3
42.83
F
S
O
O
O
N
H
Cl
200
190
180
170
160
150
140
130
120
110
100
90
80
70
60
50
40
30
20
10
0
Chemical Shift (ppm)

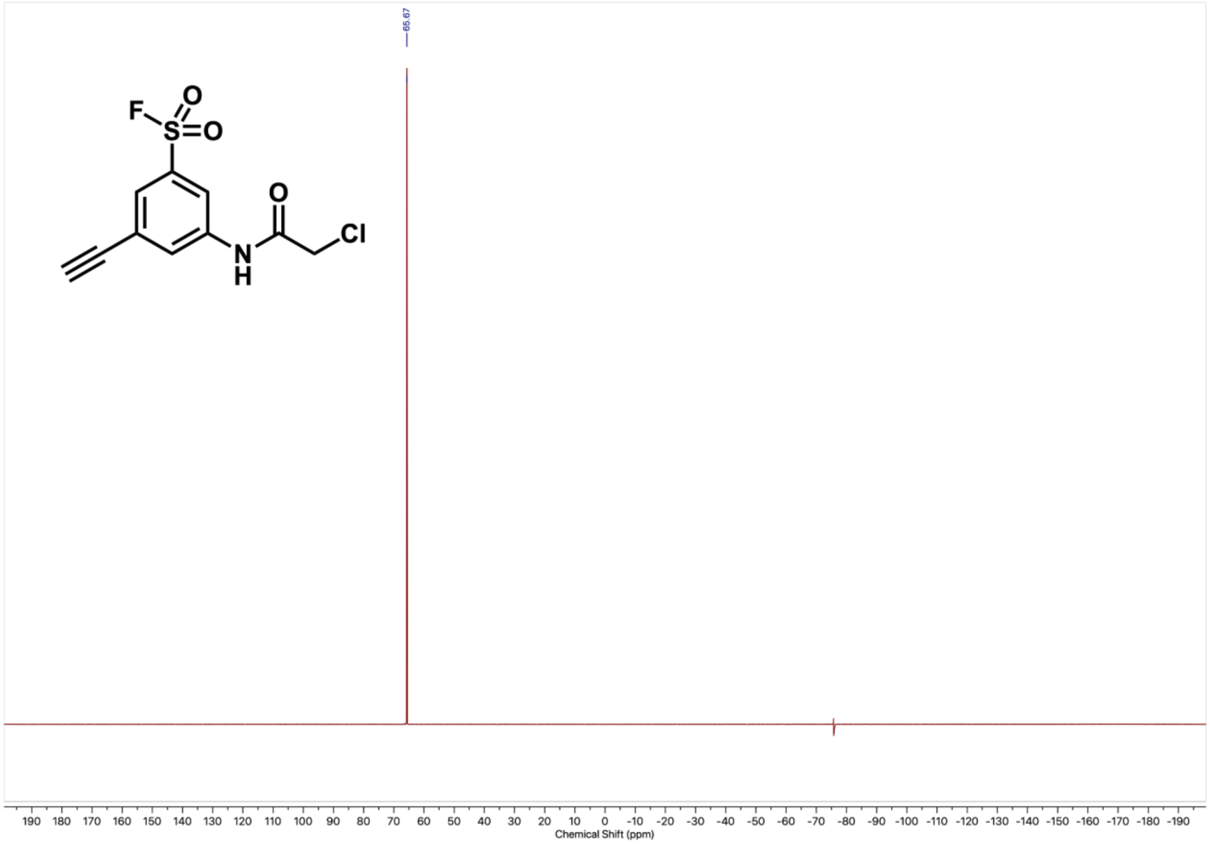

65.67
F
S
O
O
O
N
H
Cl
190 180 170 160 150 140 130 120 110 100 90 80 70 60 50 40 30 20 10 0 -10 -20 -30 -40 -50 -60 -70 -80 -90 -100 -110 -120 -130 -140 -150 -160 -170 -180 -190
Chemical Shift (ppm)

#### (F) References


1. Imprachim, N., Yosaatmadja, Y. & Newman, J. A. Crystal structures and fragment screening of SARS-CoV-2 NSP14 reveal details of exoribonuclease activation and mRNA capping and provide starting points for antiviral drug development. *Nucleic Acids Res.* **51**, 475–487 (2023).
2. Schneider, C. A., Rasband, W. S. & Eliceiri, K. W. NIH Image to ImageJ: 25 years of image analysis. *Nat. Methods* **9**, 671–675 (2012).
3. Kim, D.-K. *et al.* A Comprehensive, Flexible Collection of SARS-CoV-2 Coding Regions. *G3 (Bethesda)* **10**, 3399–3402 (2020).
4. Branon, T. C. *et al.* Efficient proximity labeling in living cells and organisms with TurboID. *Nat. Biotechnol.* **36**, 880–887 (2018).
5. Campeau, E. *et al.* A versatile viral system for expression and depletion of proteins in mammalian cells. *PLoS ONE* **4**, e6529 (2009).
6. Liu, H. & Naismith, J. H. An efficient one-step site-directed deletion, insertion, single and multiple-site plasmid mutagenesis protocol. *BMC Biotechnol.* **8**, 91 (2008).
7. Weerapana, E. *et al.* Quantitative reactivity profiling predicts functional cysteines in proteomes. *Nature* **468**, 790–795 (2010).
8. Chen, E. Y. *et al.* Enrichr: interactive and collaborative HTML5 gene list enrichment analysis tool. *BMC Bioinformatics* **14**, 128 (2013).
9. Kuleshov, M. V. *et al.* Enrichr: a comprehensive gene set enrichment analysis web server 2016 update. *Nucleic Acids Res.* **44**, W90-7 (2016).
10. Xie, Z. *et al.* Gene Set Knowledge Discovery with Enrichr. *Curr. Protoc.* **1**, e90 (2021).
11. UniProt Consortium. Uniprot: the universal protein knowledgebase in 2023. *Nucleic Acids*

*Res.* **51**, D523–D531 (2023).
